# Lipid Flipping, O-antigen Shielding, and Water Dynamics Revealed by 100-200 µs Coarse-Grained Simulations of *E. coli* Outer Membrane Proteins

**DOI:** 10.1101/2024.12.09.627556

**Authors:** Andrew Kalenkiewicz, Adrian H. Elcock

## Abstract

Gram-negative (GN) bacterial infections are a major global health concern, with antibiotic-resistant strains representing a growing problem. The outer membrane (OM) of GN bacteria is the primary interface between the cell and its environment, and it plays a crucial role in regulating how GN bacteria interact with a host’s immune system and with antimicrobial drugs. Consequently, understanding the structural details of the GN OM is an essential step toward addressing the pressing threat of GN organisms. One way to gain insights into complex molecular structures like the GN OM is to use coarse-grained molecular dynamics (MD) simulations. In this work, we use the *Martini3* coarse-grained force field to model and simulate ten different OM proteins (OMPs) from *E. coli* embedded in four different OM models that each contain lipopolysaccharides (LPS) with different O-antigen lengths. We extensively assess OMP-phospholipid interactions and demonstrate that TolC, in particular, shows a strong preference for binding cardiolipin. We also uncover a phenomenon that, to our knowledge, has not been previously described in the simulation literature—i.e., the ability of OMPs to serve as sites of phospholipid flipping from the OM inner leaflet to the outer leaflet. Finally, we show that longer LPS O-antigen chains occlude OMPs, and we provide a detailed characterization of water and ion movement throughout our simulations.

## Introduction

The outer membrane (OM) of Gram-negative (GN) bacteria is the primary interface between the cell and its environment. Essential functions such as regulation of nutrient transport, defense against host immune system components, and resistance to antimicrobial agents are all dependent on the OM and its integrity [1]. Given the cost and complexity of modeling even relatively simple membrane bilayer systems, investigations of the protein-membrane systems that utilize molecular simulations often employ coarse-grained models. For instance, Corey, et al. [2] recently described an extensive study of 42 different *E. coli* inner membrane proteins using molecular dynamics simulations performedwith the *Martini3* [3] forcefield. They identified a number of distinct patterns involving interactions of the proteins with the different phospholipids comprising the inner membrane. Another study spearheaded by Shearer, et al. [4] examined six *E. coli* outer membrane proteins and determined that they also exhibited unique interaction patterns with the surrounding outer membrane. In separate work, Vaiwala and Ayappa [5] described the construction of a coarse-grained model of lipopolysaccharide (LPS) using *Martini3* parameters, and showed that the resulting model produced results consistent with those obtained from all-atom simulations across a wide variety of descriptive parameters. As one final example, Du et al. [6] ran *Martini2* coarse-grained simulations of AcrB in a model of the *E. coli* inner membrane. They found that there was an enrichment of cardiolipin and phosphatidylglycerol in the vicinity of AcrB, which was enhanced by the binding of AcrZ (a protein known to regulate the AcrA-AcrB-TolC complex).

There is a wealth of additional evidence that protein-lipid interactions in *E. coli* play a crucial role in modulating protein functionality. Corey, et al. [7] showed that the activity of SecYEG—which transports proteins across the *E. coli* inner membrane—is influenced by the binding of cardiolipin which appears to drive dimerization of SecYEG and increases its ATPase activity. The same authors employed coarse-grained molecular dynamics simulations to identify cardiolipin binding sites on SecYEG and validated those sites *in vitro* using native mass spectrometry [7]. Laganowsky, et al. [8] used ion mobility mass spectrometry to investigate the way that the *E. coli* aquaporin AqpZ and the ammonia channel AmtB are influenced by lipid binding. They showed that AqpZ water transport activity in reconstituted liposomes is inhibited in the absence of cardiolipin, and that AmtB selectively binds and is stabilized by phosphatidylglycerol. They subsequently determined x-ray crystallographic structures of AmtB in the presence and absence of lipid, demonstrating that there were distinct conformational changes consistent with the formation of close contacts between amino acid residues and phosphatidylglycerol. As a final example, Gupta et al. [9] conducted a study of membrane protein oligomerization and found that the bacterial leucine transporter LeuT depends on cardiolipin to achieve its dimeric state. They utilized molecular dynamics simulations to determine that cardiolipin acts as a bridge, with its bi-phosphate head group interacting with basic residues on each of the two LeuT monomers.

The importance of protein-lipid interactions in general bilayer physiology has also been highlighted by studies such as that conducted by Corradi, et al. [10], who simulated coarse-grained models of more than 60 different lipid types and quantified interactions between each lipid and a set of ten different membrane protein types. They determined that there was an enrichment of specific lipid types in the “shells” directly surrounding the various membrane proteins they examined, along with a depletion of other lipid types. The membrane compositions were modeled after eukaryotic systems, which are much more complex and multifaceted than those of *E. coli*, but their results demonstrate the principle that protein-lipid interactions can be highly specific and consequently play a role in guiding the structure and organization of membrane systems. Scott et al. [11] conducted coarse-grained simulations of membrane protein-bilayer self-assembly and compared the results to experimentally determined structures, showing that their simulations faithfully reproduced the relative positions of proteins within bilayer systems. Their work provides further strong evidence that coarse-grained models of membrane protein systems are a useful tool for making realistic predictions concerning lipid bilayer function and physiology.

This paper describes our use of *Martini3* [3]—a highly popular force field used in coarse-grained molecular dynamics studies—to simulate and characterize the multifold interactions among the lipids, lipopolysaccharides, and the top ten most highly expressed OMPs within the outer membrane of *E. coli*. One goal of our work is to examine the intermolecular interactions between OMPs and the three component phospholipids that comprise the inner leaflet of the *E. coli* outer membrane, i.e. phosphatidylethanolamine, phosphatidylglycerol, and cardiolipin. As previously mentioned, Corey, et al. [2] have already shown that phospholipids can engage in specific interactions with discrete regions of *E. coli* inner membrane proteins, and there is a broad array of experimental evidence suggesting that such interactions heavily influence the folding, localization, and functionality of inner membrane proteins [7, 9]. However, there is a relative deficit of information characterizing interactions between phospholipids and OM proteins, and our work aims to fill this gap by quantifying the OMP-phospholipid interactions that occur during the molecular dynamics trajectories.

During our analysis of OMP-phospholipid interactions, we encountered an unexpected phenomenon: lipid “flipping.” In the vast majority of our 40 different simulations, we saw phospholipids flip from the lower leaflet of the OM to the upper leaflet (and sometimes back to the lower leaflet). Moreover, these flipping events occurred exclusively in close proximity to the protein. Visual analysis of the simulation trajectories showed that a common feature of these flipping events was the stabilization of the lipid headgroup by the protein backbone beads. We note that maintenance of OM leaflet asymmetry is critically important for GN cell integrity, and the presence of phospholipids in the OM has been established as a sign of cell stress associated with a reduction in the OM’s barrier functionality [12]. We carried out a detailed analysis of these flipping events, characterizing their frequency, spatial proximity to proteins, and duration. Our results shed light on a heretofore undescribed phenomenon that may play an important role in OM homeostasis for GN bacteria.

One important detail of this study is that our OMP simulations utilize a range of O-antigen (O-antigen) chain lengths for the all-important lipopolysaccharides of the OM, ultimately modeling longer chains than have been used in previous simulation studies. We note that experimental evidence from *in vivo* animal studies indicates that the expression of longer O-antigen chains can facilitate “shielding” of surface antigens from host immune factors such as antibodies [13]. Other studies have shown that regulation of O-antigen chain length plays a crucial role in protection against complement-mediated cell lysis [14]. We have modeled O-antigen chains with up to 10 O-antigen repeats, setting an upper limit that better reflects the OM architecture of pathogenically relevant *E. coli* strains [15]. Our approach has allowed us to investigate the role that O-antigen chain length plays in controlling the surface accessibility of *E. coli* OMPs, as well as the flow of water through the OMP porins OmpC and OmpF and the efflux pump TolC. We also provide a more general description of the ways in which water and ions traverse our bilayer systems, focusing on such metrics as protein proximity and the duration of traversal events. Altogether, therefore, the simulations described here provide a number of new insights into the ways that proteins, phospholipids, and LPS combine to produce the OM’s critical barrier functionality.

## Methods

### System Generation

We compiled a list of the ten most highly expressed *E. coli* OM proteins (out of 86 total) using data in the literature from researchers who used ribosome profiling to calculate absolute synthesis rates and copy numbers for the entire *E. coli* genome [16]. For rich media growth conditions, the top ten most highly expressed proteins in the OM (in decreasing order of abundance) are: OmpA, OmpC, OmpX, OmpF, OmpT, MipA, Tsx, TolC, FadL, and YdiY (Table S1, Figure 1). To obtain full-length structures of each protein, we utilized the AlphaFold Protein Structure Database [17, 18], except in the case of the three trimeric proteins in the list, i.e. TolC, OmpC, and OmpF. For the latter three proteins, we used a local implementation of AF2-Complex [19] to generate structures of the trimeric forms. Each protein was oriented for membrane insertion using the PPM (Positioning of Proteins in Membranes) server [20]. For the task of embedding each oriented protein in a patch of the *E. coli* outer membrane, we employed CHARMM-GUI’s “Bilayer Builder” utility [21, 22]. For each protein, we generated four separate all-atom bilayer systems that differed only in the number of O-antigen repeats specified for the outer leaflet LPS component. The lipids for the lower leaflet were constant across all our systems, and the ratios we chose were those recommended by Pogozheva et al. [23] to model the Gram-negative bacterial OM—i.e. phosphatidylethanolamine (PPPE), phosphatidylglycerol (PVPG), and cardiolipin (PVCL2), in copy number ratios of 75:20:5. We specified that the LPS should have an R1 core and O6 antigen repeats of either length 10, 5, 2, or 0 for the four systems:we chose the R1 core and O6 antigen because these have previously been used successfully in implementing a coarse-grained model of the *E. coli* OM [5]. The ratio for LPS with respect to the lower leaflet was 35:100 (i.e. 35 LPS molecules for every 100 lower leaflet lipids in the aforementioned 75:20:5 ratio, again as recommended by Pogozheva, et al. [23]). We chose initial box dimensions in the xy-plane as 125 × 125 Å for the seven monomer proteins and 150 × 150 Å for the three homotrimer proteins (which have a relatively larger circumference than the monomers), with a solvent buffer of 25 Å above and below the bilayer in the z-dimension. After the lipid system was packed by CHARMM-GUI, we neutralized the negatively charged R1 core of the LPS with Ca^2+^ counterions and gave the system a NaCl concentration of 0.15 M, with CHARMM-GUI automatically implementing a slight surplus of Na^+^ cations to neutralize the negative charge stemming from the protein.

**Figure 1:**
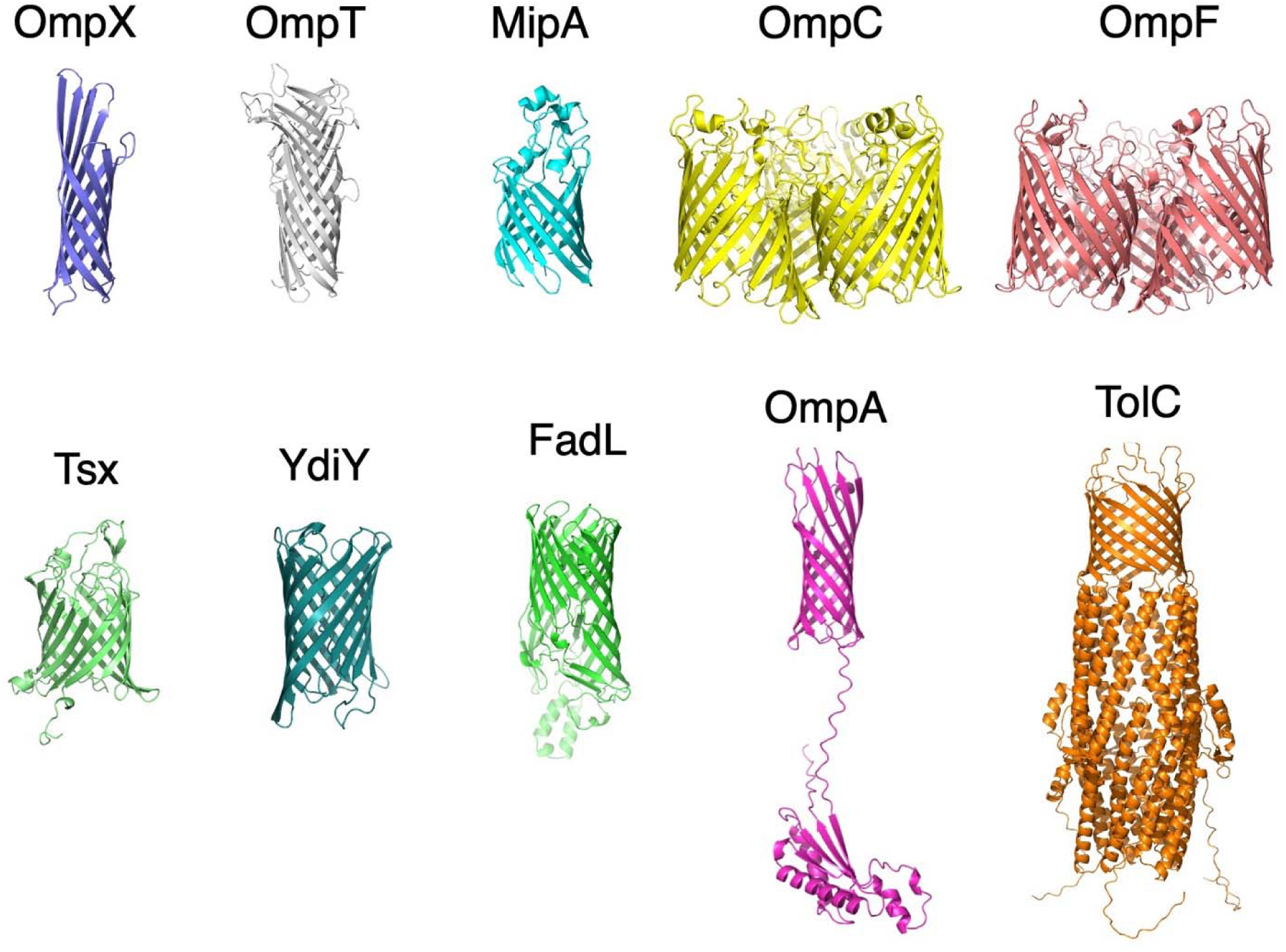
We chose to center this study on the 10 most highly expressed OM proteins, pictured here. All 10 are β-barrel proteins, but OmpC and OmpF are homotrimers in their native state. TolC is also homotrimeric, but in this case the polypeptide chains combine to form a single β-barrel rather than a trimer of 3 β-barrels as with OmpC and OmpF. OmpA is notable in that it contains a periplasmic domain that is linked to the OM-embedded β-barrel by a disordered loop. In vivo, this periplasmic domain interacts non-covalently with the peptidoglycan component of the cell envelope.

### Coarse-grained parameterization

Upon completing the last two CHARMM-GUI stages of solvation and all-atom MD-minimization, we downloaded the output all-atom structures from the CHARMM-GUI server and implemented a custom, in-house *Martini3* coarse-graining strategy (see Supporting Information, pp. 3-6). This was necessary in order to implement our desired *Martini3* lipid and LPS models. Although CHARMM-GUI is capable of building coarse-grained bilayer systems for the *Martini2* force field, at the time this work was conducted, the *Martini3* lipid libraries had not yet been implemented. *Martini3* is an updated, re-parameterized forcefield intended as a replacement for *Martini2* [3], and one of its key enhancements is the design of superior parameters for carbohydrate interactions. Since our systems include LPS (which has many carbohydrate moieties), we deemed it critical to utilize *Martini3* despite the additional required steps that this entailed. Our custom coarse-graining scheme for phospholipids (Figure S2) was based heavily on *SwarmCG*’s models [24], while our initial LPS model was borrowed from Vaiwala and Ayappa [5]. Our proteins were coarse-grained with *Martinize2* [25]. Further details of the described course-graining protocols can be found in the Supporting Information.

### Molecular Dynamics Simulations

We used the complete *Martini3* systems as input for molecular dynamics (MD) simulations performed with GROMACS [26]. The MD protocol was implemented as follows: we first subjected our systems to 200 steps of steepest descent energy minimization, then carried out two separate stages of equilibration (in which we followed the protocol of Corey, et al. [2] unless otherwise specified). The first stage was 10 ns in duration, using Berendsen semi-isotropic pressure coupling, a 5 femtosecond time step, and a velocity-rescaling thermostat with a temperature of 310.15 K. We chose a reference pressure of 1.0 bar and compressibility of 3e-5 bar^-1^ for the x and y dimensions of the system, but used much higher values of 10.0 bar and 3e-4 bar^-1^ for the z dimension pressure and compressibility, respectively. We did this because the initial configuration of the CHARMM-GUI LPS has a fully extended O-antigen chain that exhibits a strong tendency to quickly contract. In troubleshooting the initial equilibration stages of MD, we noticed that in some cases this contraction would disrupt the stability of the bilayer itself by transiently pulling the upper leaflet away from the lower leaflet. The described semi-isotropic pressure coupling parameters successfully averted this destabilization. The second equilibration stage lasted for 100 ns and also used semi-isotropic Berendsen coupling, but with the reference pressure set to 1.0 bar and the compressibility set to 3e-4 bar^-1^ for all dimensions. Following equilibration, we carried out production MD using the Parrinello-Rahman barostat with a reference pressure of 1.0 bar and compressibility of 3e-4 bar^-1^. The timestep was set to 20 femtoseconds, and trajectory frames were recorded for subsequent analysis at 100 picosecond intervals. Per the recommendations of Jong, et al. [27], we used a van der Waals cutoff and electrostatics cutoff of 1.1 nm, a relative dielectric constant of 15, and we employed the Particle Mesh Ewald method for all long-range electrostatics terms. The production phase of all simulations involving LPS O-antigen chain lengths 0 and 2 was continued for 200 μs, while the production phase of all simulations involving LPS O-antigen chain lengths 5 and 10 was continued for 100 μs. Each of the 40 simulations required a few months of dedicated time on compute nodes equipped with current generation GPUs. To help analyze interactions between the protein and the lower leaflet lipids in our simulations, we employed gmx tools [26], Visual Molecular Dynamics (VMD) [28], and the MDAnalysis [29] package in Python. We used VMD and PyMol [30] to make system figures and we used matplotlib [31] to make all plots. We also developed our own in-house simulation analysis methods (detailed below) to more precisely and efficiently examine quantities of interest across trajectories.

### Lipid Interaction Analysis

To assess protein-phospholipid interactions in our simulations, we first calculated the average number of each type of lipid (PPPE, PVPG, and PVCL2) bound to each protein throughout the last 10 μs of each respective simulation. That is, for every given simulation frame within our time window, we iterated over all lipid molecules of each type and imposed a check on whether any CG bead of that lipid molecule was within 5.5 Å of any CG bead of the protein. In choosing this cutoff distance, we followed the example of Corey, et al. in their analysis of phospholipid interactions with inner membrane proteins [2]. Since we were mainly interested in protein-phospholipid interactions in the lower leaflet, we omitted interactions involving lipid molecules that had flipped into the upper leaflet (see below). Averaging over the frames gives the average number of each lipid type bound to the protein throughout the final 10 μs of the simulation. In an attempt to quantify relative preferences of each of the different lipid types, we normalized the calculated average bound lipid numbers by both the lateral-facing solvent accessible surface area (SASA) of each protein’s membrane-spanning region (see SI for details) and by the abundance of each lipid type. We also characterized protein-phospholipid interactions by measuring per-residue phospholipid occupancy scores across the full length of each entire simulation. This method is based on that used by Corey, et al. [2] and represents the fraction of simulation time that a particular lipid type spends within the cutoff distance (5.5 Å) from a given residue on the protein.

### Lipid Flipping Analysis

Analysis of lipid flipping events was conducted as follows. First, we defined the bounds of the membrane: the “top” of the membrane was defined by the median z-coordinate of the LPS “B20” beads, which correspond to an amide and “pseudo-headgroup” for two of the carbon tails of the LPS Lipid A moiety; the “bottom” of the membrane was defined by the median z-coordinate of PPPE headgroup beads (since this is the most abundant phospholipid in the lower leaflet). We then analyzed the positions of all the headgroups for each phospholipid molecule throughout our set of simulations and recorded their positions relative to the calculated boundaries of the membrane. Each phospholipid molecule was first given a designation of “lower” or “upper” depending on whether it started the production phase in the lower or upper leaflet, defined by its headgroup z-position relative to the membrane midpoint (in all cases, however, our phospholipids started in the lower leaflet, as we did not observe any flipping events during equilibration). An upward flip was defined as occurring when the headgroup of a lipid formerly designated as being in the lower leaflet passed above 80% of the distance from the bottom of the lower leaflet to the top of the upper leaflet. A downward flip was defined as occurring when the headgroup of a lipid formerly designated as being in the upper leaflet passed below 80% of the distance from the top of the upper leaflet to the bottom of the lower leaflet. To calculate flipping event durations, for each detected event, we designed our algorithm to search backwards to find the point in time where the flipping lipid passed 20% of the distance across the membrane and designated that point as the beginning of the flip. We also recorded all amino acid residues that were within 5.5 Å of the flipping lipid at any point from the beginning to the end of the flipping event, as well as the overall minimum distance to the protein during the course of the flipping event.

### O-Antigen Shielding Analysis

For calculating shielding of the OMPs by the O-antigens of the LPS molecules, we first projected our 3D systems into the xy plane and overlaid a 1 Å × 1 Å grid. We then identified grid “cells” containing a) any protein bead, or b) both protein and LPS beads. Dividing the latter by the former provides a general estimate of the degree to which the LPS occludes the protein since grid cells will be occupied by both protein and LPS if and only if the LPS sits directly above the protein. For OmpA and TolC, shielding was calculated only using CG beads of the transmembrane domain, since including the periplasmic domains of the two proteins would give spurious results.

### Water and Ion Analysis

To analyze the degree to which water movement across the membrane was restricted by O-antigen chain length, we developed an algorithm to detect when water molecules move across the bilayer from one side to another. The routine tracks each water molecule’s trajectory throughout the entire simulation and notes when traversals occur, discarding failed traversals. We tracked traversals in both directions and also recorded the minimum distance of each water to the protein during the water’s traversal. Sodium and chloride ion traversal events were also tracked in exactly the same manner. For these analyses, we defined the bounds of the membrane as follows: we defined the bottom of the lower leaflet for a given frame by looking at the z-coordinates of the PPPE headgroups and taking the median value. We made the top of the upper leaflet correspond to bead “B83”, which represents part of the terminal core glucose sugar on LPS. Just as with the lower leaflet, we took the median z-coordinate of the B83 beads and defined this as the top of the membrane region. We used a different definition for the top of the membrane region compared to our lipid flipping analysis (see above) in order to allow us to track when waters passed in or out of the O-antigen core region.

## Results

### Lipid Interactions Analysis

We assessed protein-phospholipid interactions first by measuring the average number of each type of lipid that was bound to the protein throughout the last 10 μs of each simulation. The results—grouped according to O-antigen chain length—are shown in Figure 2. Overall, most of the proteins exhibited a degree of variability across the four simulations (see, for example, the unusually strong binding of PVCL2 (cardiolipin) to YdiY observed during the last 10 of the simulation with zero O-antigen repeats). More consistent results were seen with the three largest proteins in the dataset, presumably because the numbers of bound lipid molecules are larger and therefore more likely to be better sampled. OmpC and OmpF, for example, showed consistent relative preferences for binding to PPPE (phosphatidylethanolamine), then PVPG (phosphatidylglycerol), followed by PVCL2. In direct contrast, TolC exhibited the opposite behavior and showing a clear preference for binding PVCL2. We also measured lipid binding by looking at the raw (non-normalized) per-residue phospholipid occupancies for PPPE, PVPG, and PVCL2, respectively (see Figures S3-S26). Occupancy values represent the fraction of simulation time a particular lipid type spent within our cutoff distance (5.5 Å) of a given residue on the protein. In general, the binding occupancies for each of the ten proteins in this analysis, regardless of O-antigen length, show patterns that are consistent with what was reported by Corey, et al. in their analysis of *E. coli* inner membrane protein-lipid binding [2]. In particular, cardiolipin tended to exhibit the highest peak lipid occupancy levels relative to its abundance in the membrane, as Corey, et al. observed for the inner membrane [2]. Consistent with the above normalized, average lipid binding results, TolC showed especially high occupancy levels for cardiolipin across its residues exposed to the inner leaflet, with its highest levels reaching nearly 50%. Meanwhile, OmpC and OmpF showed relatively low cardiolipin occupancy, tending not to reach more than 20% occupancy. Other proteins with segments that showed a predilection toward cardiolipin binding include FadL, OmpX, Tsx, and YdiY—all of which reached occupancy levels of nearly 40%.

**Figure 2:**
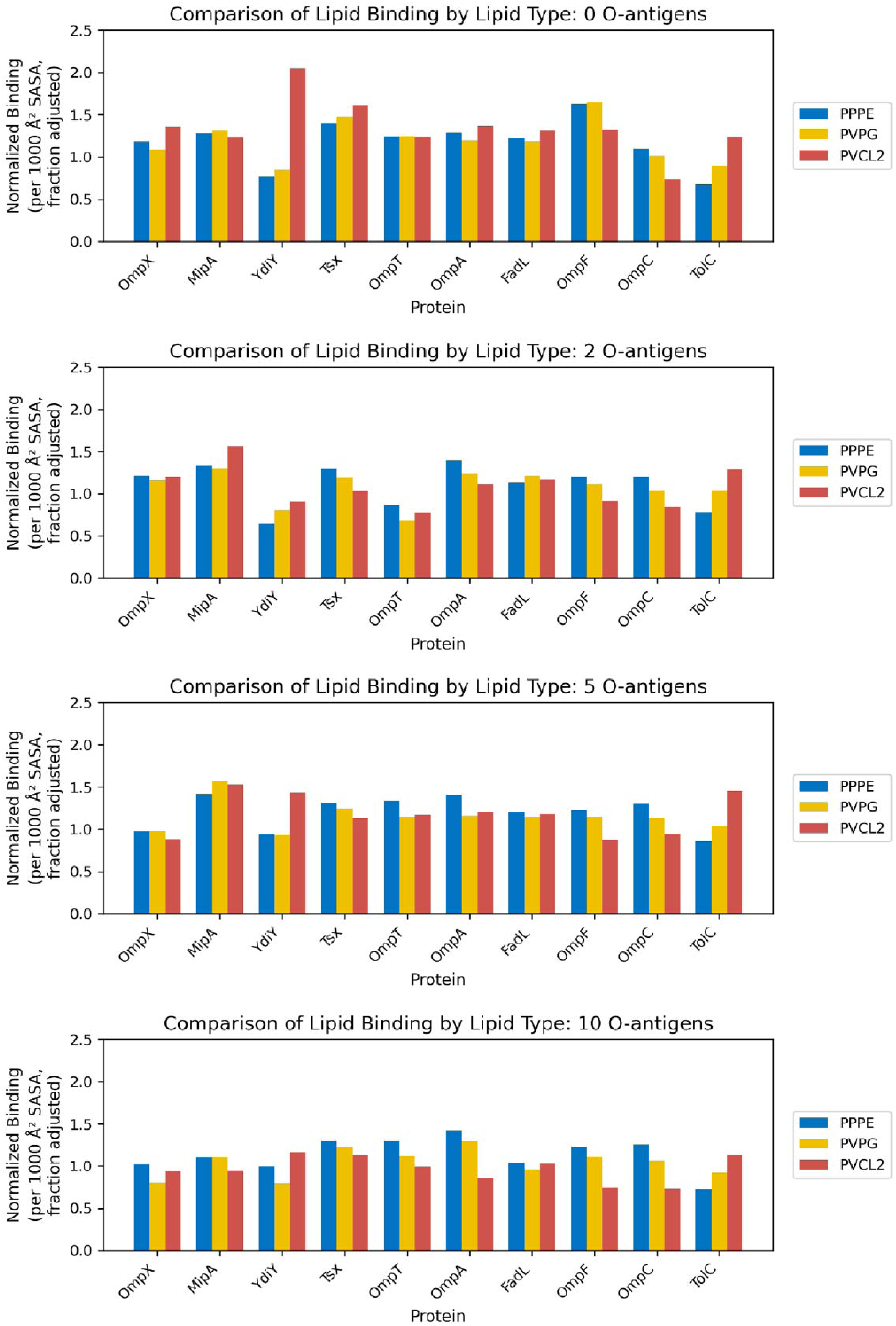
Average number of lipid molecules (of each type) bound to each protein across the entire set of 40 simulations; lipid binding numbers were normalized to the lateral facing solvent-accessible surface area of each protein, as well as to the abundance of each lipid type

### Lipid Flipping Analysis

A surprising feature that we first identified by visual analysis of the simulations was that in nearly all our simulations, phospholipids flipped from the inner leaflet of the OM to the outer leaflet (and sometimes back again to the inner leaflet). All of these flips involved either PPPE or PVPG: we did not observe any flips of PVCLs in any of the simulations. Interestingly, the percentage of flipping events that involved PVPG molecules (45%) was considerably higher than expected given its relative abundance in the membrane (20% PVPG vs. 75% PPPE) suggesting that it is significantly more prone to fliiping. A more detailed breakdown of the number of flips for each lipid type can be found in Figure S27. Figure 3 shows the number of total up and down flipping events across all 40 of our simulations, organized by protein system and by O-antigen chain length. We found that there was substantial variability from protein to protein, and even for the same protein across the four different LPS O-antigen chain lengths: for instance, the protein OmpA exhibited the most flips in the simulation with O-antigen chain length 5, whereas the protein YdiY exhibited the most flips in the simulation with O-antigen chain length zero. Not unexpectedly, in all cases, we observed a greater number of upward flips than downward flips: this is because all lipids start in the lower leaflet, so the first flip executed by any given lipid molecule must be an upward flip. We found it notable that the large-sized trimers OmpC and OmpF exhibited relatively few flips compared to the other proteins across the entire simulation set: TolC stood out as having the fewest flips of any protein in our analysis, exhibiting just one flip in the simulation with O-antigen chain length zero, and no flips at all for the three other O-antigen chain lengths.

**Figure 3:**
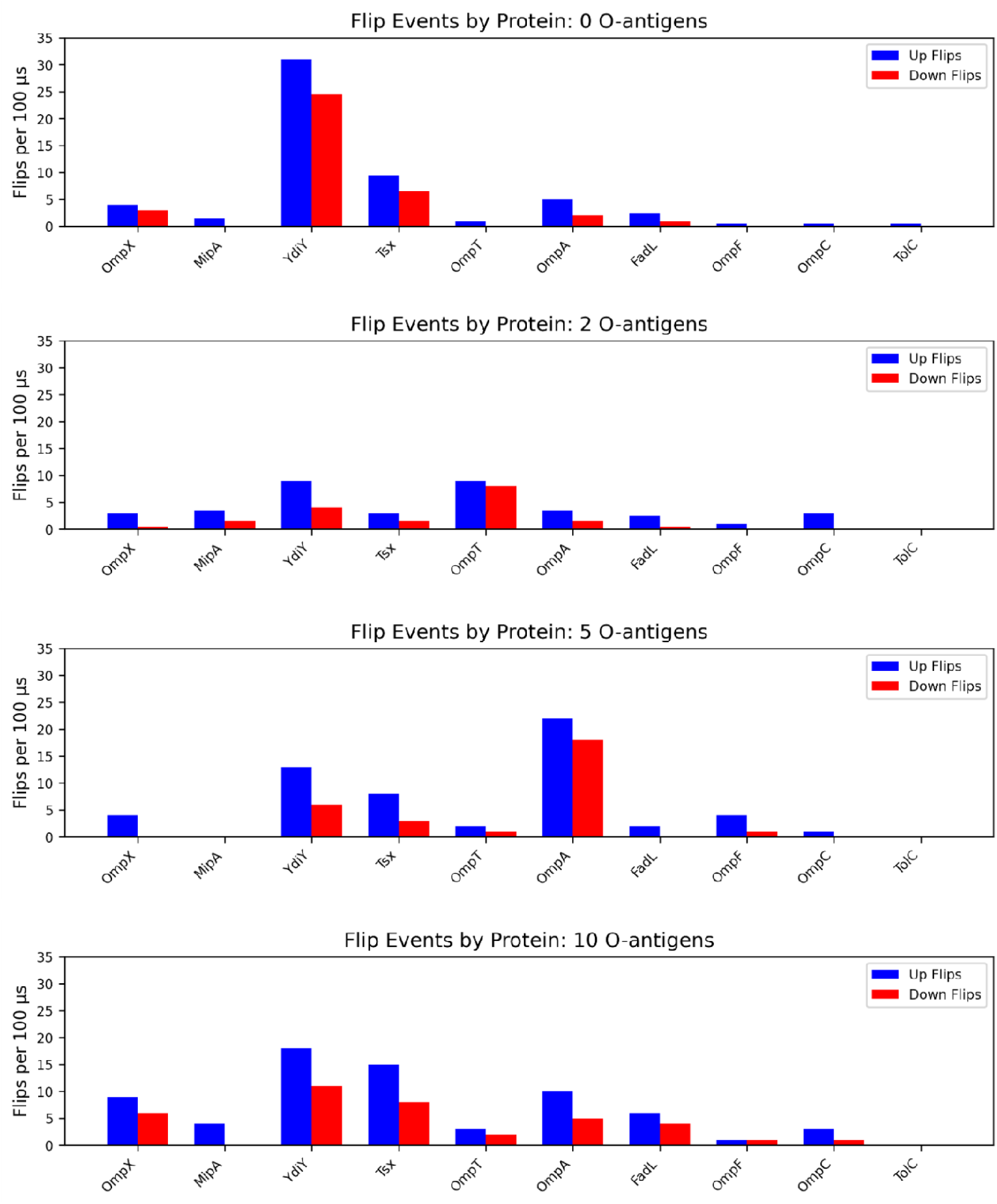
Number of flips across our 40 different simulations, arranged by O-antigen chain length and protein (with proteins ordered from shortest to longest polypeptide sequence)

Most strikingly, visual observation of the simulation trajectories indicated that the described flipping events occur exclusively at protein-lipid interfaces. Figure 4 shows a series of snapshots from our simulation of the protein YdiY with O-antigen chain length zero; everything but the protein and the flipping lipid has been hidden for simplification; the snapshots depict a PPPE /molecule starting in the lower leaflet and then flipping up to the upper leaflet. In order to ascertain whether a close interaction with the protein was a universal characteristic of all flipping events, we recorded the minimum distance to the protein during each flipping event. As Figure 5 shows, for the four sets of simulations with different chain lengths, flipping events always occurred within a relatively short distance to the protein (approximately 3 to 4 Å); this was the case for both upward and downward flips. We then explored the follow-up question of whether specific regions of a particular protein might be especially prone to inducing flipping events. Figure 6 depicts the protein YdiY from the simulation with O-antigen chain length zero, with residues colored according to their proximity to either upward (6a) or downward (6b) flipping events. One part of the surface of the protein is clearly more closely associated with flipping events, and the same surface is associated with both upward and downward flips. This same surface is also highlighted in the YdiY simulations performed with the other O-antigen chain lengths (see Figures S28 to S30) suggesting that it is, in effect, a “flipping hotspot”. Notably, the surface is enriched in aromatic sidechains (see Figure 6c), but aside from this feature it appears unremarkable, and an analysis of YdiY’s electrostatic potential, solved using PyMol’s Adaptive Poisson-Boltzmann Solver (APBS) [32], reveals no obvious difference from that of OmpF, which we use as an example of a protein associated with few lipid flips (Figure S31).

**Figure 4:**
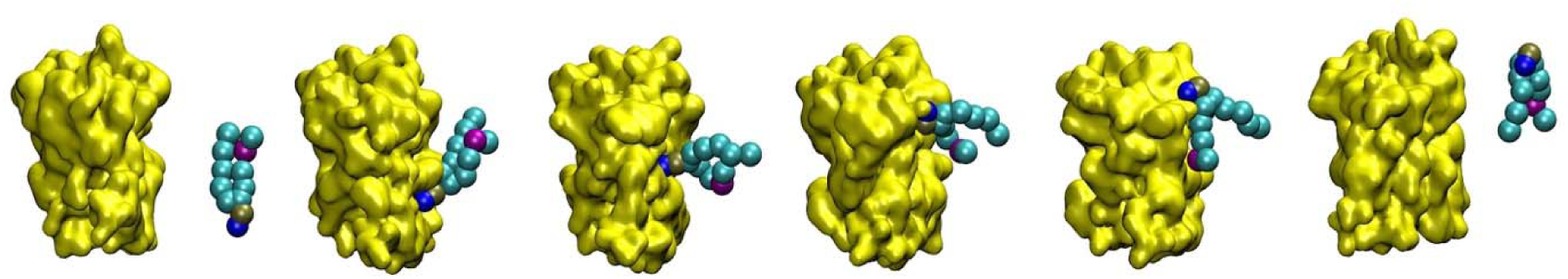
Trajectory snapshots from our simulation of YdiY with O-antigen chain length zero. The lipid is PPPE (cyan), and it begins the simulation away from the protein (yellow). As the headgroup (blue sphere) interacts with the protein, it gradually slides up the surface until the lipid has completed a 180-degree turn. The lipid dissociates at a later point.

**Figure 5:**
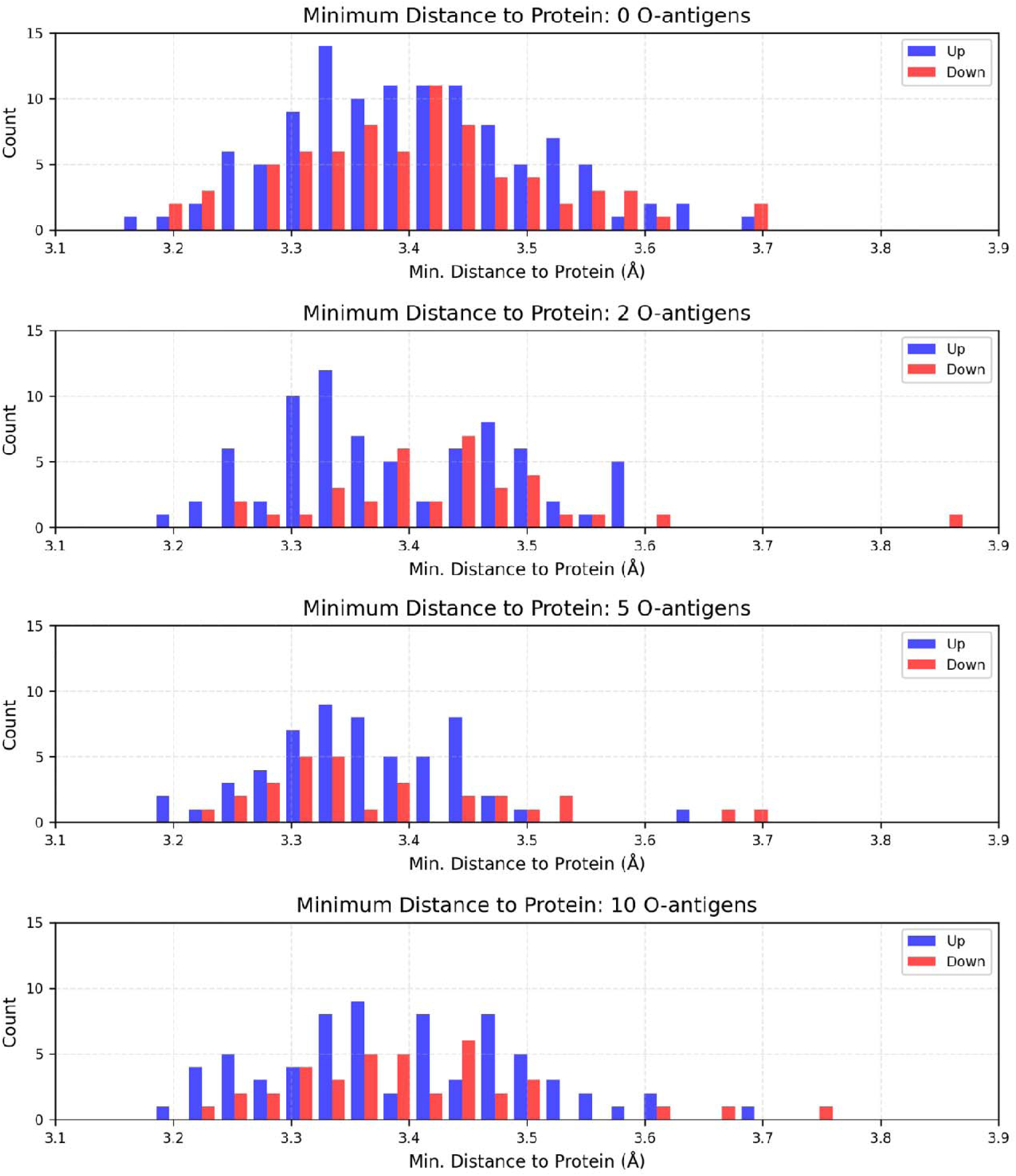
Minimum distance to protein during flipping events for each set of ten simulations with a different O-antigen chain length

**Figure 6:**
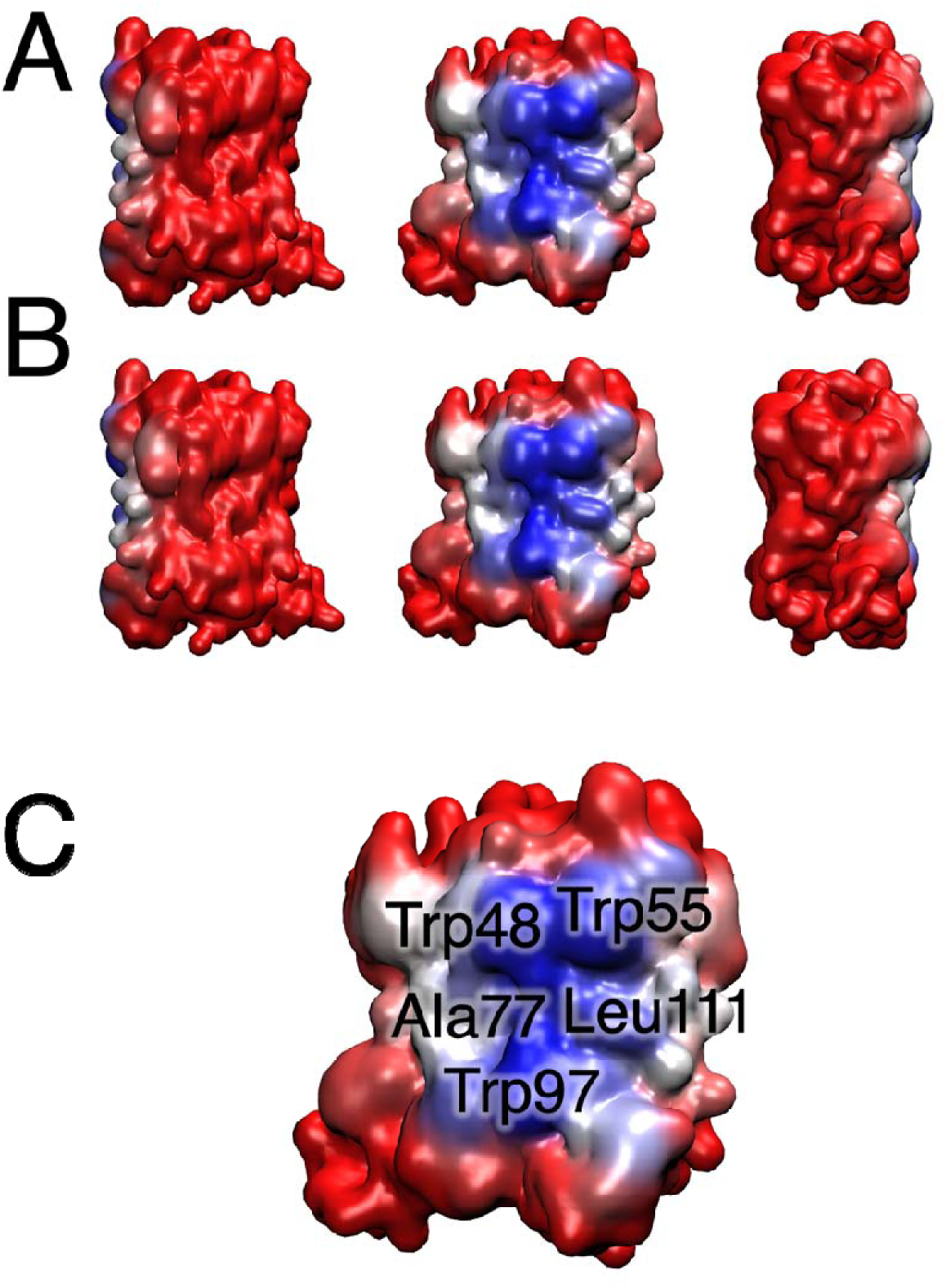
The protein YdiY from the simulation with O-antigen chain length zero, with residues colored from least likely to be associated with flipping events (red) to most likely (blue). **A** (top 3 figures) represents data from upward flips (showing sequential 120-degree views of the protein), whereas **B** represents data from downward flips. **C** labels the specific residue hotspots on the protein.

The final characteristic associated with flipping events that we measured was how long the process of flipping took. We termed this quantity the flipping event “duration.” Figure S32 shows the results from our four sets of ten simulations (as before, each set being a different O-antigen chain length). We found that the vast majority of flip durations were 50 ns or shorter. There was no noticeable difference between the duration of upward flips and downward flips. In each set of simulations, however, there were multiple outlier events that lasted up to several hundred nanoseconds.

### O-antigen Shielding Analysis

For our analysis of OMP shielding by the LPS O-antigen, we measured how shielding levels changed throughout a chosen reference simulation which we selected as OmpC with an O-antigen chain length of 10. For this simulation, we calculated shielding levels throughout the initial equilibration phase of the simulation and throughout the entire trajectory (using every 100^th^ frame). Our results (Figures S33, S34) indicate that it took approximately 20 μs for the extent of shielding to fully converge. In order to compare shielding in all 40 of our trajectories (ten different proteins, four different O-antigen chain lengths), we chose to compare the shielding percentage for the trajectory frame corresponding to 100 μs, as this represents the furthest time point that all of our trajectories reached, and as the aforementioned single trajectory results indicate, shielding values should be converged far before that point. Our results are summarized in Figure 7. As is to be anticipated, longer O-antigen chains cause a greater degree of shielding of the OMPs; the relationship between percent shielding and O-antigen chain length, however, varies substantially from protein to protein. The monomeric proteins and the single-pore trimeric protein TolC generally exhibit a substantial increase in shielding from zero to two O-antigens. The three-pore trimer proteins OmpC and OmpF, on the other hand, exhibit a comparatively smaller increase in shielding with two O-antigens, but show greater shielding with five O-antigens, and at the ten O-antigen level, their extent of shielding was on par with the rest of the protein set. As a visual example of OMP shielding, Figure 8 shows a top-down view of the four OmpC systems with different O-antigen chain lengths at the 100 μs time point of their simulations. Our results ultimately support the experimental literature suggesting that epitopes of Gram-negative OM proteins can be effectively shielded by O-antigen chains [13].

**Figure 7:**
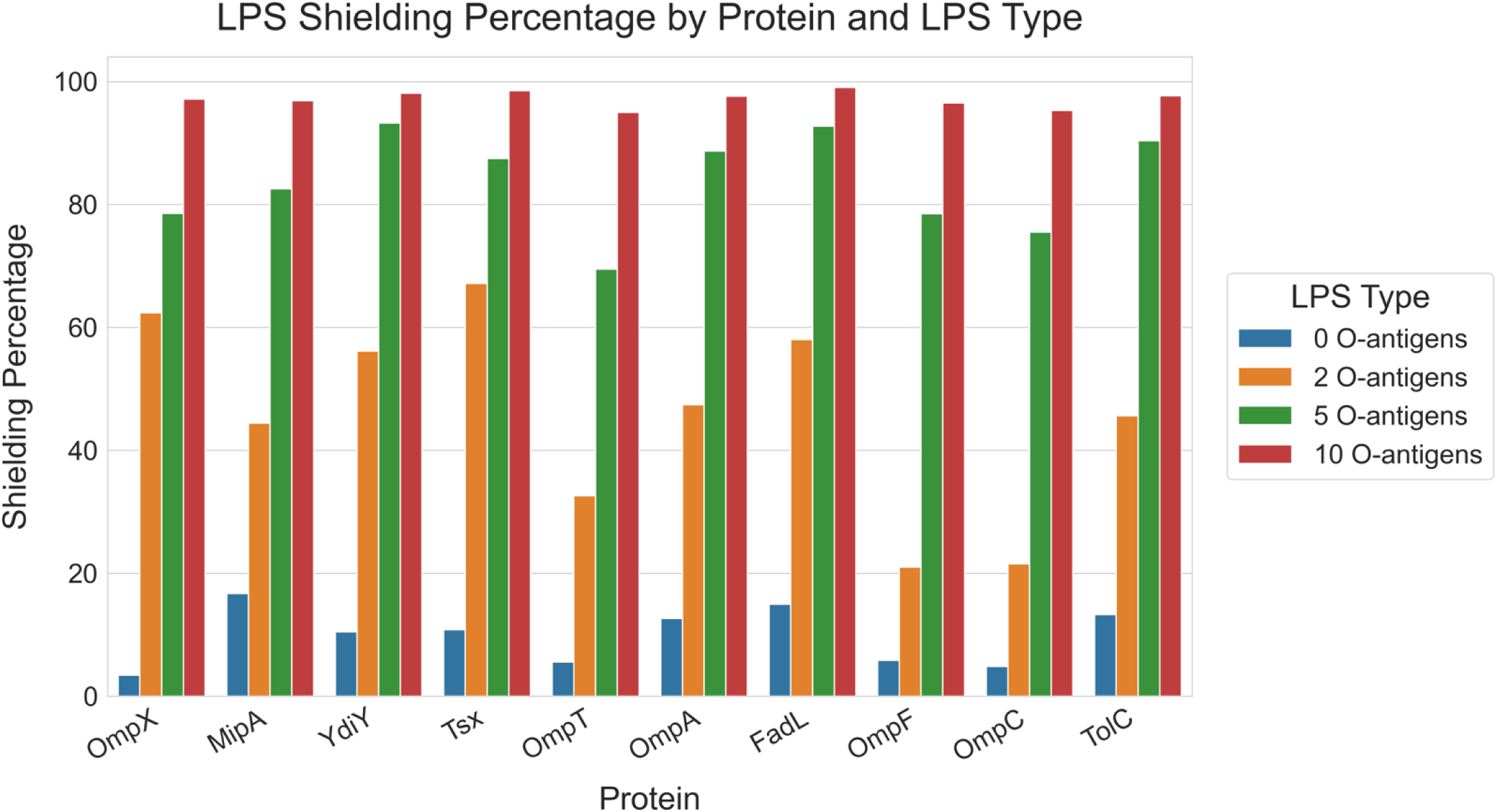
Shielding level at 100 μs into each respective simulation, organized by protein and LPS O-antigen chain length

**Figure 8:**
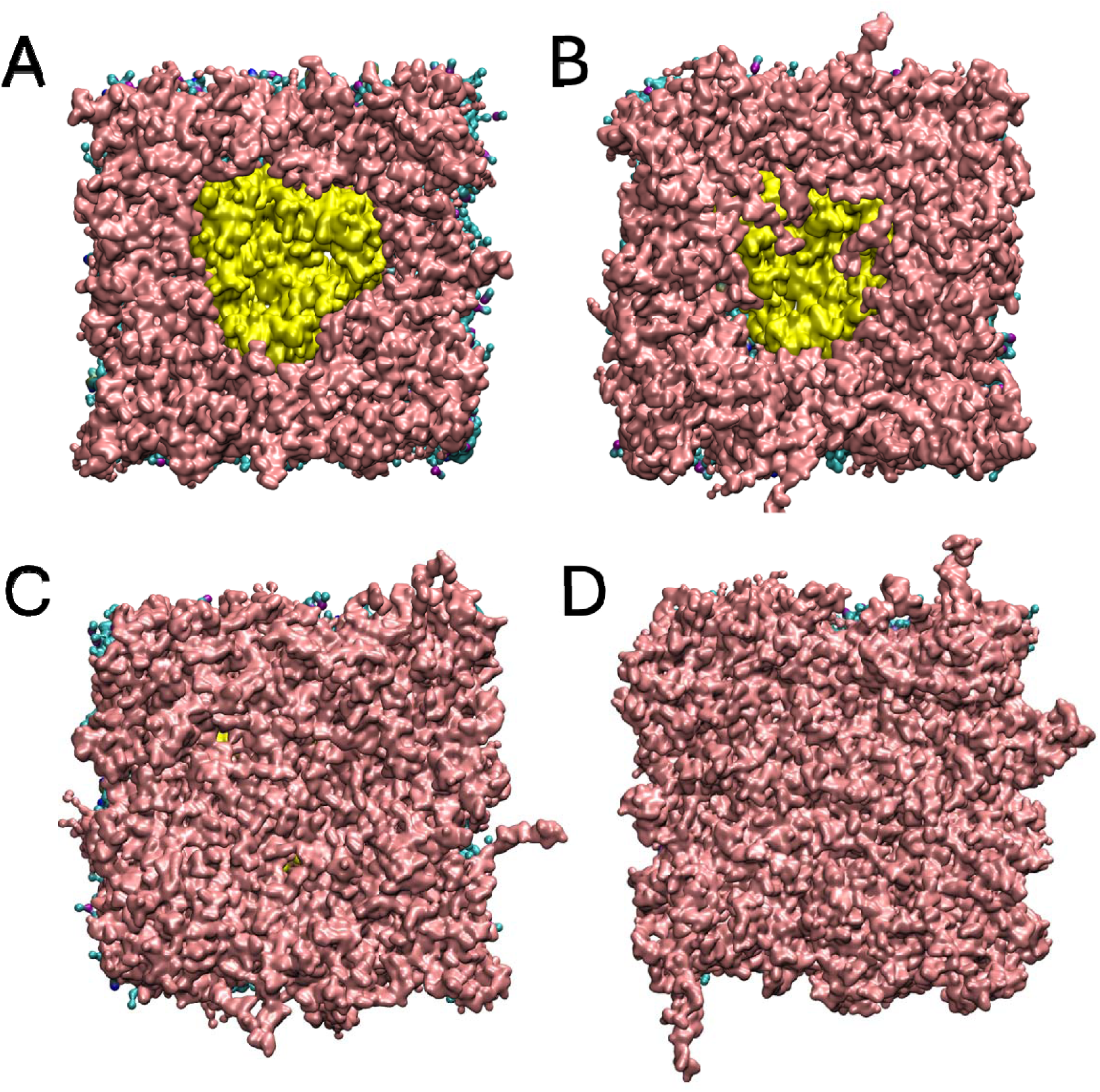
Top-down, surface view of the four OmpC systems after equilibration. LPS with 0 O-antigens exhibited the least shielding (A), and LPS with 10 O-antigens exhibited the most (D). Lengths 2 (B) and 5 (C) were intermediate. LPS is depicted in pink, while the protein is depicted in yellow.

### Water and Ion Movement Analysis

We originally hypothesized that longer O-antigen chains would effectively restrict water movement through the OmpC and OmpF porins by sterically blocking access to the pore from the extracellular side of the membrane. To test this hypothesis, we tracked water “membrane traversal” events across all 40 protein systems in our analysis. The results are displayed in Figure 9, with simulations grouped according to O-antigen chain length. Along with TolC, the OmpC and OmpF porins generated vastly more water traversals than the monomeric proteins, but non-negligible levels of water traversal were observed in all 40 simulations. Of the monomeric proteins in the dataset, YdiY again exhibited the most water traversals.

**Figure 9:**
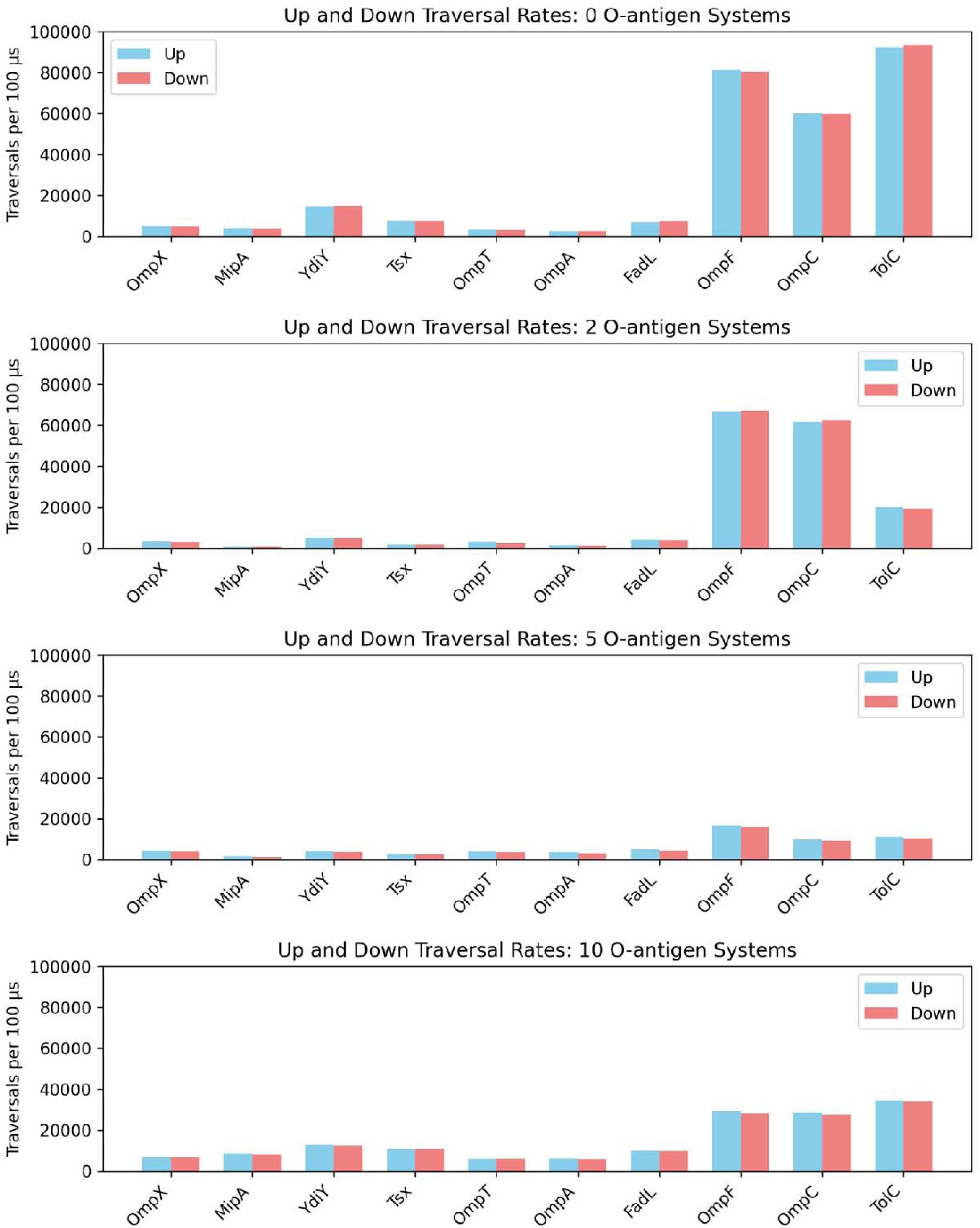
Rates at which waters traversed across the membrane (per 100 μs) for each respective protein system and each of the four O-antigen chain lengths

We found that longer O-antigen chains restricted water movement through OmpC and OmpF, as the traversal counts for the simulations with O-antigen chain lengths five and ten were generally lower than for O-antigen chain lengths zero and two. Interestingly, TolC saw the most drastic decrease in traversal events going from zero to two O-antigens, whereas the decrease from zero to two O-antigens for OmpF was much less, and in the case of OmpC there was surprisingly a small increase. TolC’s decrease can be partly explained by the previously discussed O-antigen shielding characteristics, where TolC was more extensively shielded by two O-antigens compared to the OmpC and OmpF porins. Figure 10 provides a visual depiction of OmpC vs. TolC shielding at 100 μs in their respective simulations, demonstrating that TolC is almost completely obscured while OmpC is still largely exposed. Unexpectedly, a greater number of water traversals were observed in the systems with ten O-antigens compared to the systems with five O-antigens. This was the case not only for the two porins and TolC, but also for most of the non-porin monomers. An extensive investigation of this phenomenon has not yet produced any clear explanation.

**Figure 10:**
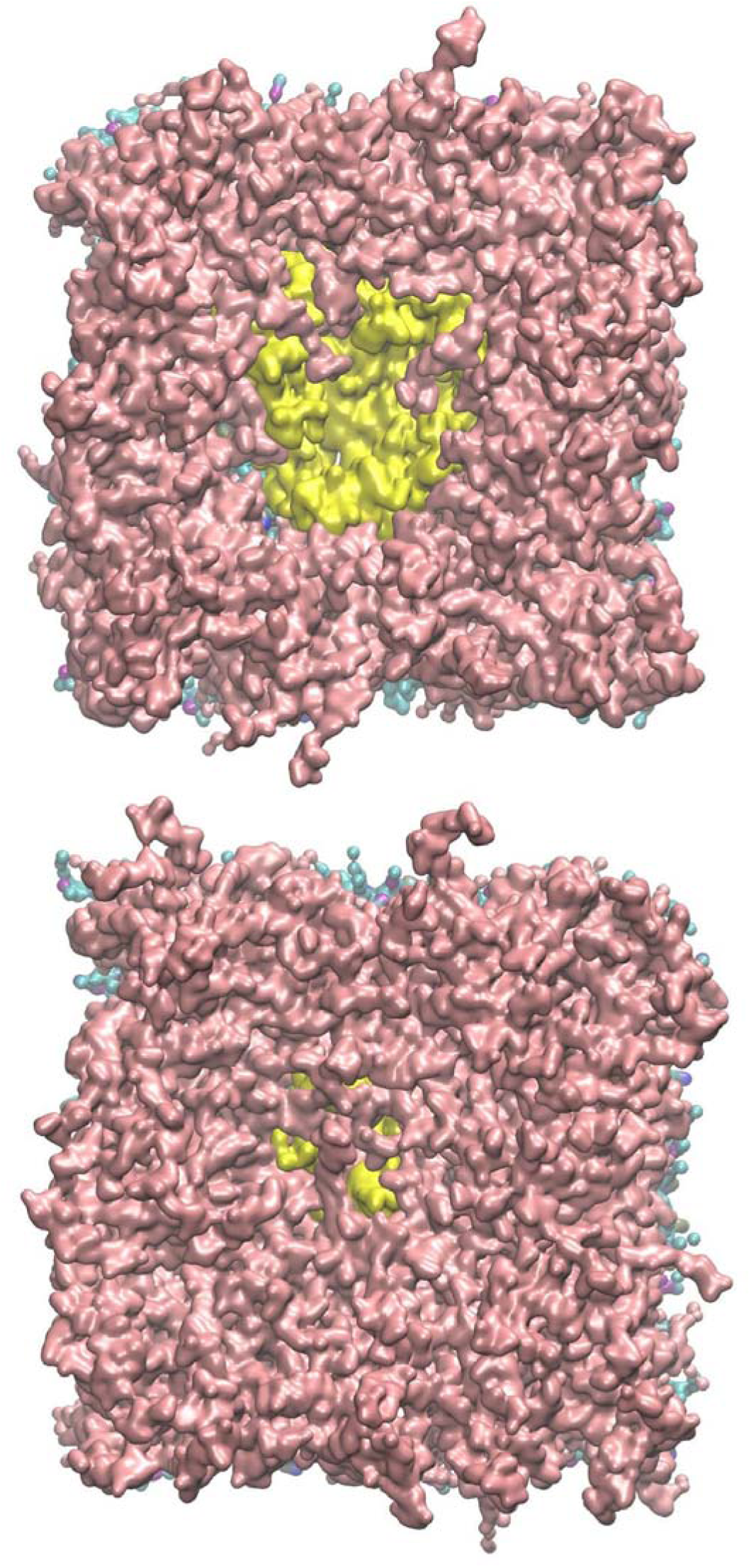
TolC (bottom) is much more extensively shielded by 2 O-antigen LPS compared to OmpC (top).

We wanted to assess the extent to which water traversal events were associated with proximity to the proteins. Therefore, for each water traversal event, we measured the minimum distance to the protein that the given water achieved during its traversal. These results are displayed in Figure S35. Aggregating the results for each O-antigen chain length set, the vast majority of all traversal events took place very close to the protein (approximately 3-5 Å; see also Figure S36). This was the case for all 4 O-antigen chain lengths. However, there were also a non-negligible number of traversal events extending away from the proteins, up to more than 60 Å. This suggests that while the proteins facilitate water traversals across the membrane, waters are also able to traverse the membrane throughout most of the simulation box.

To gain a clearer picture of how waters traverse the membrane in the OmpF and OmpC porin protein systems, we also measured the minimum distance to any of the three porin pore centers that a given water achieved during its traversal. Pore centers were defined as the midpoint of two manually chosen residues on either side of each β-barrel opening of the porin trimers. The results (Figure S37) clearly show that the majority of waters from these systems traverse the membrane through the pores. However, there is a slight bimodality to the data, with a relative peak appearing at approximately 18 Å from the pore centers. This roughly corresponds to the minimum radius of each β-barrel with respect to its pore (the pores are not directly centered in the β-barrels), suggesting that the waters from the 18 Å peak are crossing at the interface of the protein and the membrane bilayer.

Finally, we measured the duration of the water traversal events (Figure S38), defined as the amount of time each traversing water spent within the membrane region during its traversal. Most traversals were relatively short, with the highest peak located at the far left of the distribution. However, the range of traversal durations spanned nearly the entire length of the trajectories, indicating that a small subset of waters from the simulations take an especially long time to traverse their respective membrane protein systems. Since we only counted completed traversal events, it is possible that our analysis misses potential traversals in progress that are slated to take even longer than the length of our simulations. This explains the discrepancy in maximum traversal lengths for the set of simulations with O-antigen chain lengths zero and two (which were run to 200 μs) compared to the set of simulations with O-antigen chain lengths five and ten (which were run to only 100 μs).

In addition to analyzing water movement across the outer membrane in our OM protein simulations, we also investigated the movement of sodium and chloride ions. It should be noted that there is a greater number of sodium ions in each simulated system, because a substantial surplus of sodium ions—approximately twice the number of chlorides in the case of OmpC and OmpF, and 40% greater for TolC—was required to neutralize the negative charges on the protein and lipids. For this reason, we calculated ion traversals in terms of “per 100 ions” present in the system. Apart from that, our ion analysis was conducted in an identical matter as for water, looking specifically for ions that managed to cross entirely from one side of our defined membrane region to the other. The upward and downward sodium traversals we detected for the various protein systems and LPS chain lengths are shown in Figure 11. For the zero O-antigen system set, TolC exhibited the highest number of sodium traversals, followed by OmpC and then OmpF. MipA, YdiY, and Tsx exhibited a very small number of traversals, while OmpX, OmpT, OmpA, and FadL exhibited no sodium traversal events at all. For the two O-antigen systems, TolC’s traversal numbers were greatly reduced. OmpC and OmpF’s numbers were also lower, albeit to a lesser extent than TolC. This mirrors what was seen for the water traversal analysis (Figure 9) and is explainable by the degree of shielding for TolC vs. the porins with two O-antigens (see Figure 10). For the five and ten O-antigen systems, traversal numbers were generally lower for OmpC, OmpF, and TolC.

**Figure 11:**
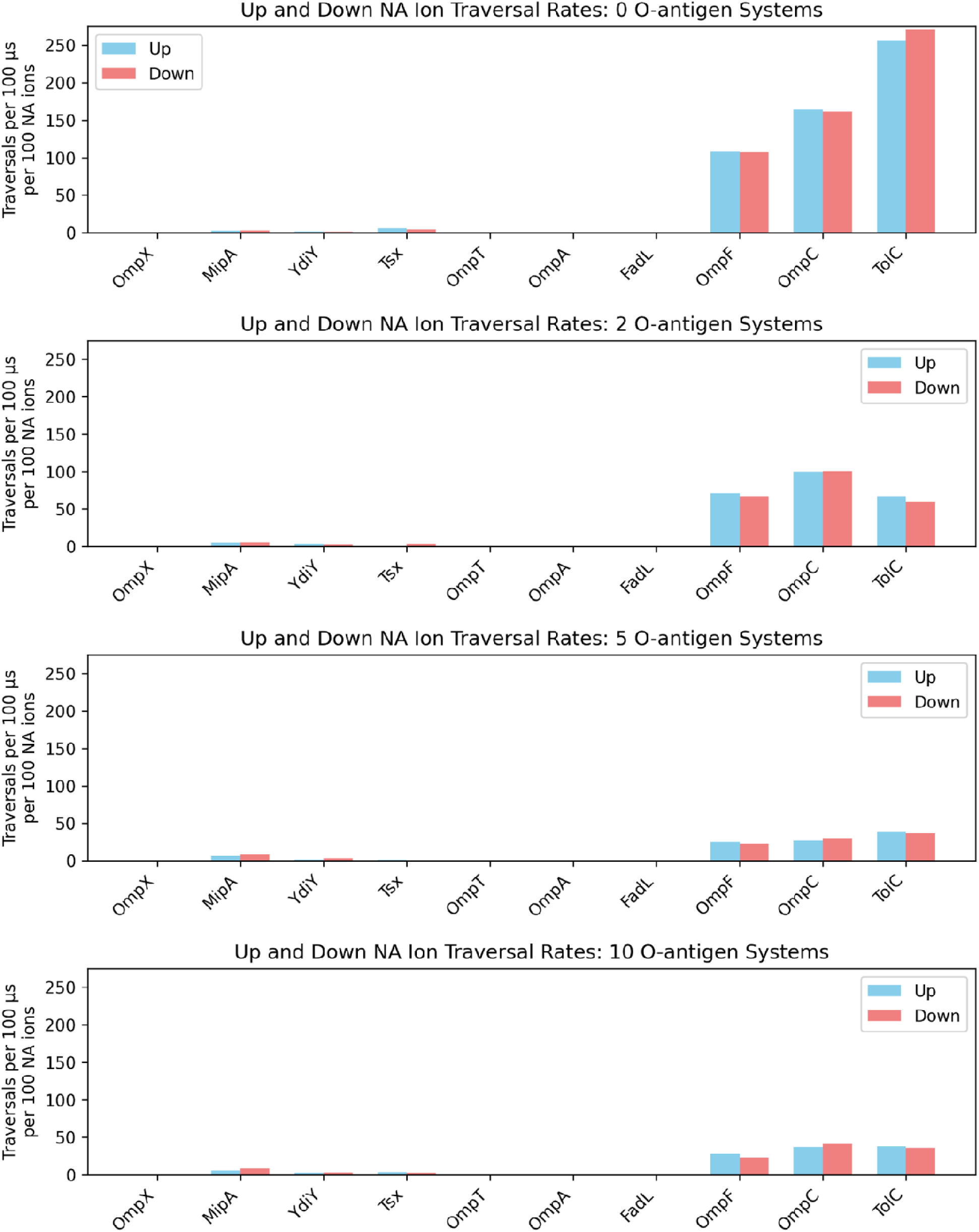
Rate at which sodium ions traversed the membrane in each respective protein simulation across the four O-antigen chain lengths

In terms of distance to the protein, the sodium ion membrane traversal events generally took place directly adjacent to the protein, although a handful of events from the zero antigen systems took place up to 20 Å from the protein (see Figure S39). This is consistent with the water traversal results, where most traversal events also occurred adjacent to the proteins, but some took place away from the protein. The “distance to nearest pore” calculations (Figure S40) illustrate that for the OmpC and OmpF systems, almost all of the sodium ion traversal events take place via a pore (save for a few traversal events in the zero O-antigen systems). The range of traversal durations (Figure S41) was substantially less than for water, with the longest durations in the zero and two O-antigen simulations (total simulation length: 200 μs) extending to 125 μs, and the durations for the five and ten O-antigen systems (total simulation length: 100 μs) being all under 50 μs.

Chloride ion traversal events were recorded and characterized in the same manner as sodium ion traversal events. As before, the overwhelming majority of traversal events took place in the systems with OmpF, OmpC, and TolC (Figure 12). For the set of simulations with O-antigen chain length zero, TolC saw the highest number of traversals, followed by OmpF and then OmpC. The MipA and Tsx systems also saw a very small number of traversals. TolC’s traversals were once again drastically reduced in the two O-antigen systems, likely due to the relatively high degree of shielding TolC experiences at this O-antigen chain length (see Figure 10). The most striking difference between the chloride and sodium ion traversal results was the switch in ion traversal preference for OmpC and OmpF: with sodium, OmpC exhibited more traversals than OmpF, but with chloride, OmpF exhibited more traversals than OmpC.

**Figure 12:**
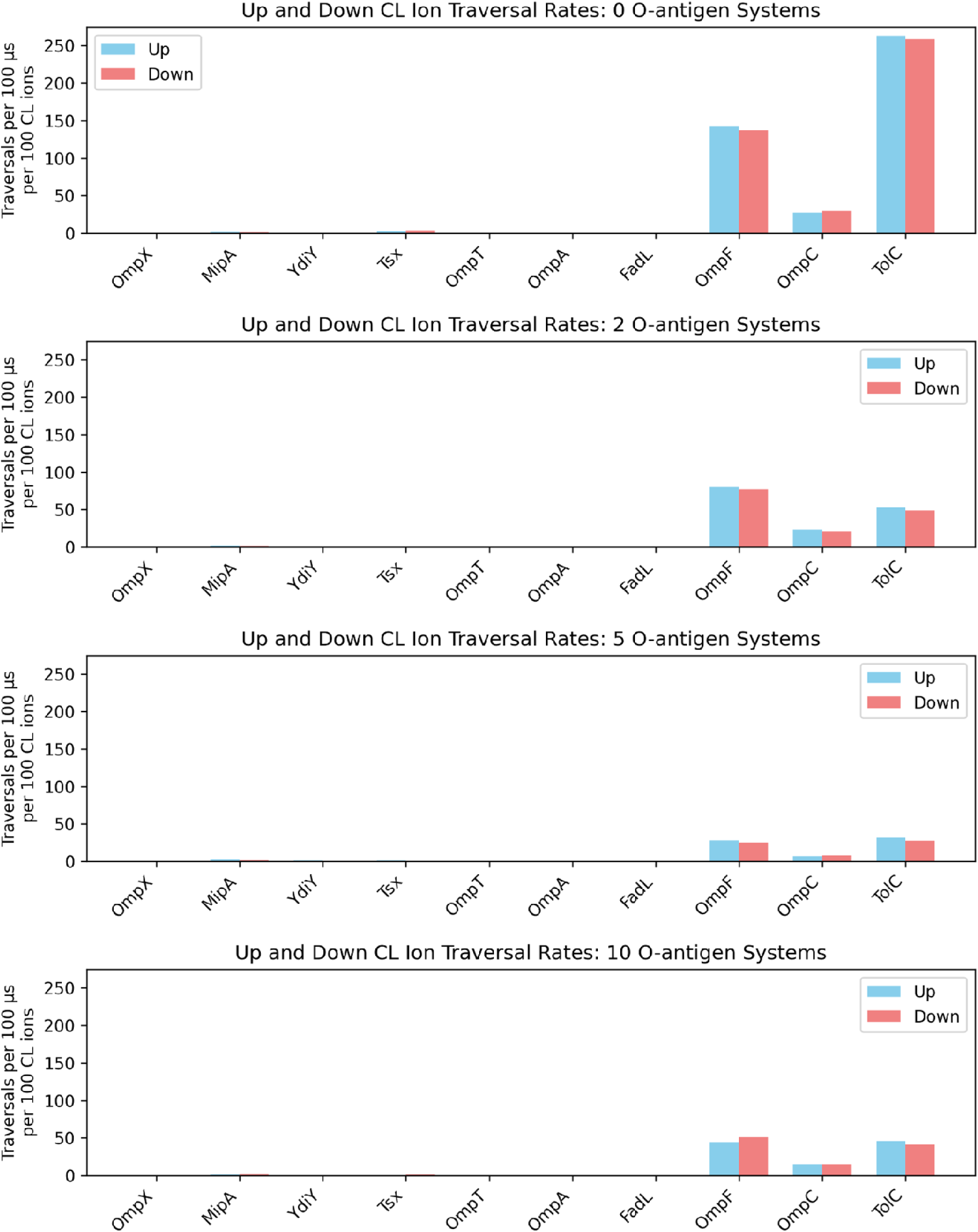
Rate of chloride ion membrane traversal events in the various protein systems, grouped according to O-antigen chain length

The “minimum distance to protein” (Figure S42) and “minimum distance to nearest pore” (Figure S43) calculations for chloride make it clear that all chloride traversals for OmpC and OmpF took place via a pore. Unlike the results for sodium traversals, there were no exceptions where ions traversed the membrane far from the protein and/or far from a pore. Interestingly, the chloride traversal event durations (Figure S44) were overall shorter than for sodium.

## Discussion

The simulations reported here have produced a range of findings both expected and unexpected. Our analysis of protein-phospholipid interactions reveals that, at least according to the Martini3 forcefield, TolC exhibits particularly high affinity for PVCL2 (cardiolipin), suggesting that this lipid may play a regulatory role in TolC’s functionality. Interestingly, although there is no explicit evidence for this in the literature to date, there is work that shows that cardiolipin interacts favorably with AcrB. As discussed previously, AcrB, AcrA, and TolC form an *E. coli* trans-envelope complex, with AcrB being embedded in the *E. coli* inner membrane. Du, et al. [6] used coarse-grained molecular dynamics simulations to show that cardiolipin in the inner membrane preferentially interacts with AcrB and Acrz and then performed experimental work to show that the interaction has functional consequences. It is interesting to note, therefore, that both the IM-embedded and OM-embedded protein components of the AcrAB-TolC trans-envelope complex appear to show a preference for binding the same phospholipid.

One phenomenon we did not initially expect to observe was the occurrence of lipid “flipping” events throughout the overwhelming majority of our simulations; these events always took place next to the protein (at a distance of roughly 3-4 Å). This observation is interesting in light of the fact that, in the literature, defects in protein trafficking to the OM are associated with the presence of phospholipids in the outer leaflet [12]. While the latter results might be interpreted as indicating that the absence of proteins in the OM is associated with flipping, it is possible that—in the presence of trafficking defects—the proteins that do make it to the OM still facilitate flipping events. We are not aware of previous work in the literature that describes lipid flipping for *E. coli* OM protein simulations. There have been reports from computational simulations of LPS in the absence of OMPs that suggest the presence of phospholipids in the outer leaflet of the OM imparts certain structural properties that make the bacteria less resistant to surface tension disturbances that can lead to membrane rupture [33]. This is consistent with other data reporting that the loss of OM leaflet asymmetry is a sign of physiological stress [34]. *E. coli* has protein machinery that flips phospholipids from the outer leaflet back into the inner leaflet, which provides further evidence that maintenance of an asymmetric OM is physiologically important [35]. Our simulations suggest that OM flipping can occur spontaneously at OMP-lipid interfaces in the OM, which may mean that bacteria are constantly fighting to counteract a basal level of passive phospholipid diffusion to the outer leaflet. The driving force for this phenomenon may be a simple case of mass action: due to the intrinsic asymmetry of the outer membrane, there is a strong concentration gradient between the inner and outer leaflets. This gradient is at least partly maintained by the energy input of ATP hydrolysis in the protein machinery that traffics phospholipids out of the outer leaflet [35]. In this context, the flipping of phospholipids into the outer leaflet is thermodynamically independent of OMPs and would be expected to occur without them on a long enough timescale. The present simulations, however, suggest that OMPs can be proficient “flipping catalysts” that speed up the kinetics of inter-leaflet lipid diffusion. The fact that we saw zero flips occur away from a protein suggests that non-catalyzed flipping occurs over a much longer timescale than our simulations.

The degree of lipid flipping that we observed in our simulations varied considerably, with some proteins—such as YdiY—being associated with a relatively high number of flips, and other proteins— such as OmpC, OmpF, and TolC—being associated with very few flips. This observation indicates that protein composition may play an important role in regulating the stability and integrity of the outer membrane. An interesting follow-up experiment would be to overexpress YdiY in *E. coli* and assess whether this leads to a higher proportion of phospholipids in the outer membrane after exposing the cells to stressors. It would be interesting to assess YdiY’s predilection toward flipping events in the context of its function, but as of this writing, it remains unknown [36].

As we expected, longer O-antigen chains consistently exhibited a greater level of OMP shielding than shorter O-antigen chains. The relationship between shielding percentage and O-antigen chain length does not appear to be strictly linear, however. Both our analysis of shielding in the individually analyzed OmpC simulation and our broad analysis of shielding in late production-phase snapshots for the 10-protein set support the existence of a multifactorial relationship between O-antigen chain length and shielding level that varies from protein to protein. This may have important implications for how bacteria interact with host immune components. As discussed previously, there is evidence that O-antigen chains can act to shield specific epitopes [13]. The work described here indicates that, for the pathogenic strains that express long O-antigen chains, immune factors can more readily access the large porin proteins of the OM compared to smaller monomers.

Regarding water behavior, we found that longer O-antigen chains generally inhibited water movement through the OMP porins, and we also conducted a detailed inspection of the ways that water interacts with and moves through the *E. coli* outer membrane. We found that water moved readily through the OMP porins as well as through the TolC efflux pump, but we also discovered that water was able to cross the membrane in the monomeric OMP systems. Distance measurements indicated the latter traversals occur primarily at the interface between the proteins and the OM bilayer. One other possibly interesting feature of the observed results is the fact that we saw greater water movement through the OmpF porin than through the structurally similar OmpC. Experiments measuring the flow rate of organic solutes through these two different proteins have shown that flow is greater through OmpF than OmpC, which is attributed to OmpF’s larger pore size [37]. If pore size is indeed the main factor, it is not surprising that this feature would extend to the solvent. One rather unexpected finding associated with our water analysis was that the ten O-antigen systems exhibited a higher number of traversals than the five O-antigen systems. At this point, we do not have a satisfactory explanation for this phenomenon, though we speculate that it may have to do with certain features of LPS packing around the proteins.

We also tracked ion movement and discovered that while ions were able to pass readily through the well-defined protein pores of OmpC, OmpF, and TolC, they had a much more difficult time passing next to or away from the proteins. Consequently, the proteins without well-defined pores saw negligible numbers of ion traversals across the membrane. One interesting feature of all four O-antigen chain length systems is that OmpC always exhibited a higher number of sodium traversals than OmpF. The experimental literature reports that OmpC is more cation-selective than OmpF, which aligns with what was observed here [38].

We recognize that there are a number of caveats and limitations associated with the present work. First and foremost, coarse-graining eliminates molecular details that may be important for accurately reproducing dynamic molecular behavior. We do note that *Martini3* is well-regarded and widely used, but other work from within our lab has shown hints that even all-atom force fields can produce physiologically questionable behavior—such as the tendency for protein-protein and protein-lipid interactions to be overly “sticky” [39]. Another clear deficit of the described work is that it is based on single protein simulations. In reality, the outer membrane is a densely packed, heterogeneous array of many kinds of proteins. It is certainly possible that our results regarding protein-phospholipid interactions and OMP shielding by O-antigens would be different in a multi-protein setting. For example, there may be proteins that sequester specific lipid types (like cardiolipin) in a heterogeneous, multi-protein array, reducing the degree of interaction with the rest of the proteins. If this is the case, our average lipid binding and per-residue occupancy scores may be overestimating protein-lipid interactions for a specific subset of proteins.

## Notes

### Competing Interest Statement

The authors have declared no competing interest.

