## Supporting Information for "Lipid Flipping, O-antigen Shielding, and Water Dynamics Revealed by 100-200 µs Coarse-Grained Simulations of *E. coli* Outer Membrane Proteins"

**Extended Methods**

**Protein Structure Preparation**

We referenced each protein’s UniProt [13] entry to determine whether the protein in question is expressed as a monomer or multimer. For the aforementioned list of proteins, OmpC, OmpF, and TolC are reported to be homotrimers, while the rest are all monomers. To obtain the initial 3D structure of each protein, we decided to take advantage of the vast wealth of data from AlphaFold2 [14] predictions. For the seven monomers, this was a straightforward process, because the authors of AlphaFold2 have provided the public with a database of 3D models for every *E. coli* gene [15]. In the case of the three multimers, however, we implemented AF2-Complex [16], which allowed us to generate models in their known oligomeric states as reported by UniProt. Although we recognize that all our AlphaFold-derived models are merely predicted structures, we chose this route for several reasons. To begin, it has been widely agreed that AlphaFold represents a major leap forward in the quality of predicted protein structures relative to previous methods [17]. Moreover, a core benefit of using AlphaFold2’s comprehensive database of *E. coli* protein structures is that they are guaranteed to be full-length, complete models with no gaps, “stubs”, or otherwise missing atoms. Although it is true that the finer-grained conformational details of AlphaFold2 models may deviate from experimentally determined structures, we note that our approach ultimately involves coarse-graining that eliminates such details anyway. As further justification, Figure S1 includes snapshots of our various AlphaFold2 models colored by pLDDT score; pLDDT stands for “predicted local distance difference test” and represents the general confidence of the AlphaFold2 prediction [18]. All the predicted OM protein structures have relatively high confidence. Meanwhile, Table S2 shows the piTM and Interface scores for our AF2-Complex predicted structures; these scores are confidence metrics specific to the AF2-Complex method. Like the monomers, all of the complexes have reasonably high confidence, which reflects the fact that their experimental structures have all been solved.

Since this study is primarily focused on examining outer membrane proteins in their fully processed state, the first step in processing our input structures was to remove the peptide signal sequence from the AlphaFold2 models (and also omit the signal sequence when conducting our AF2-Complex predictions for the three proteins expressed as homotrimers). We obtained signal sequence information for each protein from UniProt. Our AF2-Complex results provided us with five putative models for each of the three homotrimers, from which we simply chose the structure with the highest sum of the piTM and Interface scores. We then submitted our final ten different protein models to the Positioning of Proteins in Membranes (PPM) server to predict their orientation in the bacterial OM [13].

The biological functions of our set of 10 proteins are variable; OmpC and OmpF are known to be porin proteins. They engage in non-specific solute transport, and their expression levels can change in response to changes in growth medium osmolarity. OmpC tends to be preferentially expressed at lower osmolarity levels, while OmpF is expressed more at high osmolarity levels [21]. OmpA facilitates a noncovalent attachment between the outer membrane and the peptidoglycan layer underneath. It achieves this via its 2-domain structure, with the N-terminal domain being a β-barrel that embeds itself in the OM, and the C-terminal domain being a globular unit sticking into the periplasm that non-covalently binds to peptidoglycan [22]. The two domains are linked by an unstructured loop. OmpT, meanwhile, is a surface membrane serine protease that cleaves a variety of extracellular substrates. It has specificity towards pairs of basic residues and plays a role in bacterial virulence and resistance to antibiotics, e.g. by cleaving protamine and other antibiotic peptides and by activating human plasminogen to plasmin [23]. OmpX is part of a family of conserved proteins that participate in surface adhesion and facilitate bacterial entry into mammalian cells, and accordingly, it also plays a major role in bacterial virulence [24]. The function of MipA is less clearly established. Its name is derived from “MltA-interacting protein”; MltA lyses peptidoglycan, and MipA is thought to play a role in coordinating the enzymes that facilitate the reorganization and septation of the peptidoglycan wall [25]. The protein Tsx is a nucleoside-specific channel but is also reported to serve as a receptor for bacteriophages or certain antibiotic molecules such as colicin [26]. TolC is a well-characterized efflux pump that plays a major role in the development of multidrug antibiotic resistance [27]. It is also able to secrete toxins and proteases involved in bacterial virulence. FadL’s name is derived from its well-known role as a channel protein for long-chain fatty acid transport, and the protein is also known to be the receptor for the bacteriophage T2. Finally, there is a comparative lack of information available on YdiY relative to the aforementioned nine proteins. According to the *E. coli* database EcoCyc [28], its amino acid sequence indicates that it may be an outer membrane receptor, and other sources have characterized the protein as being acid-inducible [29].

**Coarse-grained Parameterization**

In determining how to coarse-grain our lipids, we wanted to use *Martini3* models that would differentiate between the different length aliphatic tails that characterize both PVPG and PVCL2 (16C and 18C)—which the base *Martini3* lipid library does not do. We therefore followed the work of Empereur-Mot, et al. [20], who developed a protocol—called *SwarmCG*— for differentiating between otherwise identically modeled lipids in the original *Martini3* implementation. Our atom-to-bead mapping scheme and parameter definitions are pictured and labeled in Figure 2, in which we borrow the bond and angle nomenclature of Empereur-Mot, et al. For PPPE and PVPG, we started with the base *Martini3* bead types and parameters for POPE and POPG (which have the same headgroup and similar length aliphatic tails to PPPE and PVPG, respectively) and then—where appropriate—swapped out the original bead types and bond/angle parameters with bead types and parameters from the POPC model devised by Empereur-Mot, et al. In the course of this process, there was only one case for each lipid in which the chemical architecture of the aliphatic tails could not be directly reflected from the POPC parameters. In the case of PPPE, this was for the angle formed by the *Martini3* beads SC1-C1-C4h comprising the 16C aliphatic tail. For PVPG, the ambiguous parameter was the angle formed by beads C1-C1-C4h of the 18C aliphatic tail. In both instances, however, we modeled the ambiguous parameters according to the most chemically similar entity we could identify. The POPC molecule from Empereur-Mot, et al. has an angle parameter for beads SN4a-C1-C4h, and since SN4a and SC1 are similarly sized beads, we deemed this angle an appropriate replacement. In a similar manner, we noticed that the POPC molecule from Empereur-Mot, et al. has an angle parameter for C4h-C1-C1, which is the mirror image of the ambiguous angle for PVPG, and hence we also deemed this to be an appropriate replacement. Once we had completed our *Martini3* model for PVPG, we were quickly able to define a *Martini3* model for PVCL2, since its aliphatic tails are identical to those for PVPG. To create mapping files that would allow us to transform the all-atom PPPE, PVPG, and PVCL2 molecules from our CHARMM-GUI systems, we employed the web-based utility *CGBuilder* [21].

For LPS, we employed the *Martini3* model designed and parameterized by Ayappa, et al. [3]. However, the number of O-antigens contained in their model is limited to two, and since we wanted to construct systems with longer O-antigen chains, we devised a method to extend the parameters from the terminal O-antigen and apply the same values to every O-antigen appearing thereafter in the 5 and 10 O-antigen systems. Similarly, we applied the atom-to-bead mappings for the final O-antigen in the LPS model devised by Ayappa, et al. to all the additional O-antigen subunits included in our 5 and 10 O-antigen LPS models. The last part of our all-atom CHARMM-GUI systems to coarse-grain was the protein component, for which we employed *Martinize2* [22] to construct coarse-grained elastic network models— using a force constant of 500 kJ/mol/nm^2^ and lower/upper bond cutoff distances of 0 and 0.9 nm, respectively (these being the default values for *Martinize2*). For the OmpC, OmpF, and TolC systems, we specified that the elastic network unit should comprise the entire homotrimer. These elastic network terms are additional harmonic bonds that are added between nonbonded *Martini* beads in order to maintain them at or near the separation distance found in the initial protein model; in doing so, these terms ensure that the protein maintains its native fold throughout the simulation.

In the process of combining the coarse-grained lipids, LPS, and protein, we noticed that the lipid numbers determined by CHARMM-GUI for most of our systems entailed an imbalance in the number of lipid tails in the lower leaflet versus the upper leaflet of the OM. In all cases, the lower leaflet had a slight excess of lipid tails relative to the upper leaflet. In order to generate balanced numbers of lipid tails in the upper and lower leaflets, we implemented a protocol to randomly remove phosphatidylethanolamine and phosphatidylglycerol molecules such that the number of lipid tails in the lower leaflet and upper leaflet would be equal. Then, with the lipids, LPS, and protein combined and the lipid tails balanced, the final stage of our *Martini3* system setup consisted of combining our coarse-grained systems with the original ion atoms from the all-atom CHARMM-GUI system and then solvating with *Martini3* waters. For this latter step, we first used the Python script *Insane* [23] to create a box of *Martini3* waters with the same dimensions as the CHARMM-GUI system and a 40 Å gap for the lipid bilayer. We then implemented the GROMACS [24] solvate utility, specifying a solvent radius of 0.21 nm (the default value for *Martini3* waters).

**Lipid Interaction Analysis**

For determining the lateral solvent-accessible surface area of the membrane-spanning region of each protein, we defined the top of the region by taking the median Z-coordinate of the “B20” beads of LPS, which correspond to an amide and “pseudo-headgroup” for two of the carbon tails of the Lipid A moiety. We defined the bottom of the membrane-spanning region by taking the median Z-coordinate of all the PPPE (phosphatidylethanolamine) headgroup beads (since PPPE is the most abundant of the phospholipid types). Any protein beads within this boundary were deemed “membrane-spanning.” We determined lateral SASA using the Shrake-Rupley algorithm [25] with a probe radius of 5 Å. Determination of lateral-facing residues was done by a radius-based approach, wherein each protein was approximated as a cylinder, and the residues were divided into angular sectors of 10 degrees. Within each sector, we searched for the maximum radius and any atoms within 70% of the maximum radius. These residues were marked as “outer” residues and were the only residues used for SASA determination. We did this to avoid counting interior-facing residues toward the SASA as much as possible, since these residues are not capable of interacting with lipids.

| Protein | Copy Number | Percent Copy Number | Cumulative Percentage |
| --- | --- | --- | --- |
| OmpA | 207618 | 29.55% | 29.55% |
| OmpC | 163538 | 23.28% | 52.83% |
| OmpX | 125295 | 17.83% | 70.66% |
| OmpF | 88988 | 12.67% | 83.33% |
| OmpT | 40237 | 5.73% | 89.05% |
| MipA | 20925 | 2.98% | 92.03% |
| Tsx | 14911 | 2.12% | 94.15% |
| TolC | 8768 | 1.25% | 95.40% |
| FadL | 6912 | 0.98% | 96.38% |
| YdiY | 5888 | 0.84% | 97.22% |

Table S1: Ribosome profiling data [18] shows that the distribution of protein copy numbers is heavily skewed toward the top 10 most-expressed proteins.

| **OMPF** | **Interface** | **piTM** | **Average** |
| --- | --- | --- | --- |
| model_1 | 0.0142 | 0.119 | 0.0666 |
| model_2 | 0.902 | 0.9139 | 0.90795 |
| model_3 | 0.9009 | 0.9074 | 0.90415 |
| model_4 | 0.8944 | 0.9022 | 0.8983 |
| model_5 | 0.8949 | 0.9041 | 0.8995 |
| **OMPC** | **Interface** | **piTM** | **Average** |
| model_1 | 0.0147 | 0.1572 | 0.08595 |
| model_2 | 0.8722 | 0.8767 | 0.87445 |
| model_3 | 0.8639 | 0.8502 | 0.85705 |
| model_4 | 0.8736 | 0.8697 | 0.87165 |
| model_5 | 0.8789 | 0.8642 | 0.87155 |
| **TOLC** | **Interface** | **piTM** | **Average** |
| model_1 | 0.7434 | 0.7295 | 0.73645 |
| model_2 | 0.7242 | 0.7148 | 0.7195 |
| model_3 | 0.6805 | 0.6603 | 0.6704 |
| model_4 | 0.7556 | 0.7318 | 0.7437 |
| model_5 | 0.7785 | 0.7643 | 0.7714 |

Table S2: AF2 Complex confidence scores for the 3 homotrimer OM proteins. Five models were generated, with the highlighted model being the one that was used for simulation, based on having the highest average of the Interface and piTM score (both piTM and Interface are confidence scores that range from zero to one).

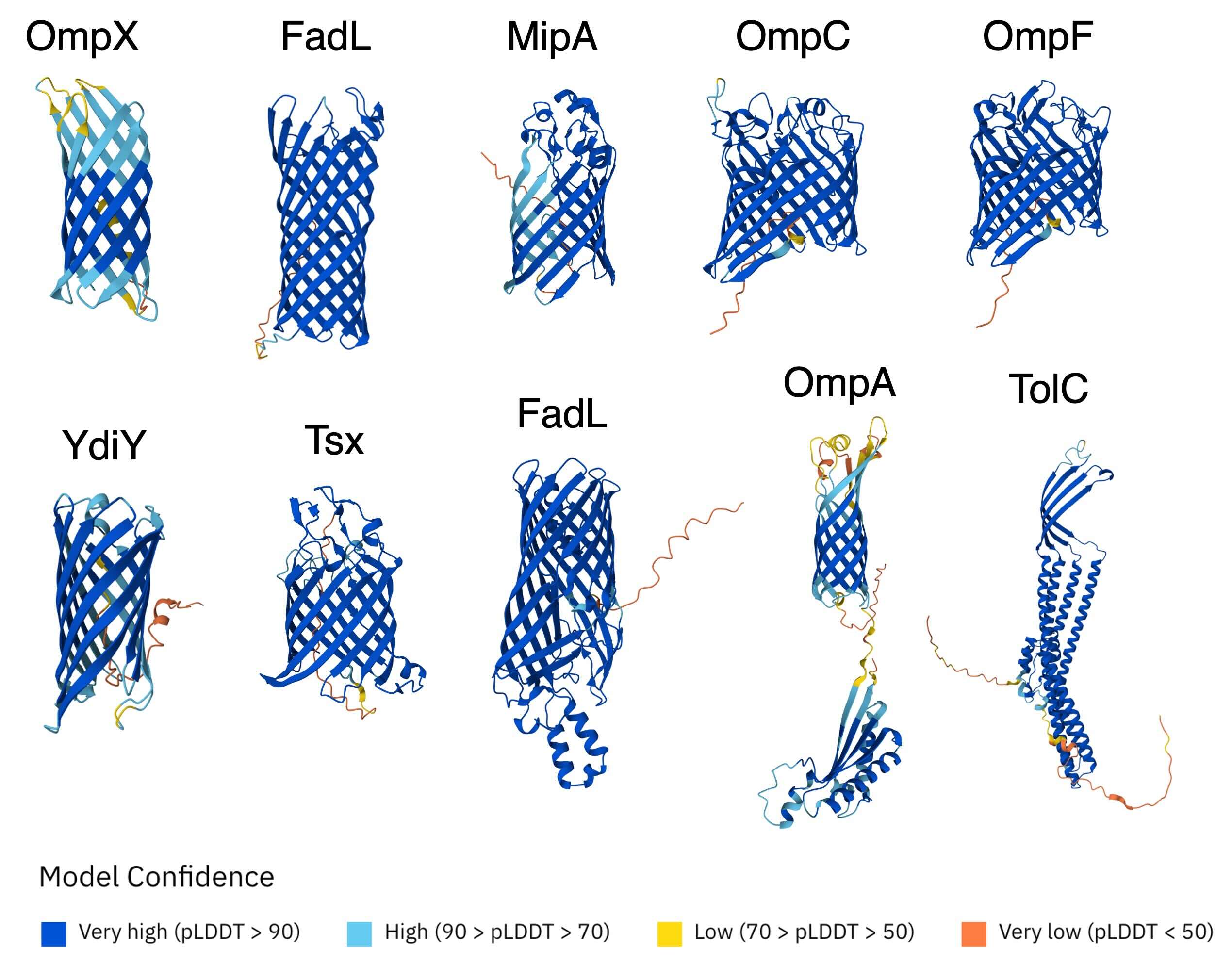

Figure S1: AlphaFold2 models with residues colored according to their pLDDT confidence level. All of my models exhibited high confidence across their core structure. Signal peptides (which I removed) and various loops showed lower confidence. Note that only the monomeric protein is shown for the homotrimers—OmpC, OmpF, and TolC were also evaluated according to the complex predictor AF2-Complex.

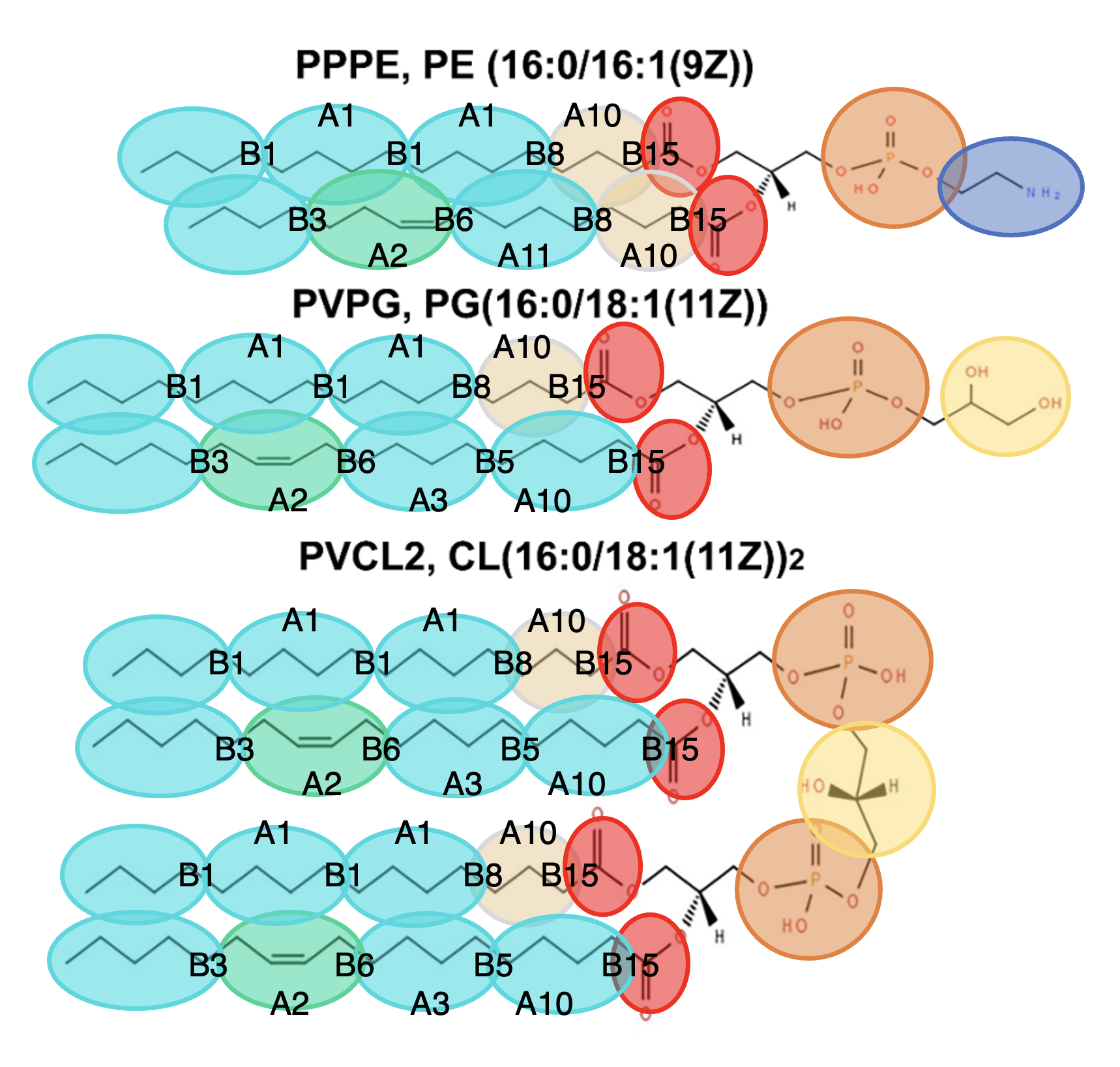

Figure S2: Illustration of our coarse-graining scheme for the 3 lipids from the inner leaflet of the E. coli outer membrane. We were guided by SwarmCG’s method of using different-sized Martini3 carbon beads to differentiate aliphatic chains of length 16C vs 18C. Our bond and angle nomenclature is also borrowed directly from SwarmCG.

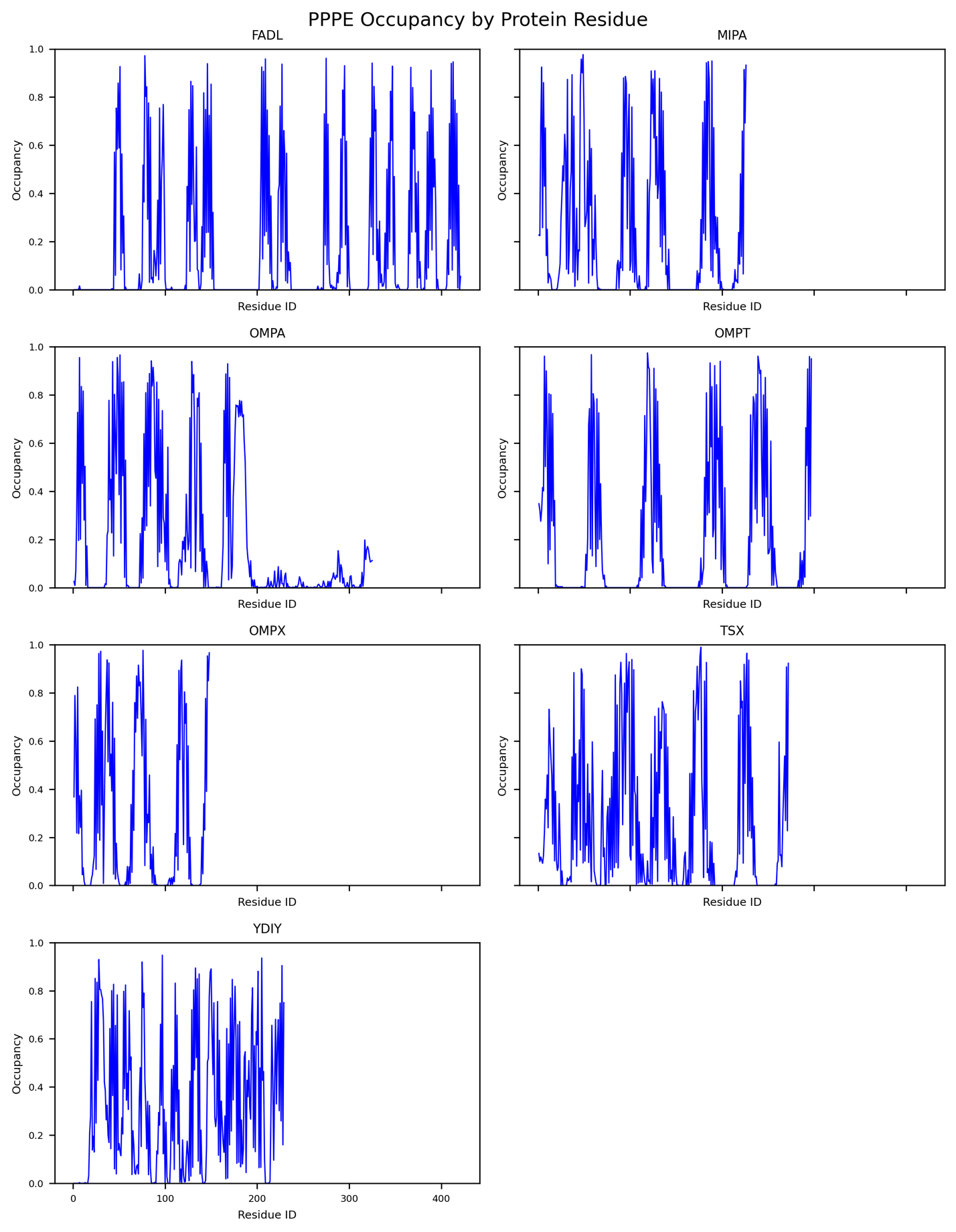

Figure S3: PPPE occupancy levels for my monomer OMP simulations with O-antigen chain length 0

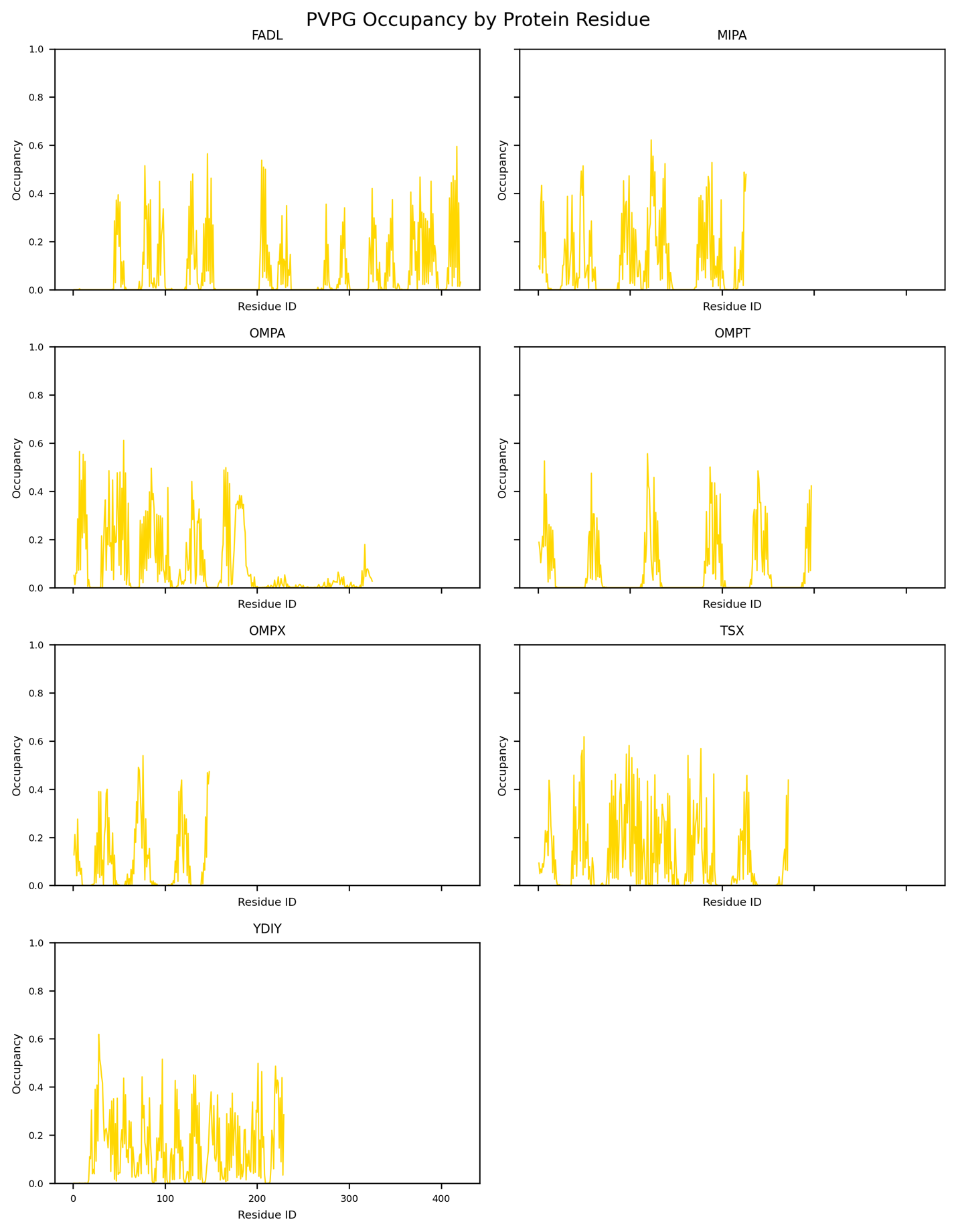

Figure S4: PVPG occupancy levels for my monomer OMP simulations with O-antigen chain length 0

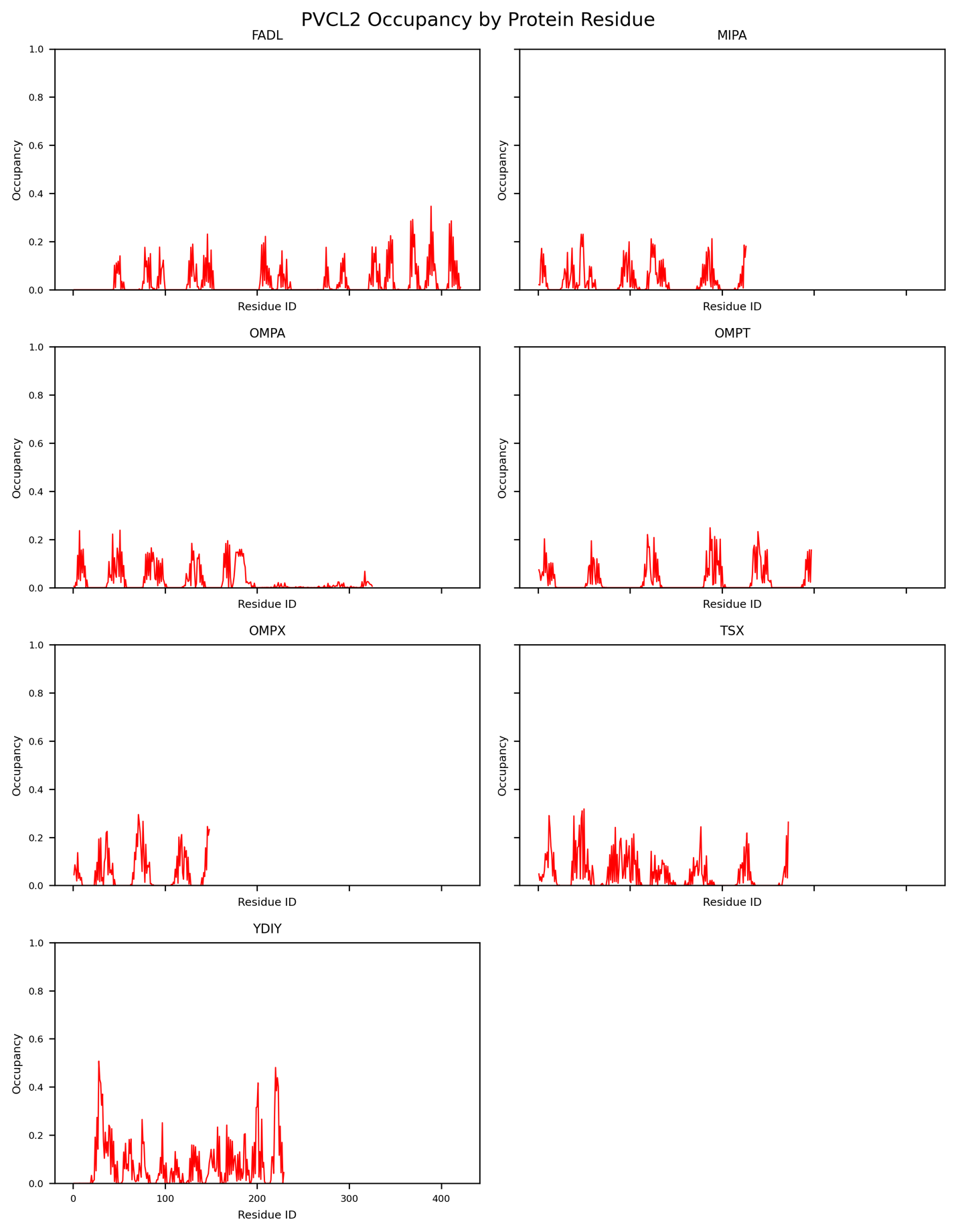

Figure S5: PVCL2 occupancy levels for my monomer OMP simulations with O-antigen chain length 0

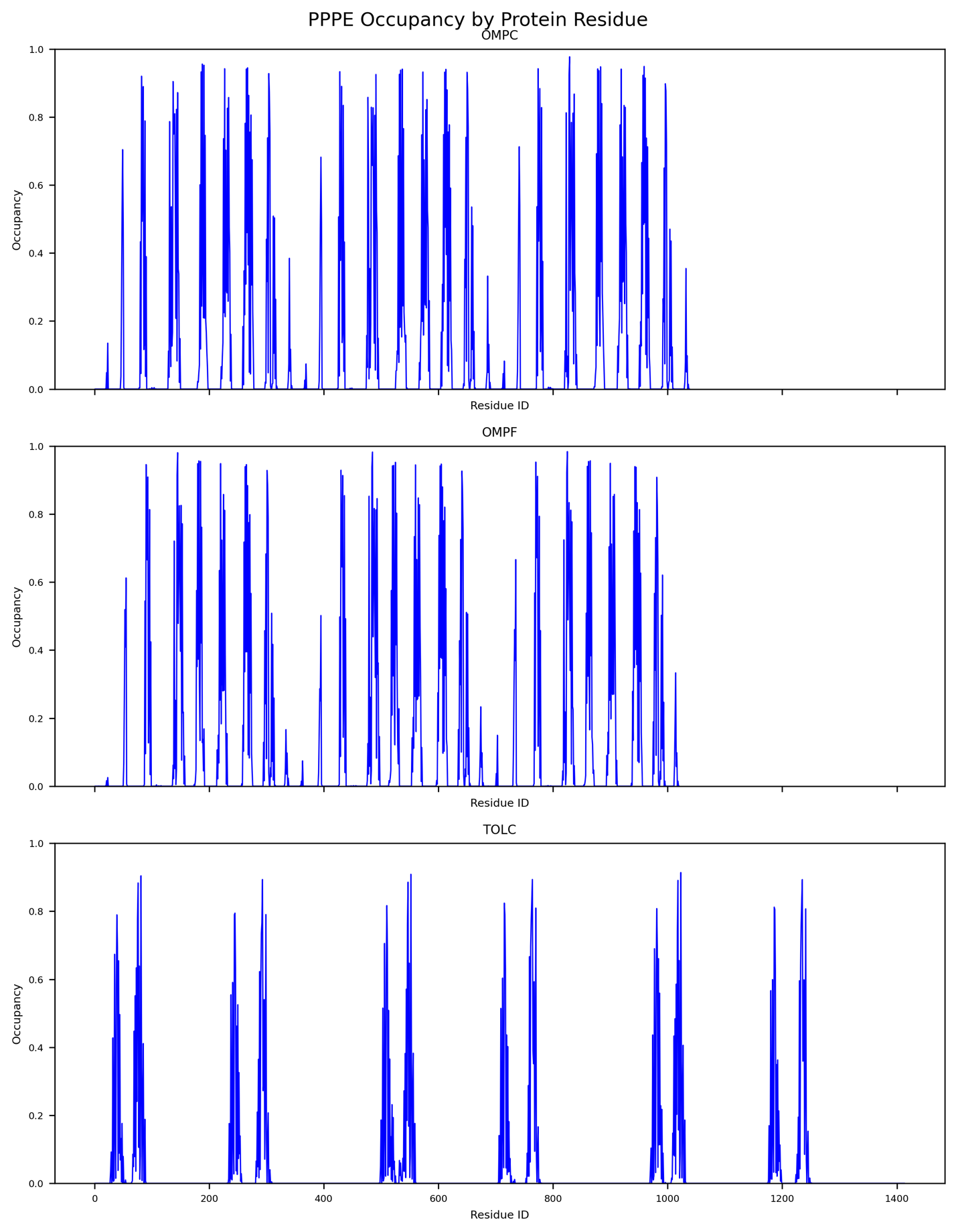

Figure S6: PPPE occupancy levels for my trimer OMP simulations with O-antigen chain length 0

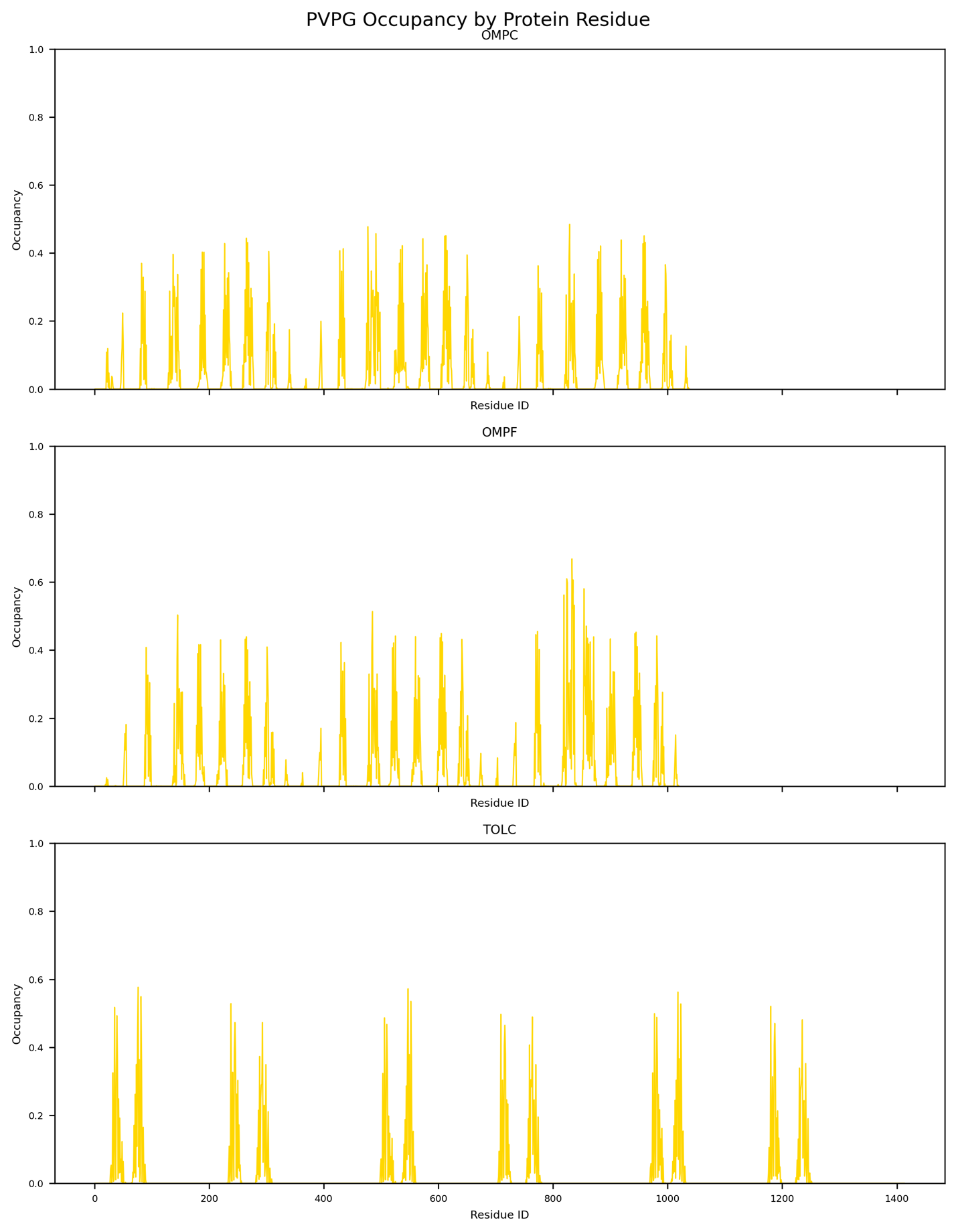

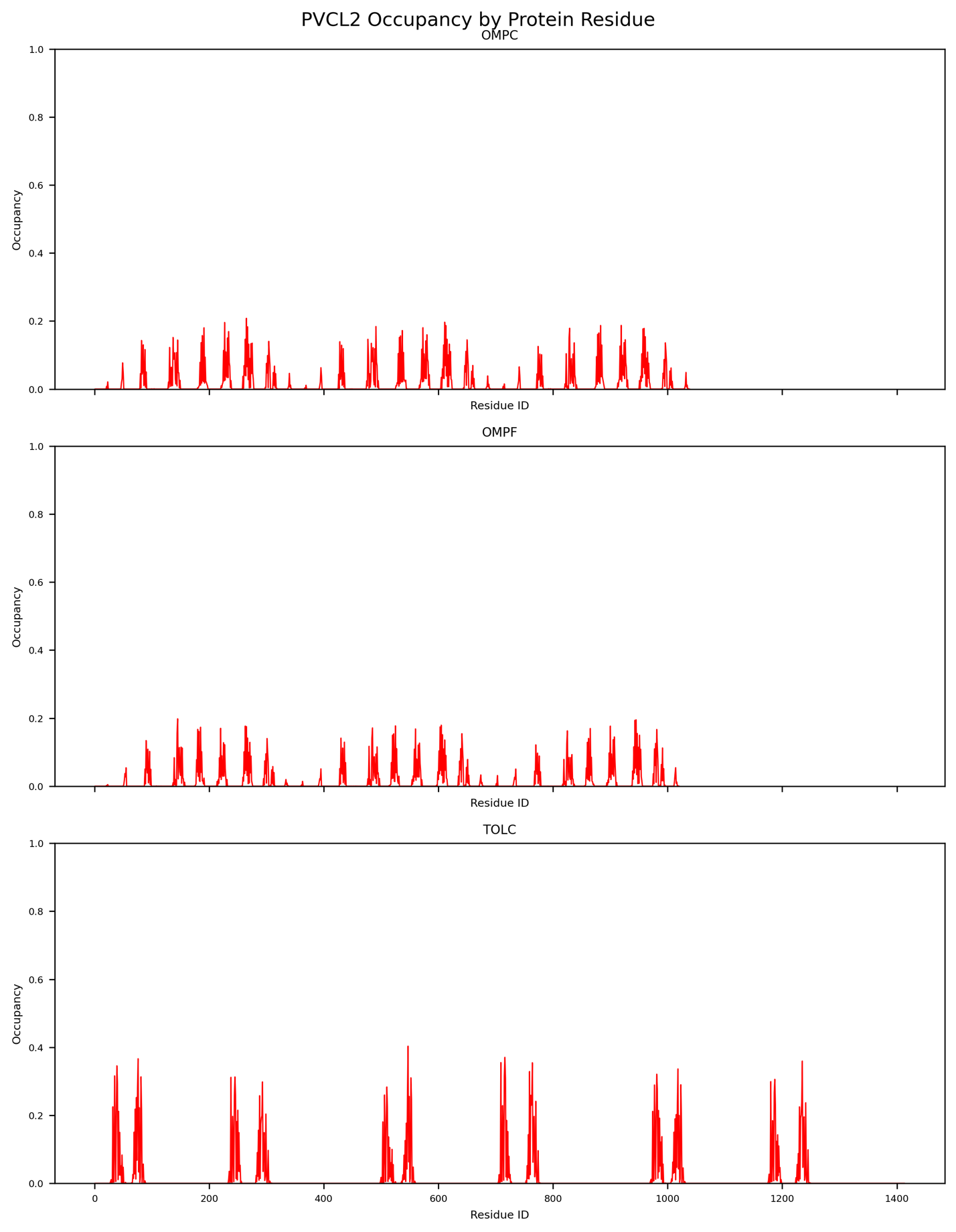

Figure S7: PVPG occupancy levels for my trimer OMP simulations with O-antigen chain length 0

Figure S8: PVCL2 occupancy levels for my trimer OMP simulations with O-antigen chain length 0

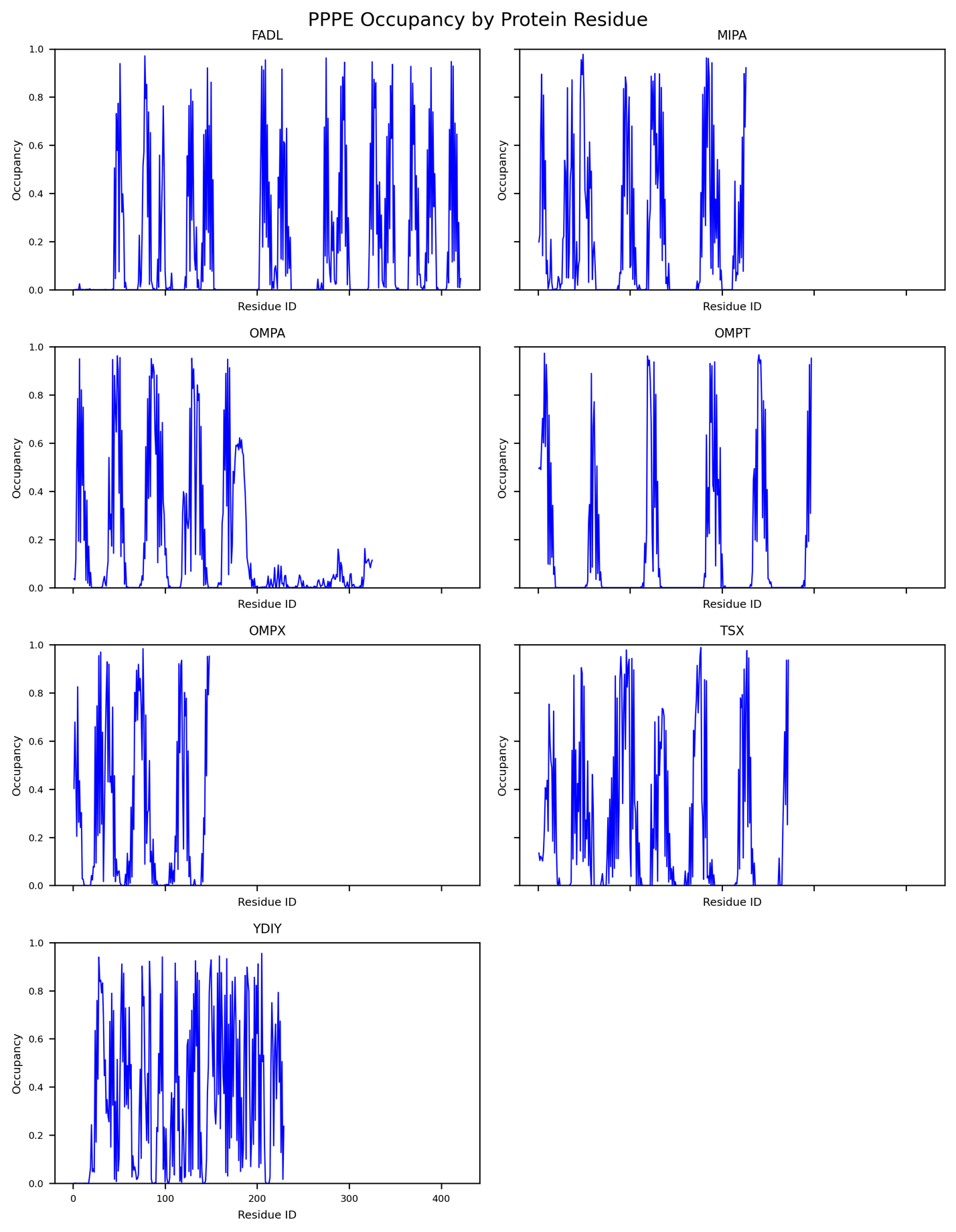

Figure S9: PPPE occupancy levels for my monomer OMP simulations with O-antigen chain length 2

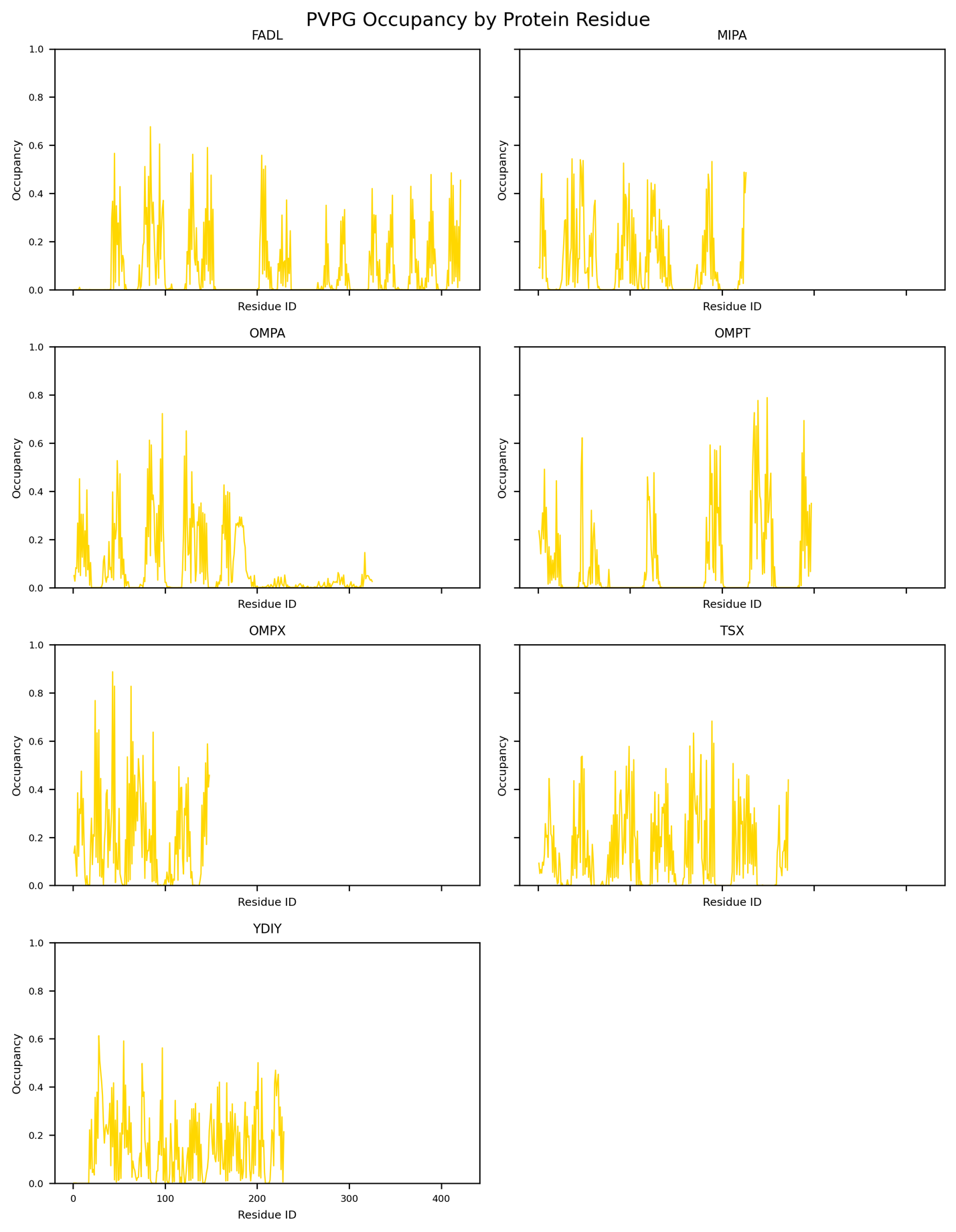

Figure S10: PVPG occupancy levels for my monomer OMP simulations with O-antigen chain length 2

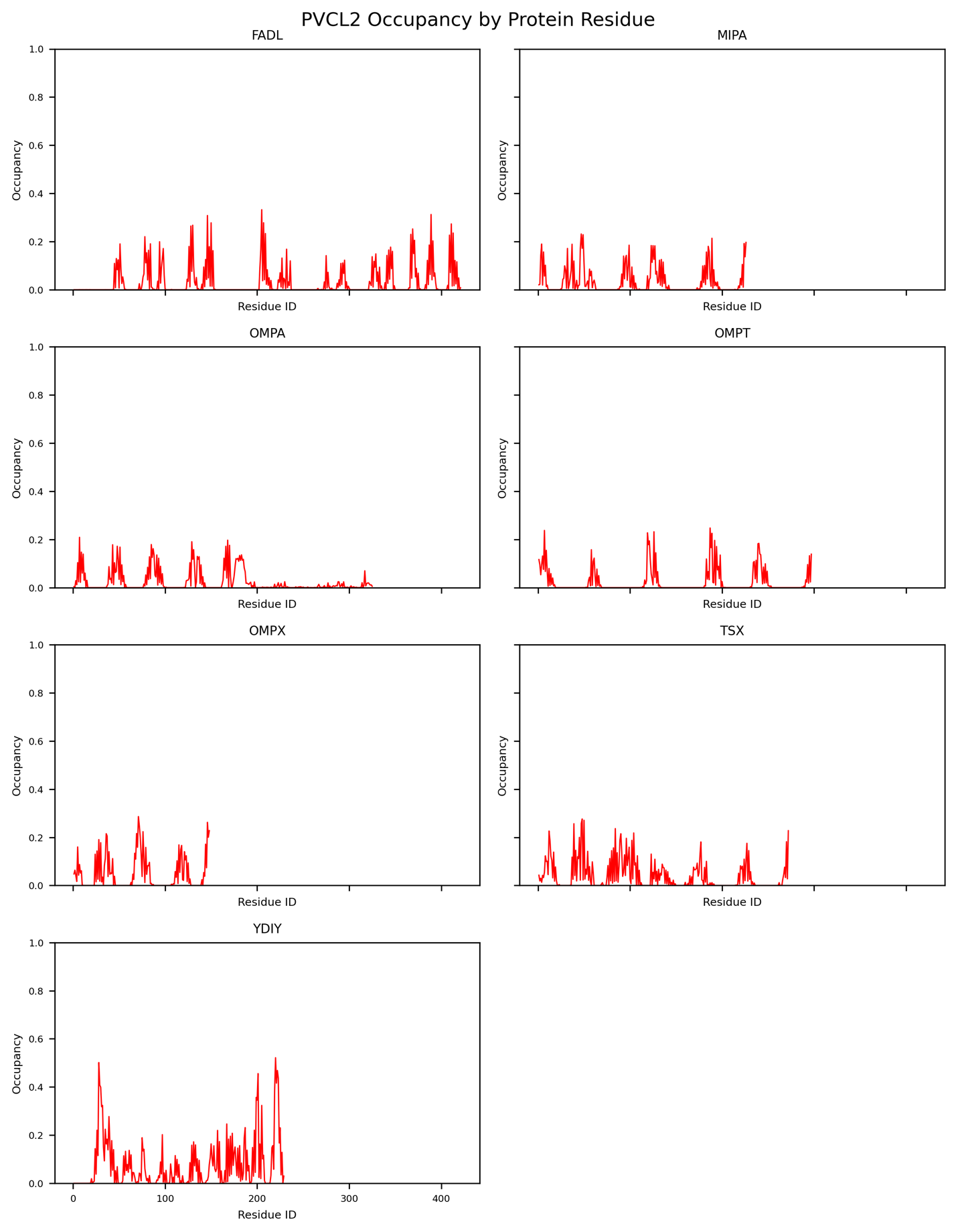

Figure S11: PVCL2 occupancy levels for my monomer OMP simulations with O-antigen chain length 2

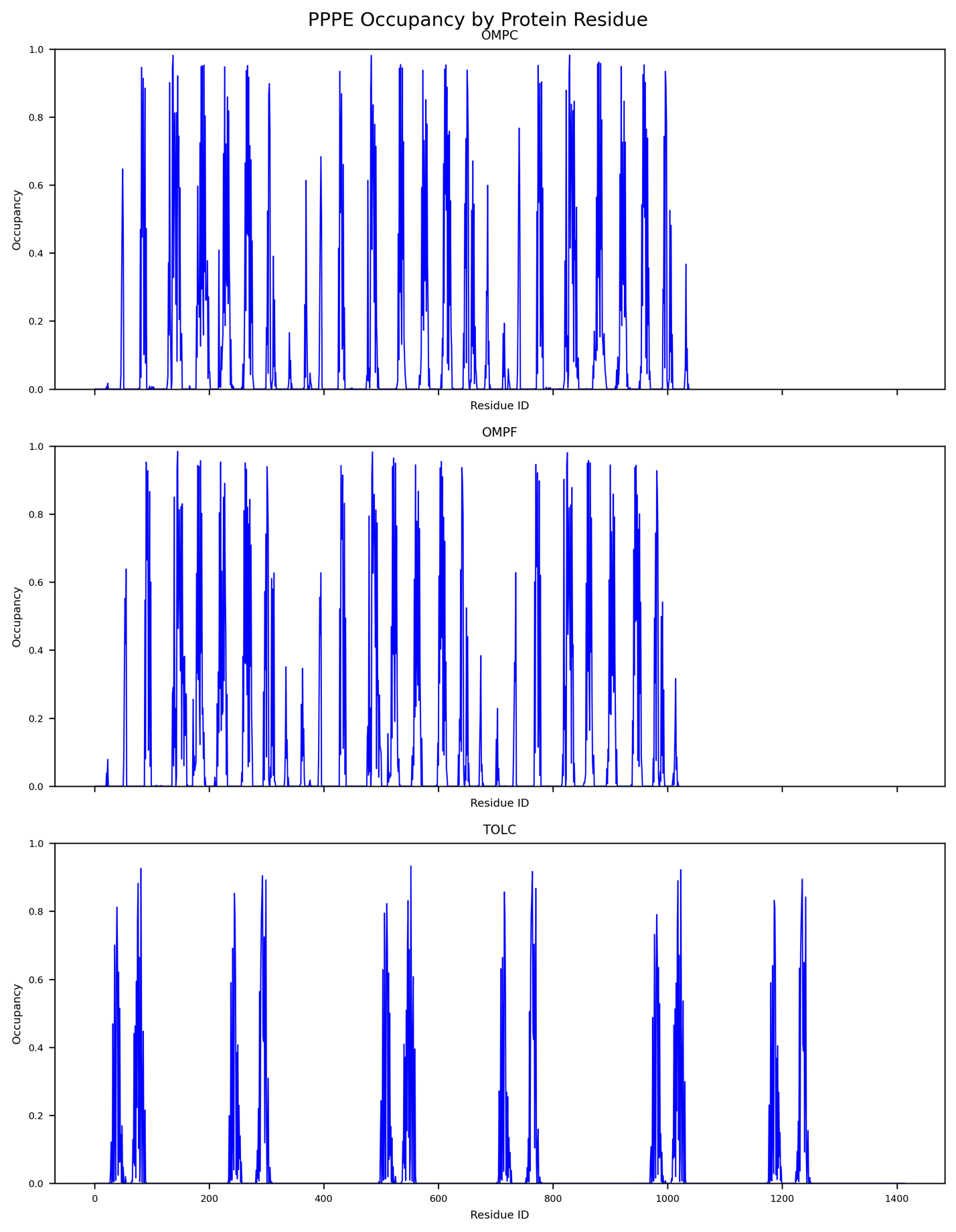

Figure S12: PPPE occupancy levels for my trimer OMP simulations with O-antigen chain length 2

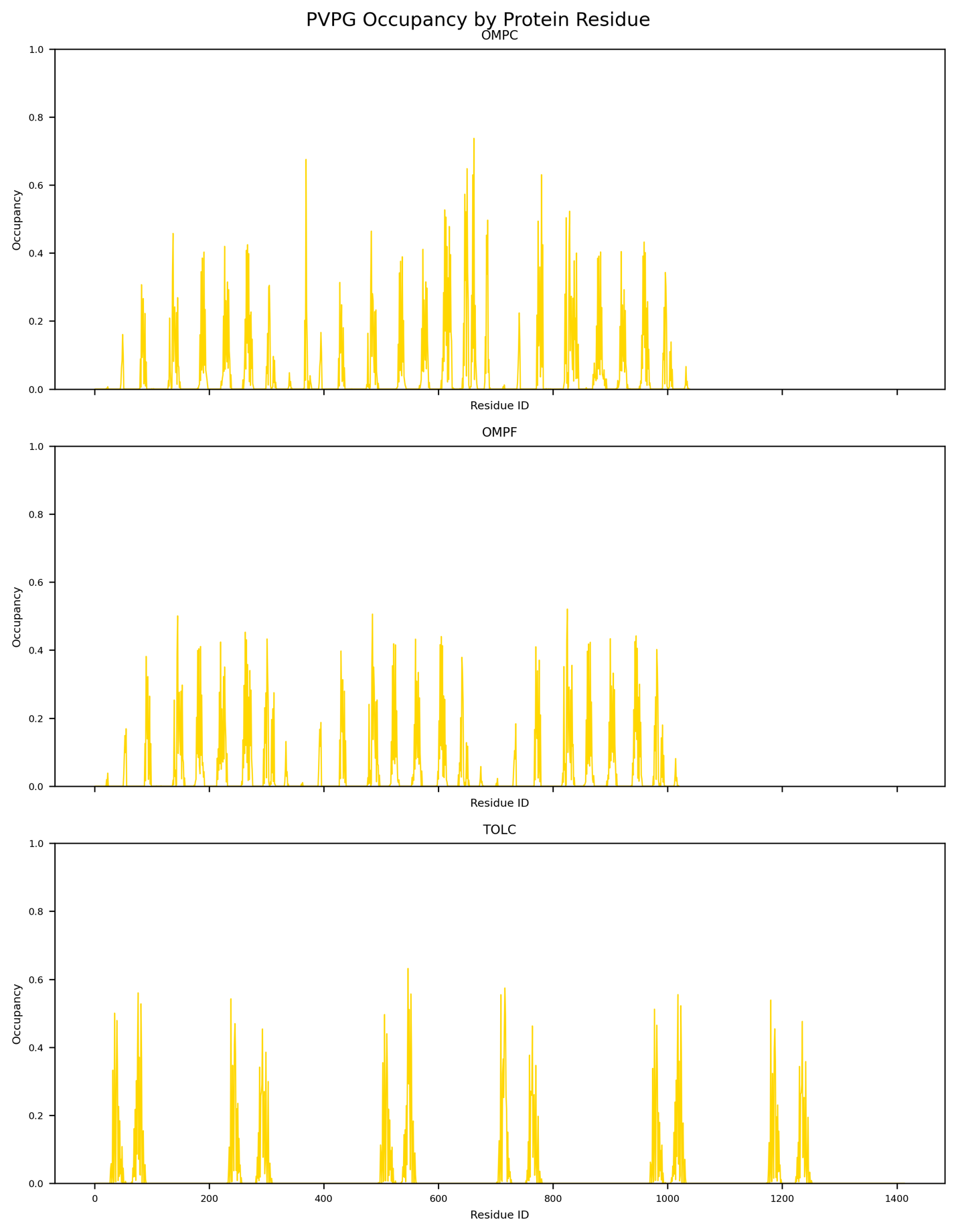

Figure S13: PVPG occupancy levels for my trimer OMP simulations with O-antigen chain length 2

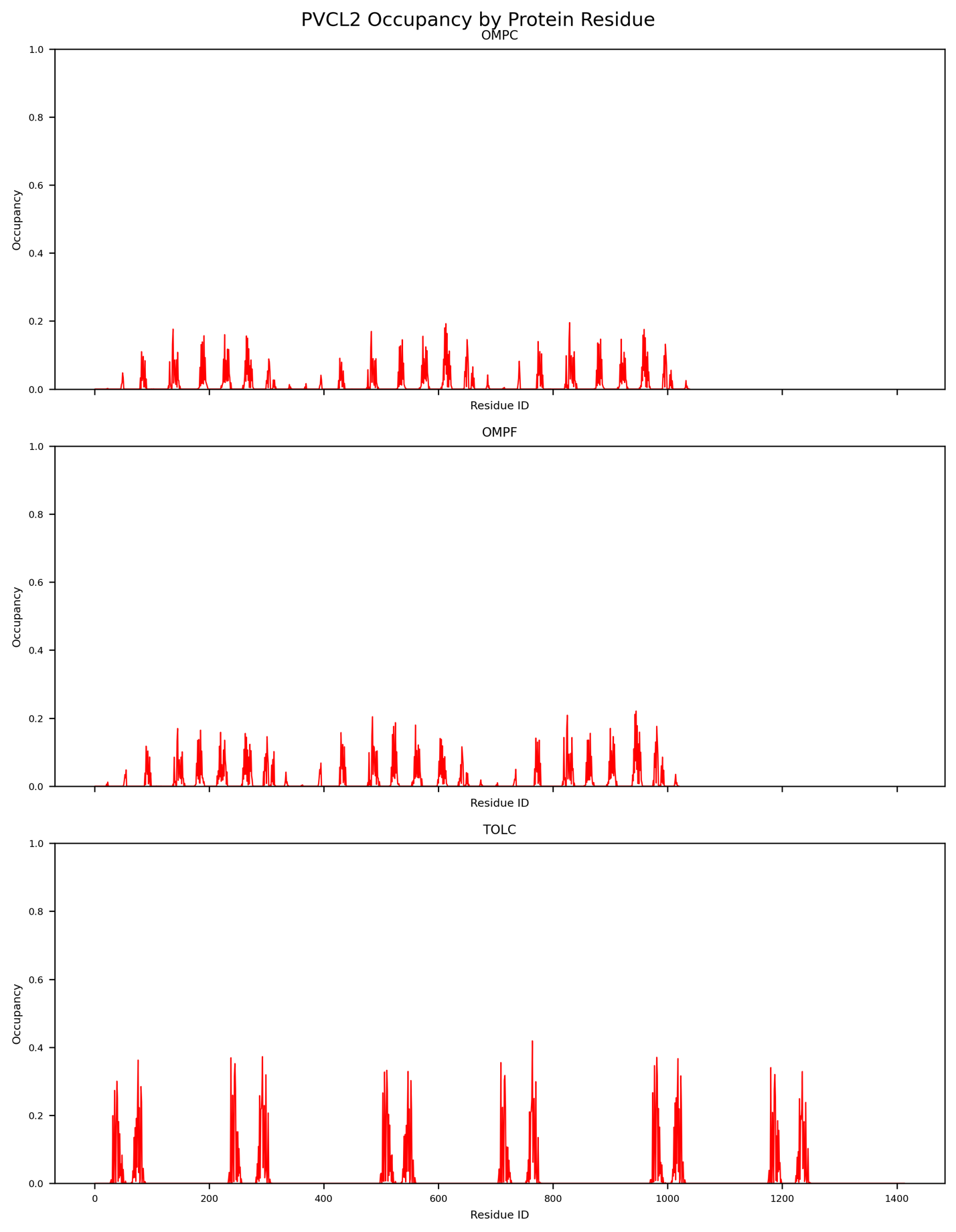

Figure S14: PVCL2 occupancy levels for my trimer OMP simulations with O-antigen chain length 2

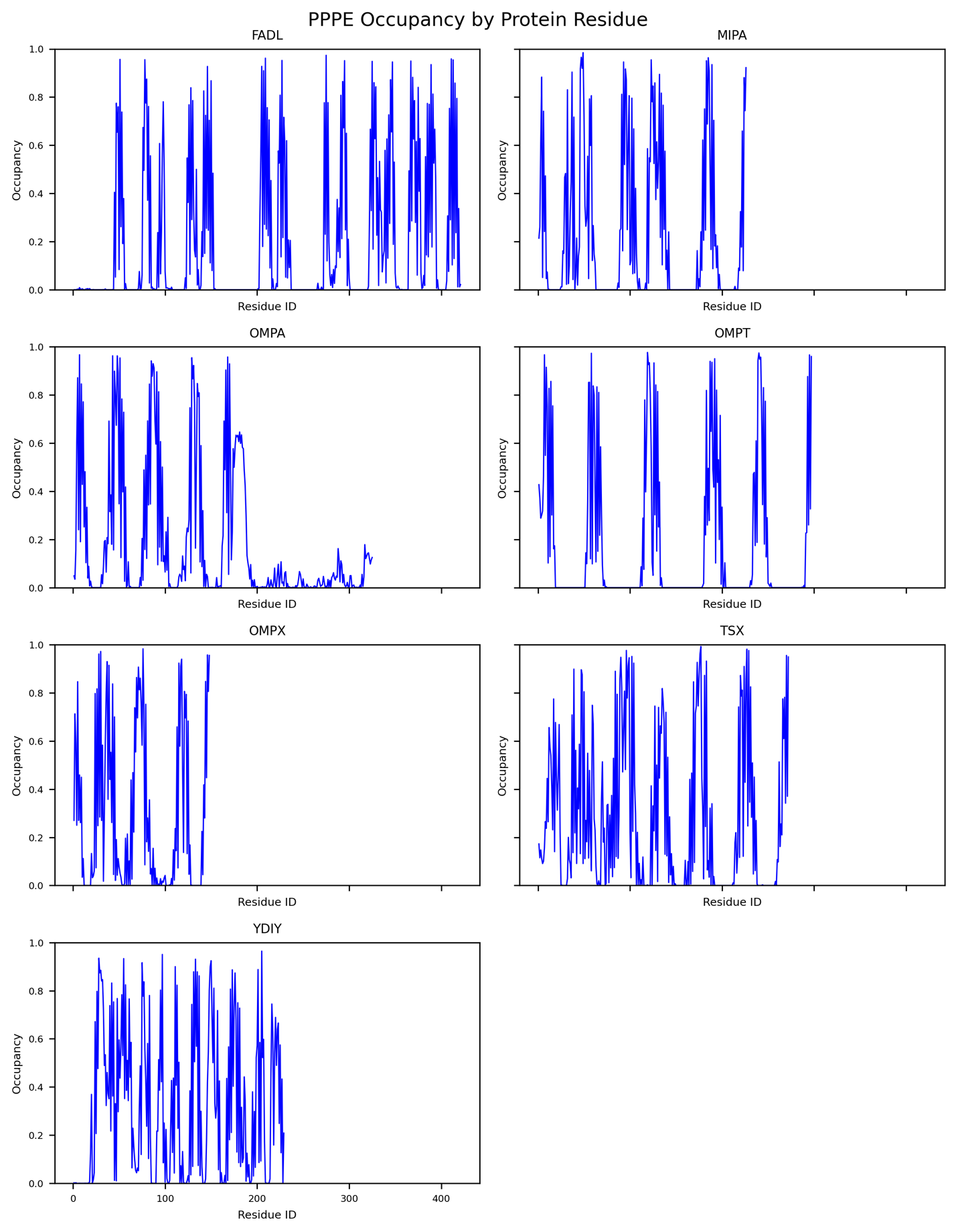

Figure S15: PPPE occupancy levels for my monomer OMP simulations with O-antigen chain length 5

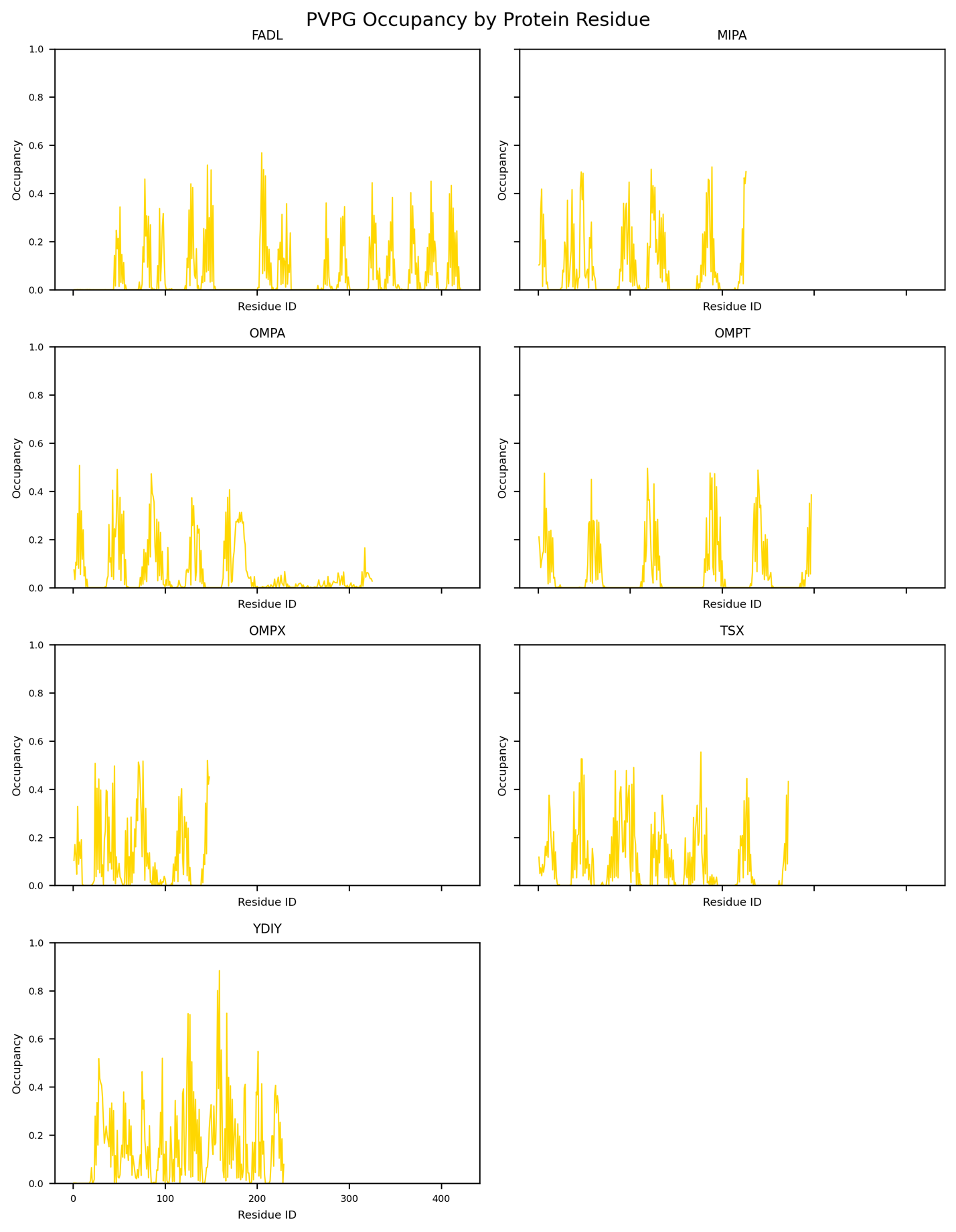

Figure S16: PVPG occupancy levels for my monomer OMP simulations with O-antigen chain length 5

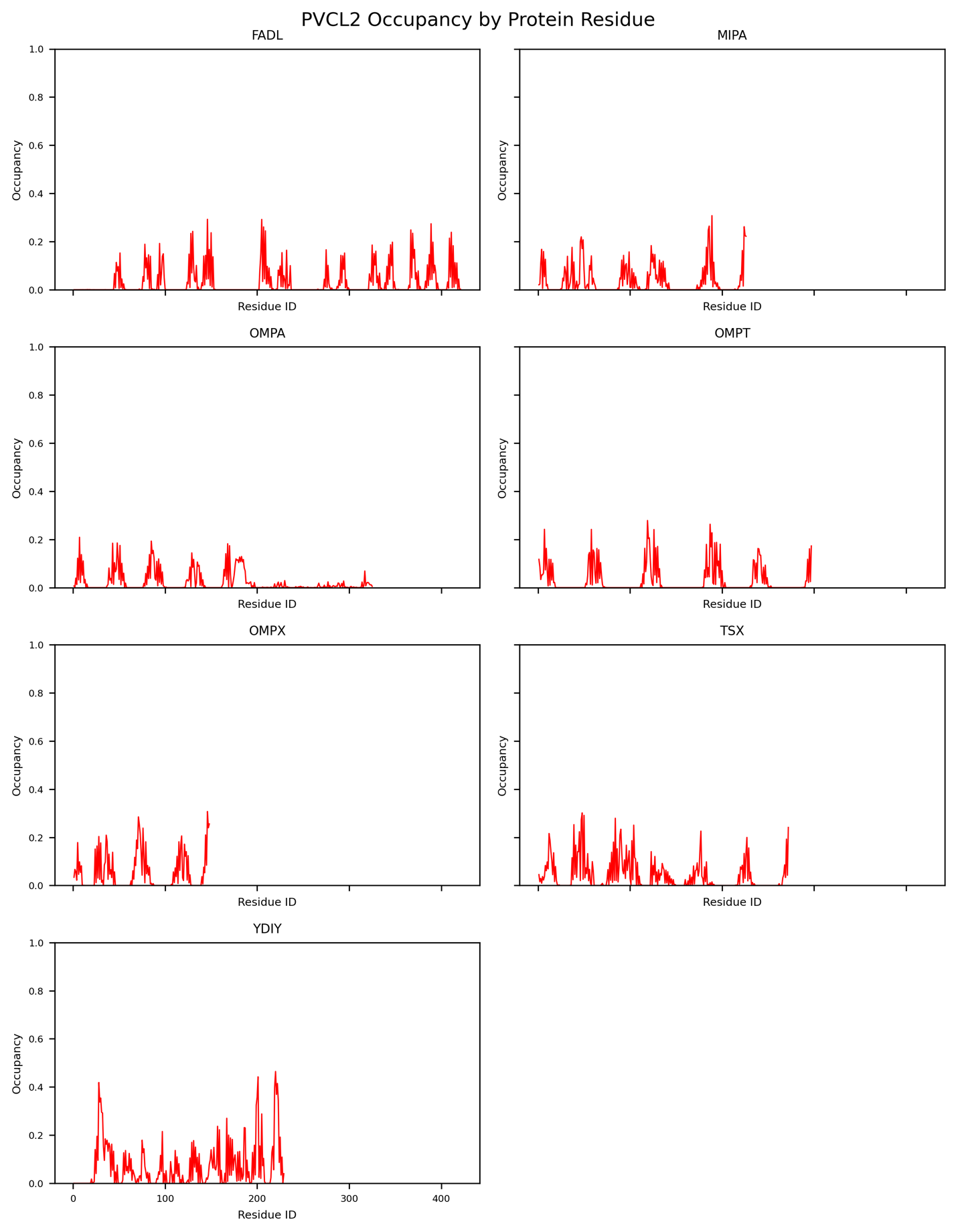

Figure S17: PVCL2 occupancy levels for my monomer OMP simulations with O-antigen chain length 5

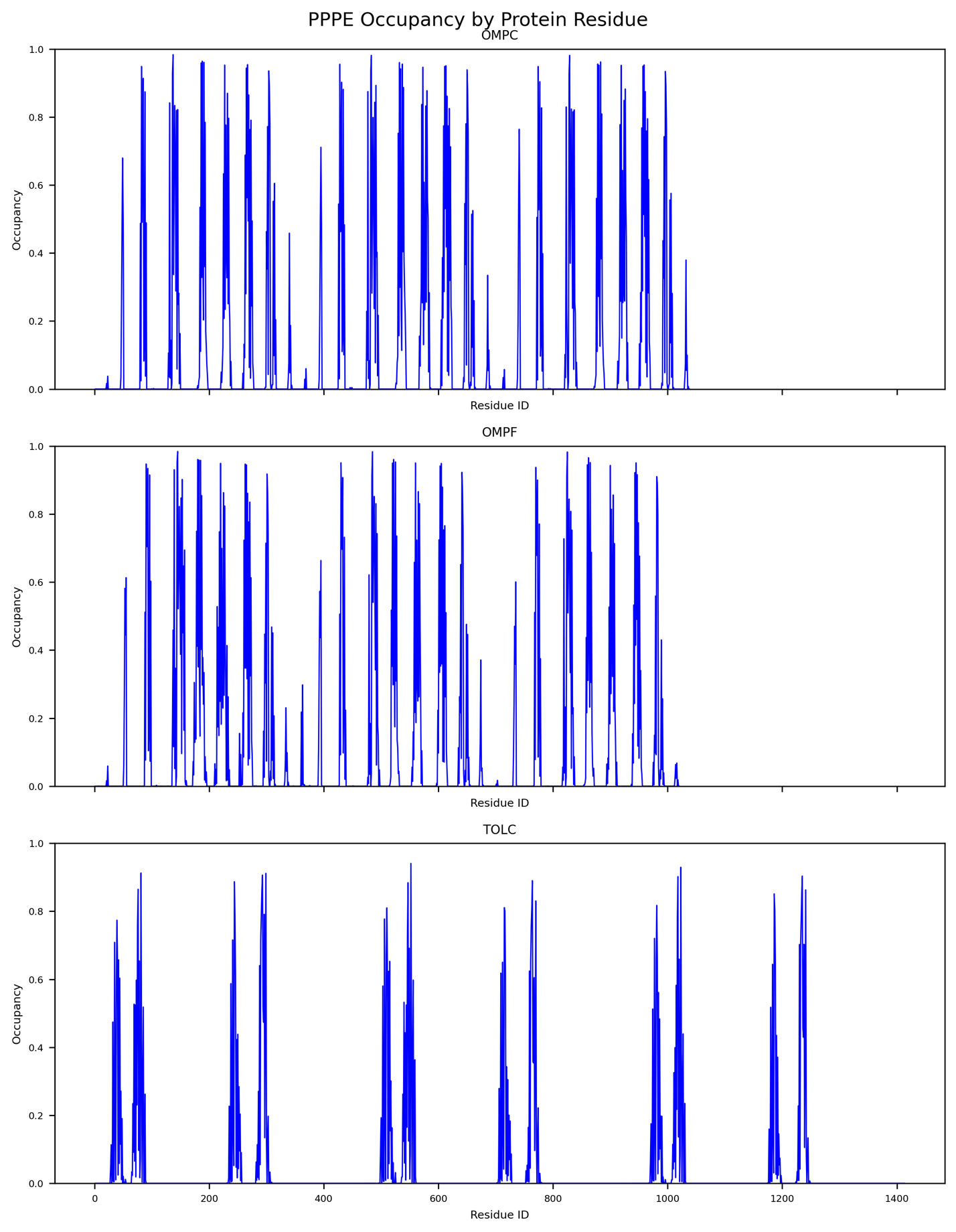

Figure S18: PPPE occupancy levels for my trimer OMP simulations with O-antigen chain length 5

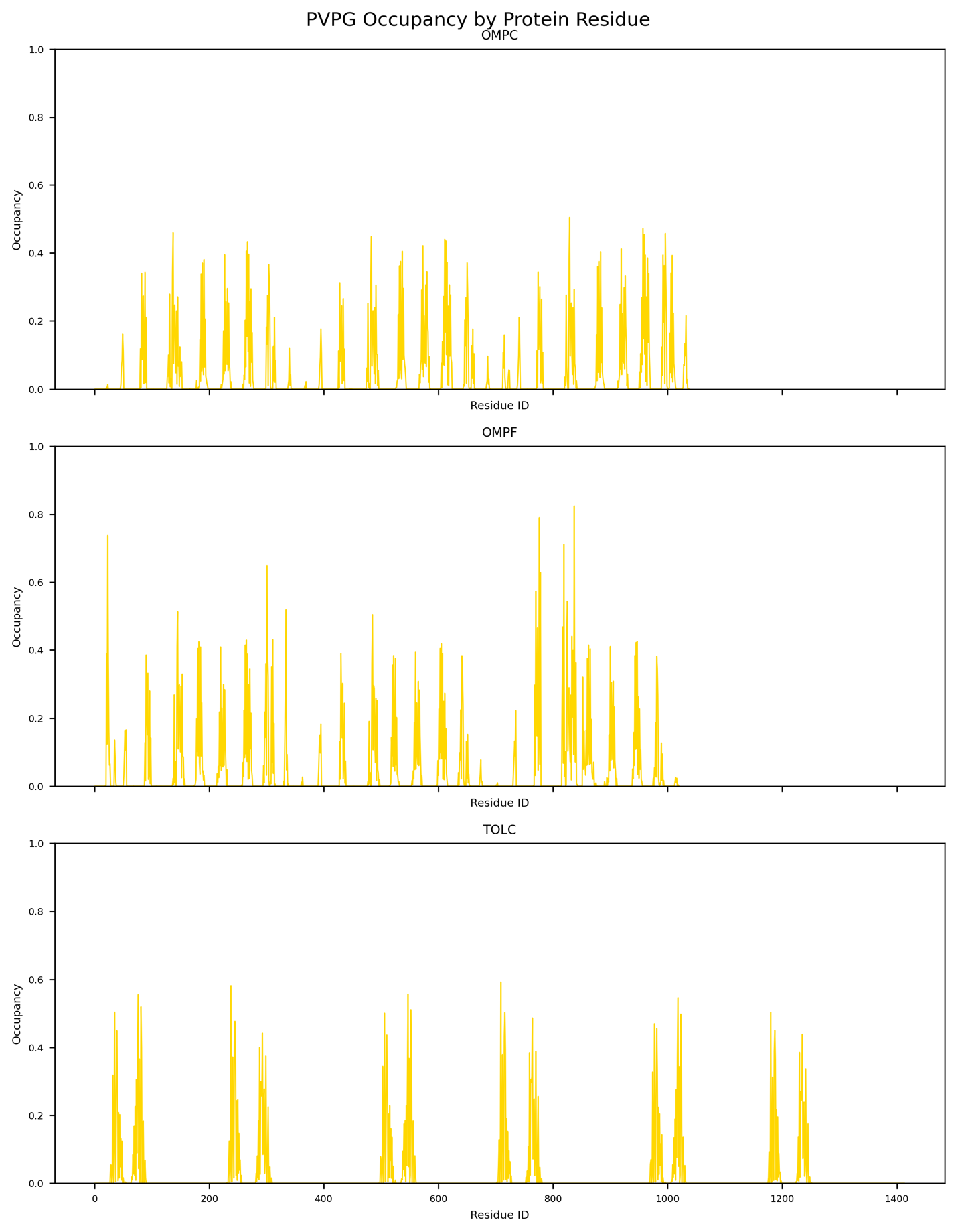

Figure S19: PVPG occupancy levels for my trimer OMP simulations with O-antigen chain length 5

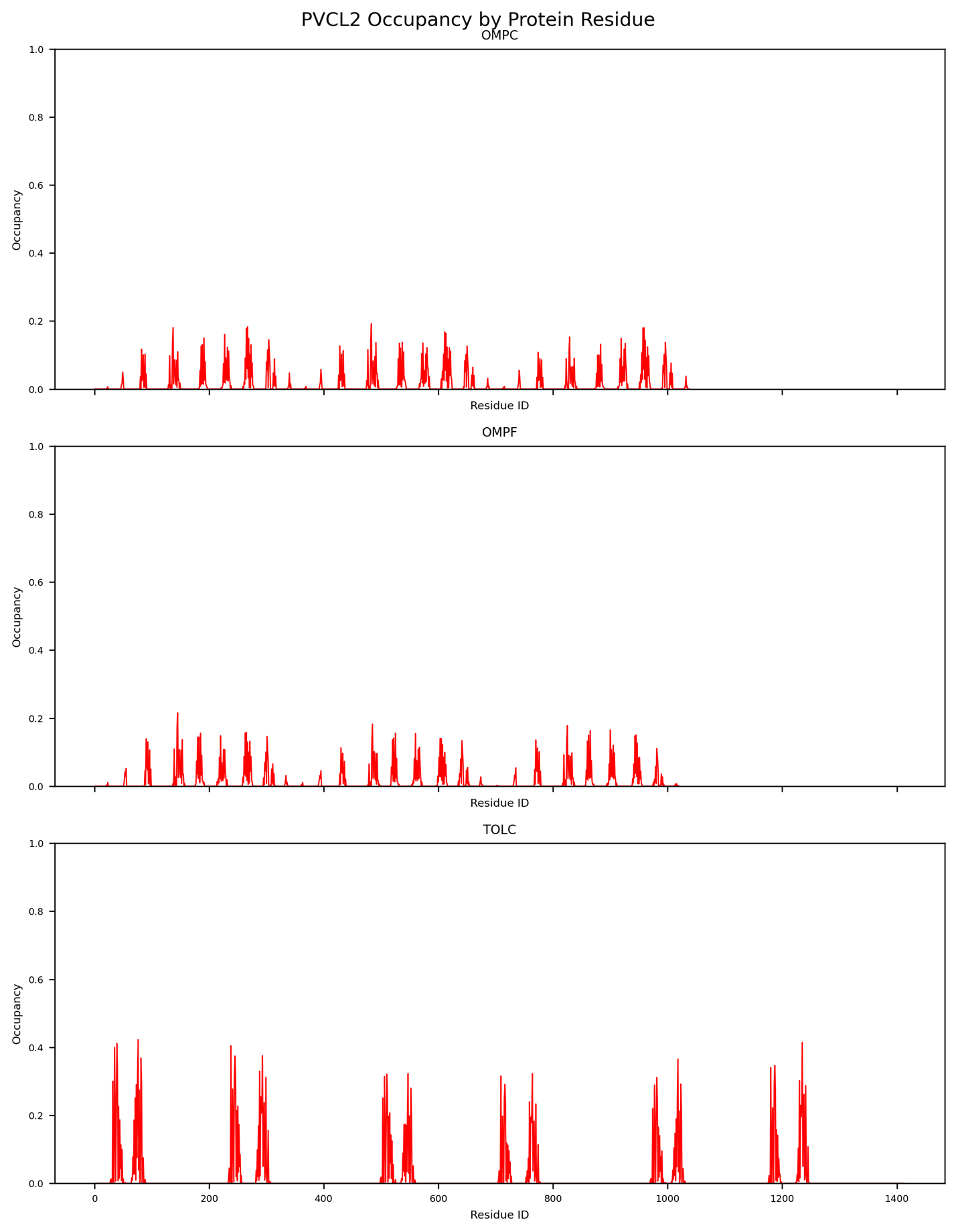

Figure S20: PVCL2 occupancy levels for my trimer OMP simulations with O-antigen chain length 5

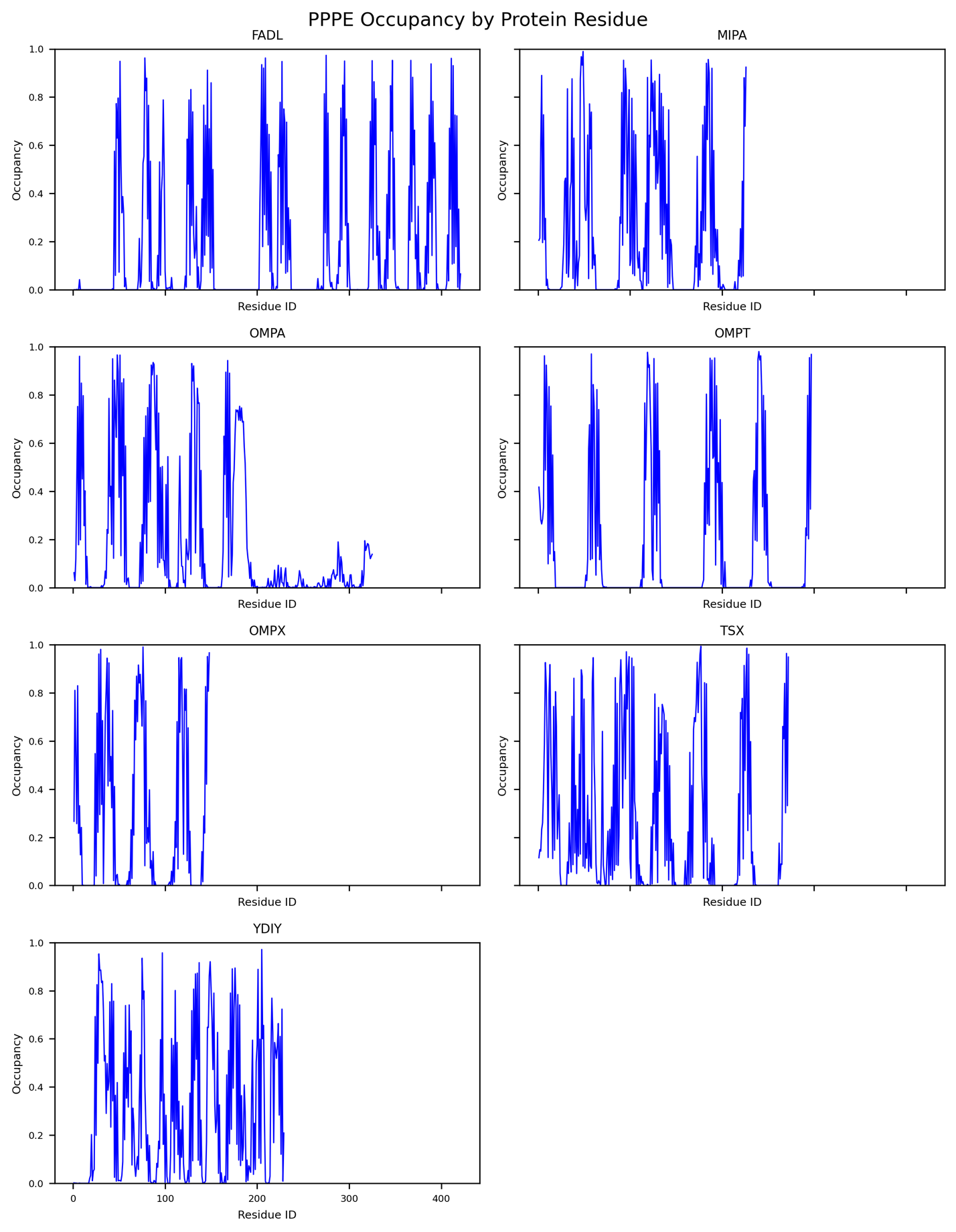

Figure S21: PPPE occupancy data for the 7 monomeric protein systems with LPS O-antigen chain length 10

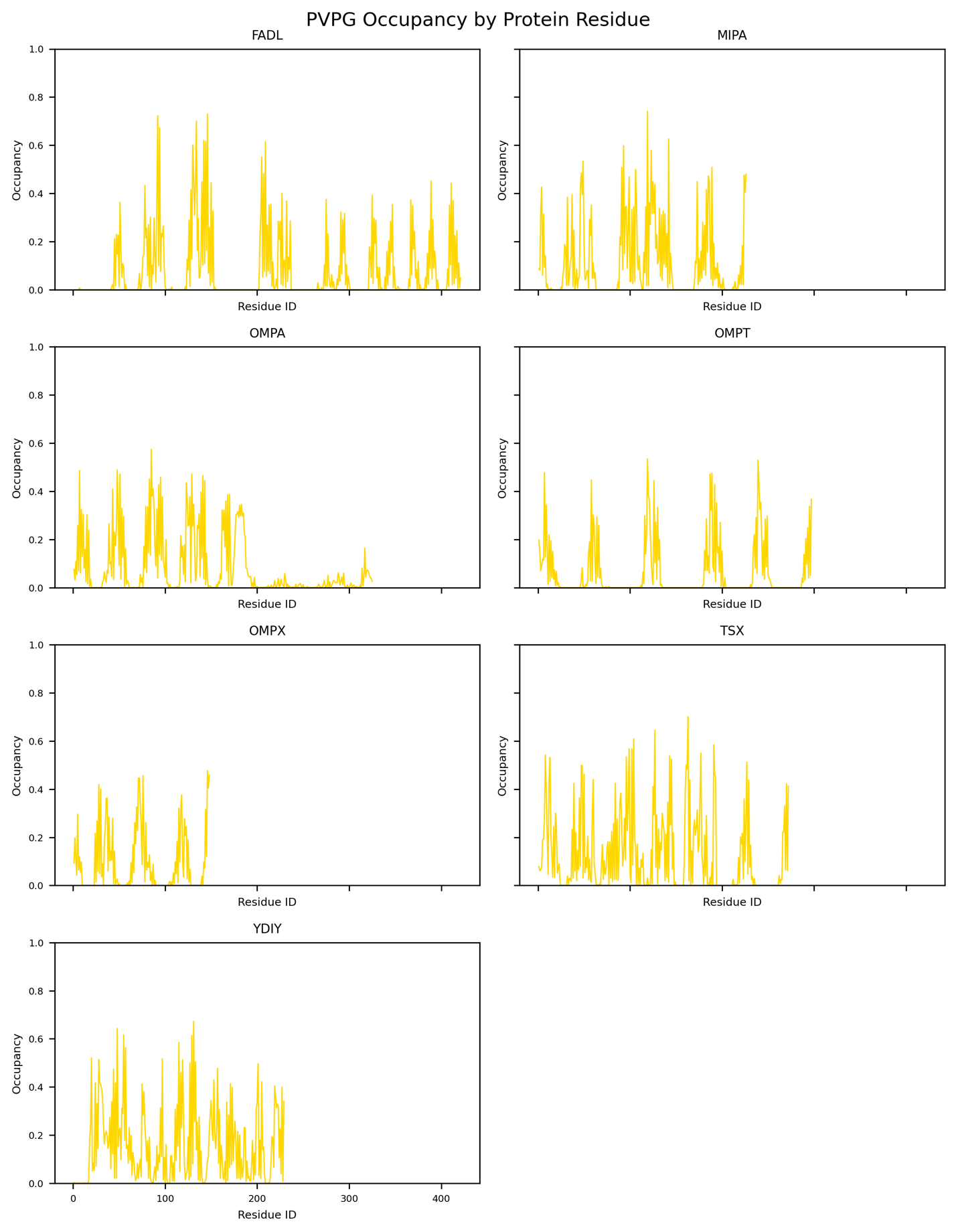

Figure S22: PVPG occupancy data for the 7 monomeric protein systems with LPS O-antigen chain length 10 (note that shorter proteins do not use the entire X-axis length).

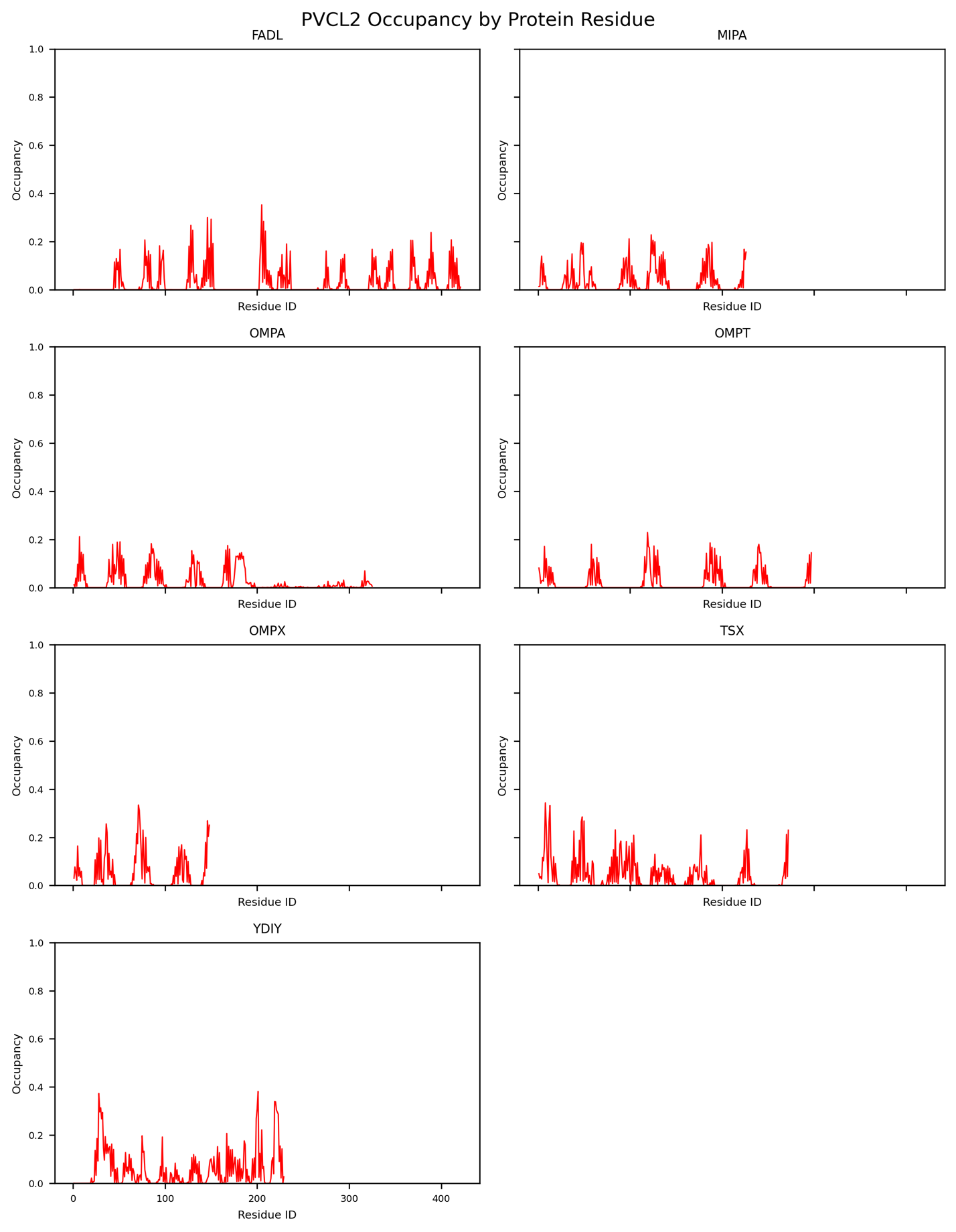

Figure S23: PVCL2 occupancy data for the 7 monomeric protein systems with LPS O-antigen chain length 10 (note that shorter proteins do not use the entire X-axis length).

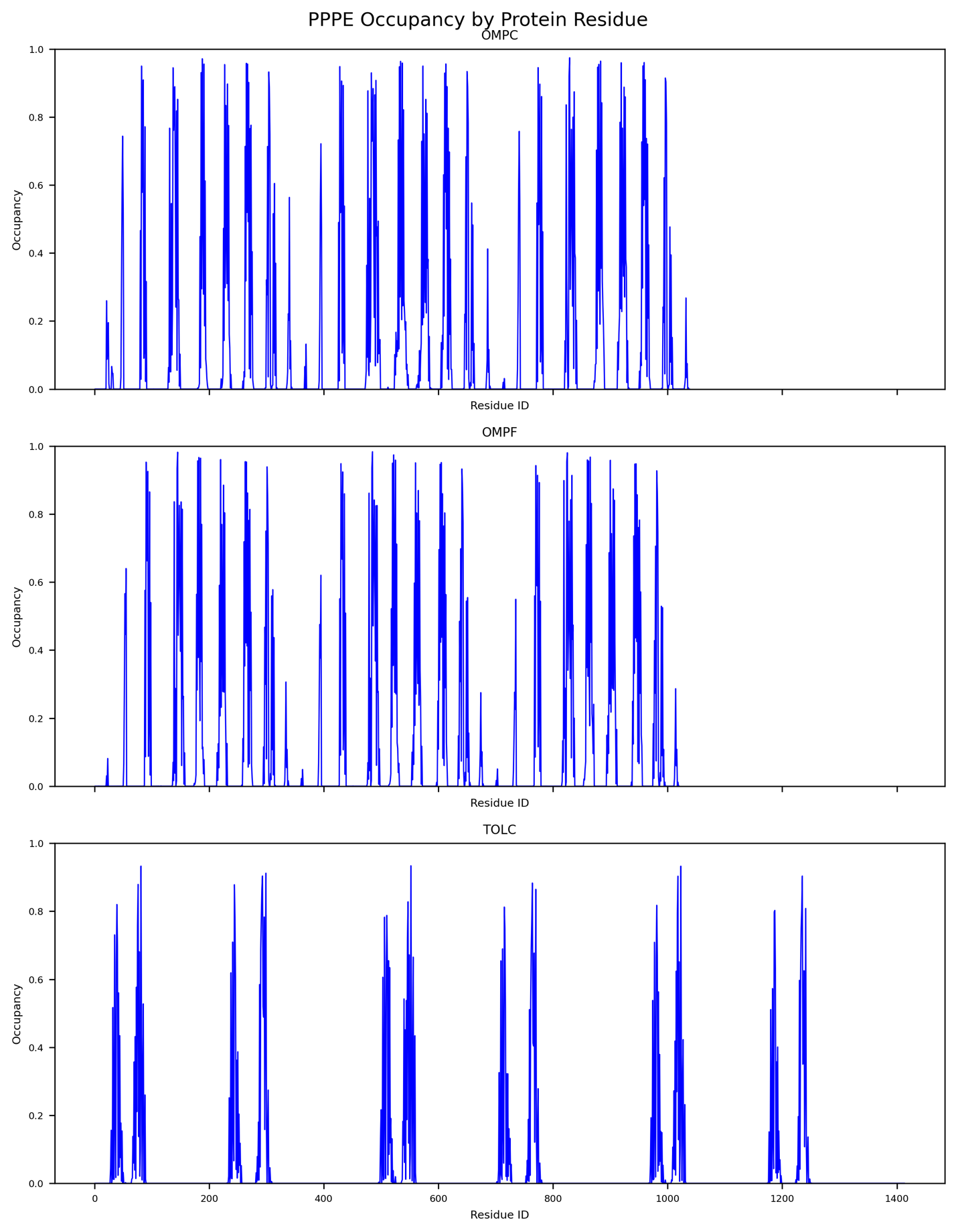

Figure S24: PPPE occupancy data for the 3 trimer protein systems with LPS O-antigen chain length 10.

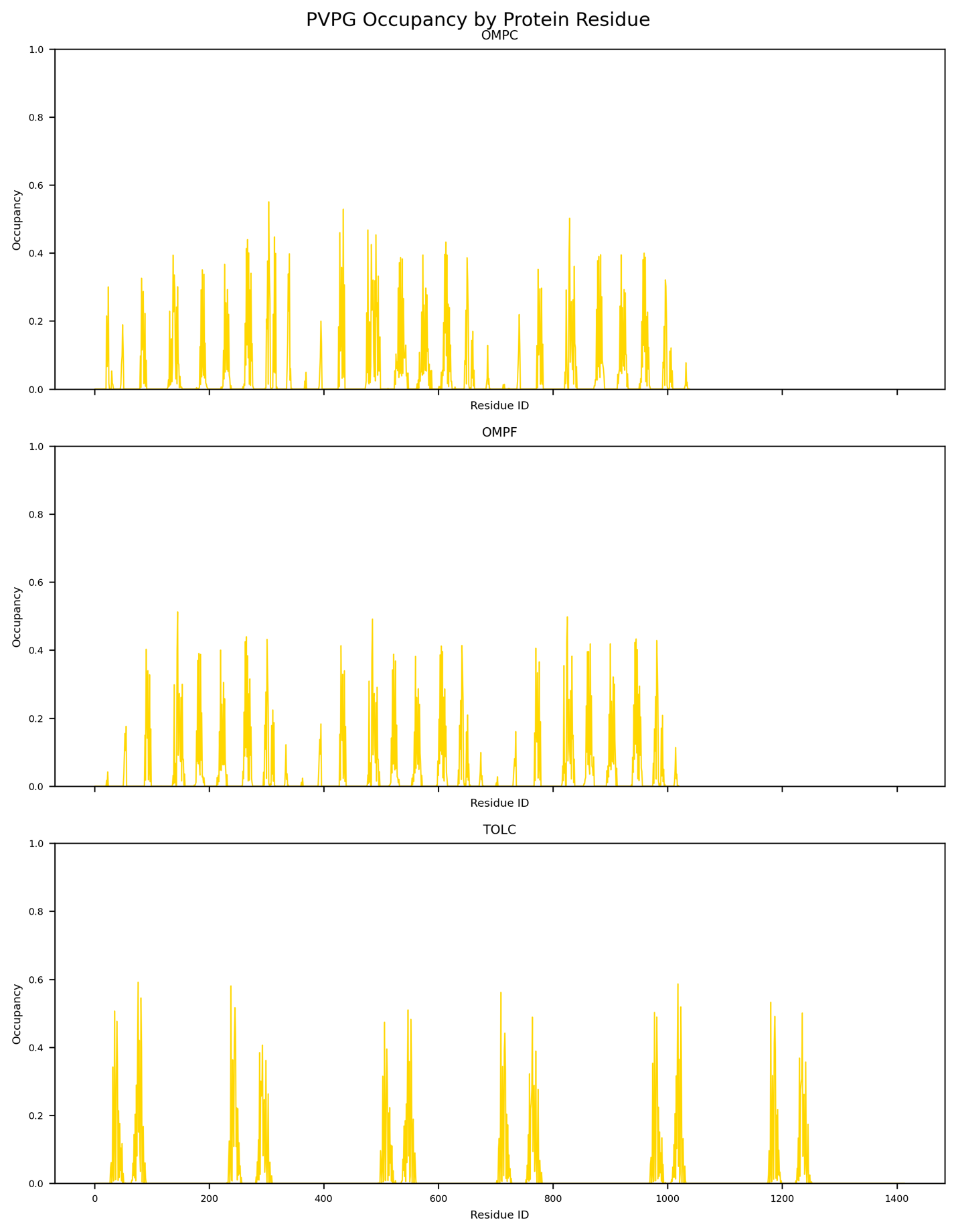

Figure S25: PVPG occupancy data for the 3 trimer protein systems with LPS O-antigen chain length 10.

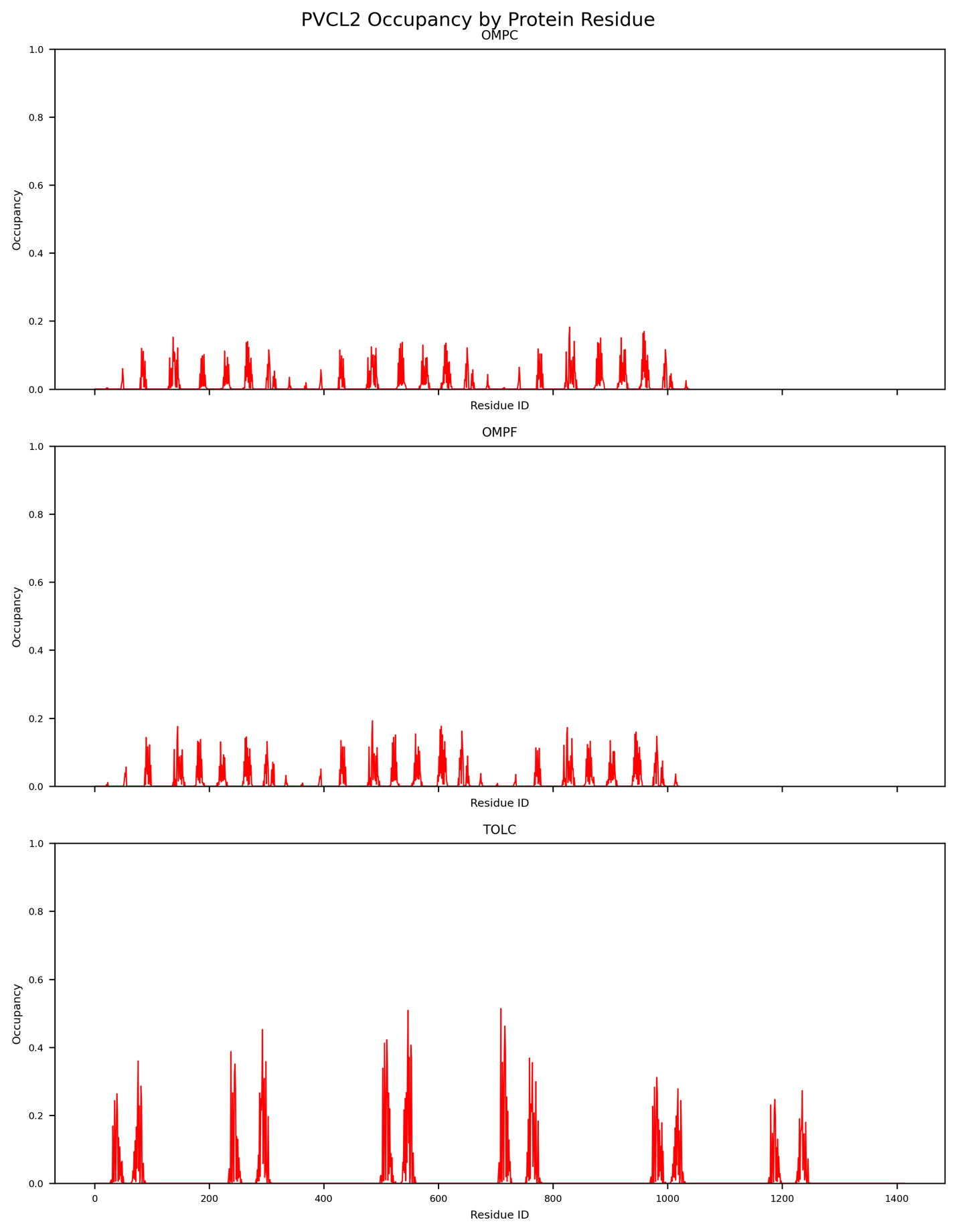

Figure S26: PVCL2 occupancy data for the 3 trimer protein systems with LPS O-antigen chain length 10.

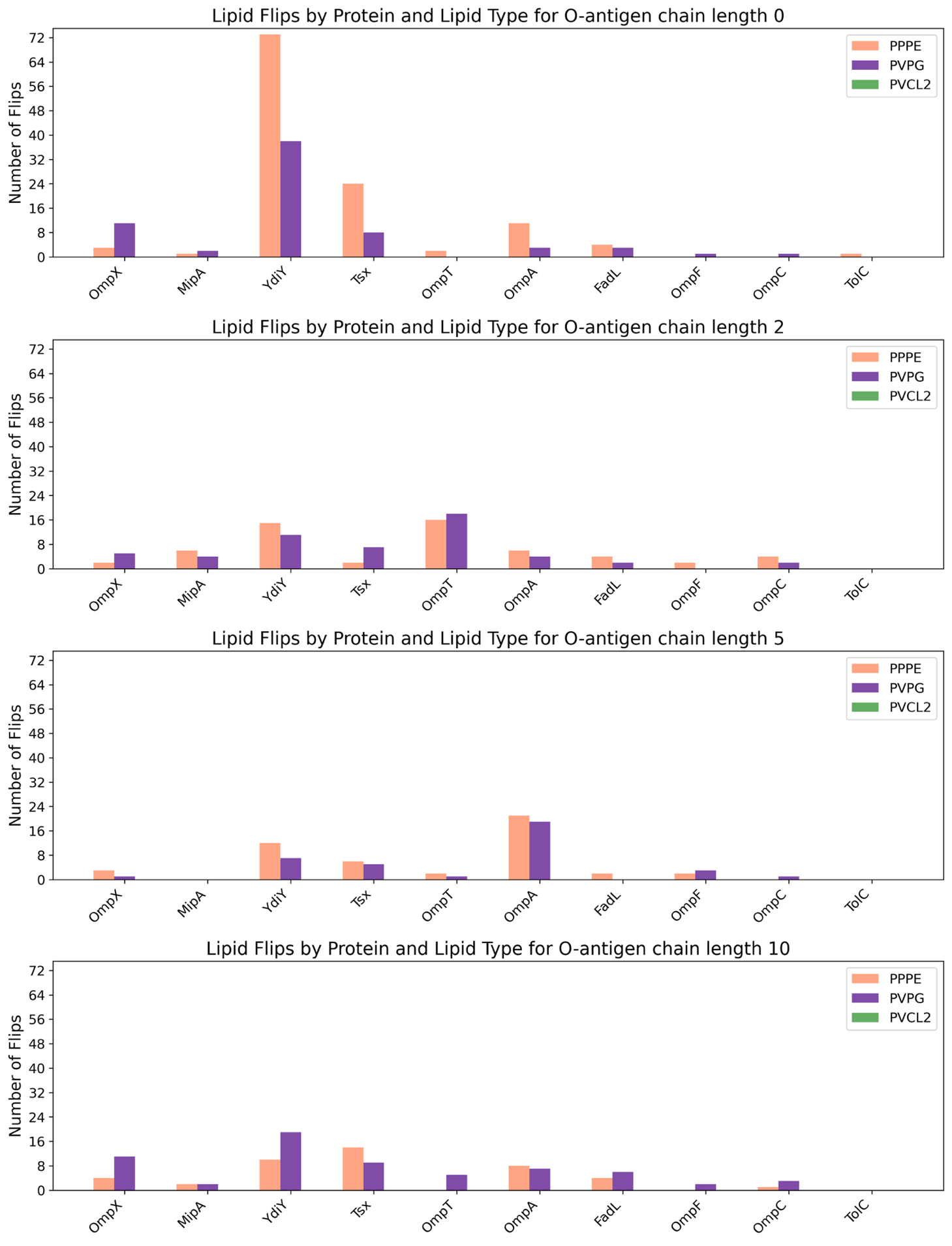

Figure S27: Lipid flips grouped by protein and colored according to lipid type. PVPG exhibited a disproportionately high fraction of flipping events relative to its abundance in the membrane.

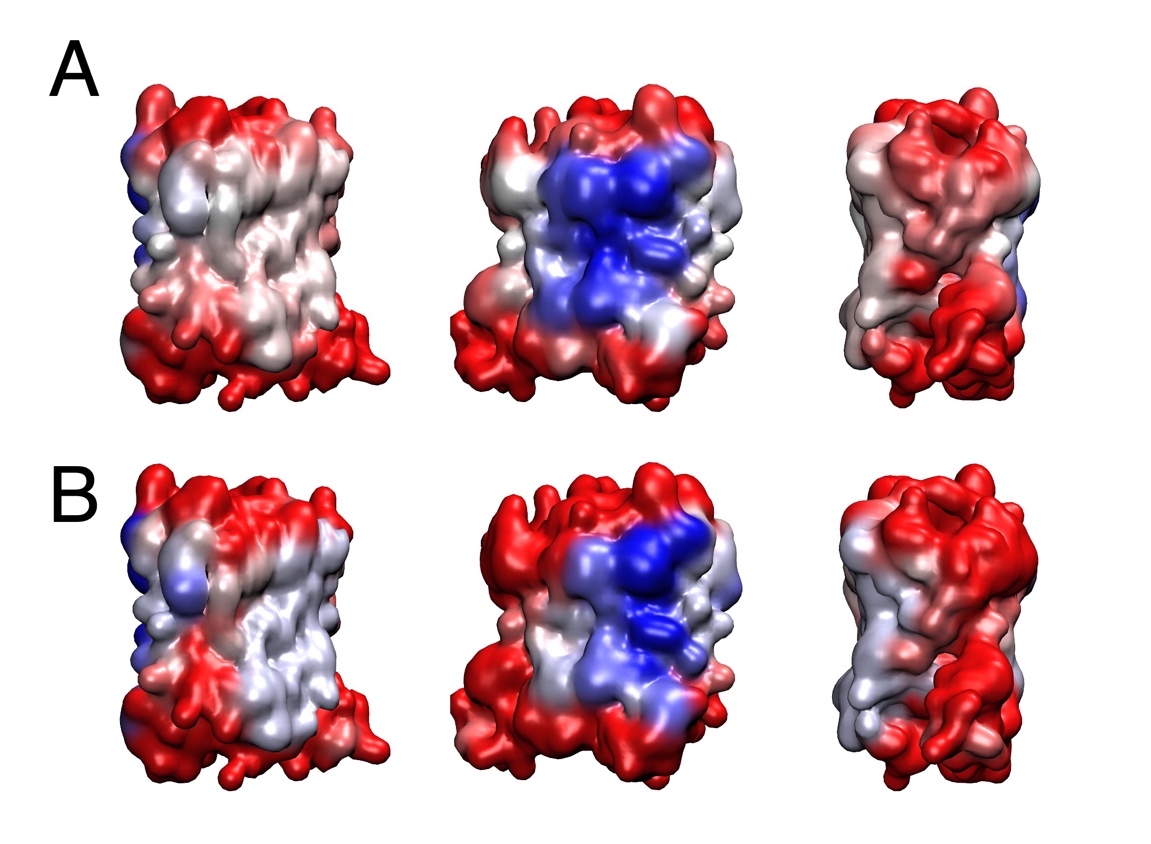

Figure S28: Analogous figure to Figure 6, but for O-antigen chain length two. Once again, **A** (top 3 figures) represents data from upward flips, whereas **B** represents data from downward flips.

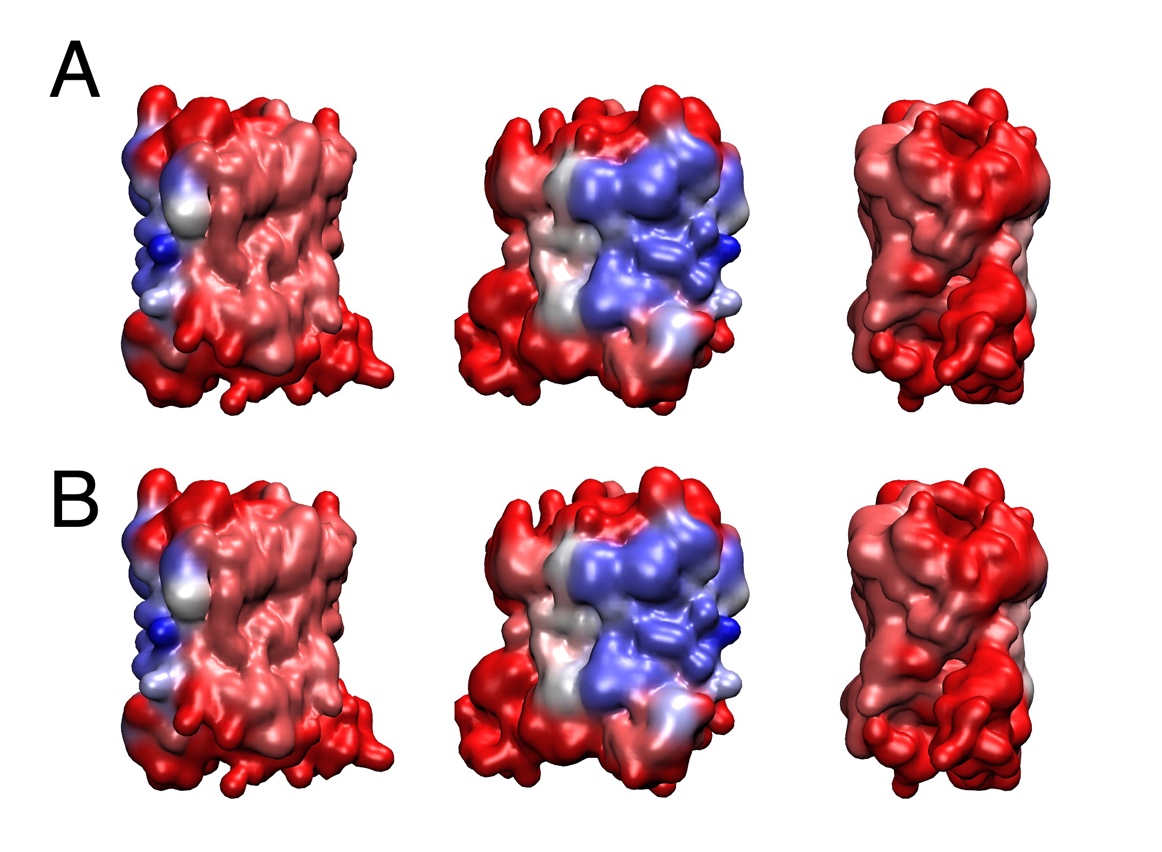

Figure S29: Analogous figure to Figure 6, but for O-antigen chain length 5. Once again, **A** (top 3 figures) represents data from upward flips, whereas **B** represents data from downward flips.

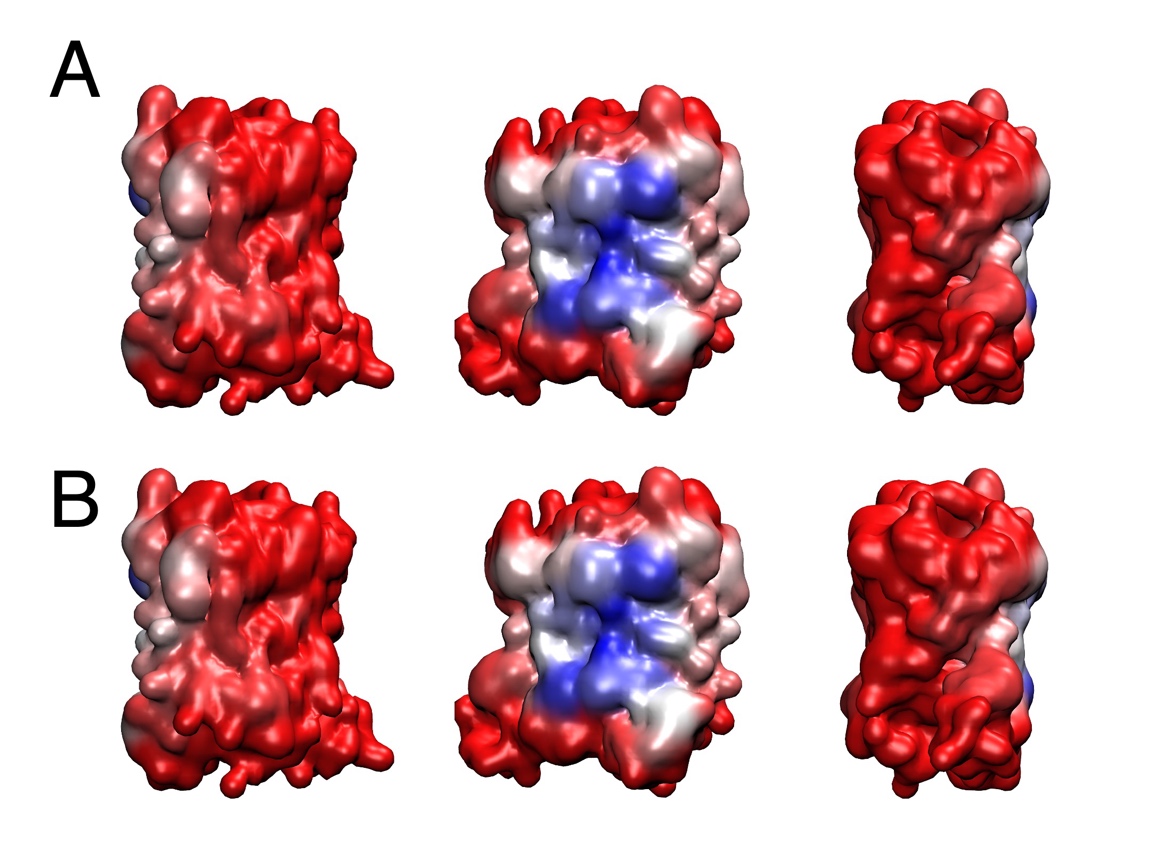

Figure S30: Analogous figure to Figure 6, but for O-antigen chain length 10. Once again, **A** (top 3 figures) represents data from upward flips, whereas **B** represents data from downward flips.

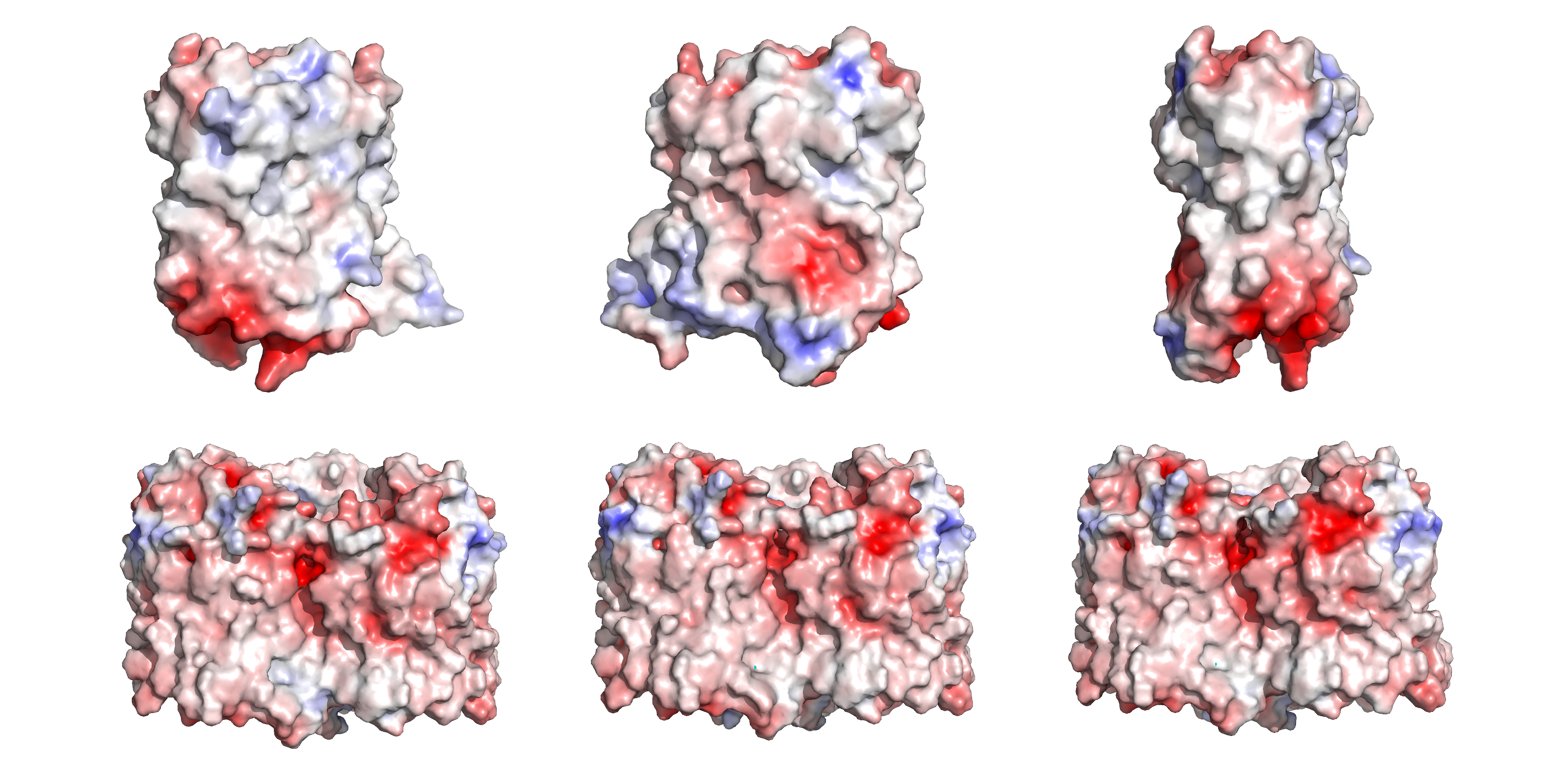

Figure S31: Adaptive Poissson-Boltzmann Solver results for YdiY (top 3 images) and OmpF (bottom 3 images). Three views of each protein are shown, rotated 120 degrees.

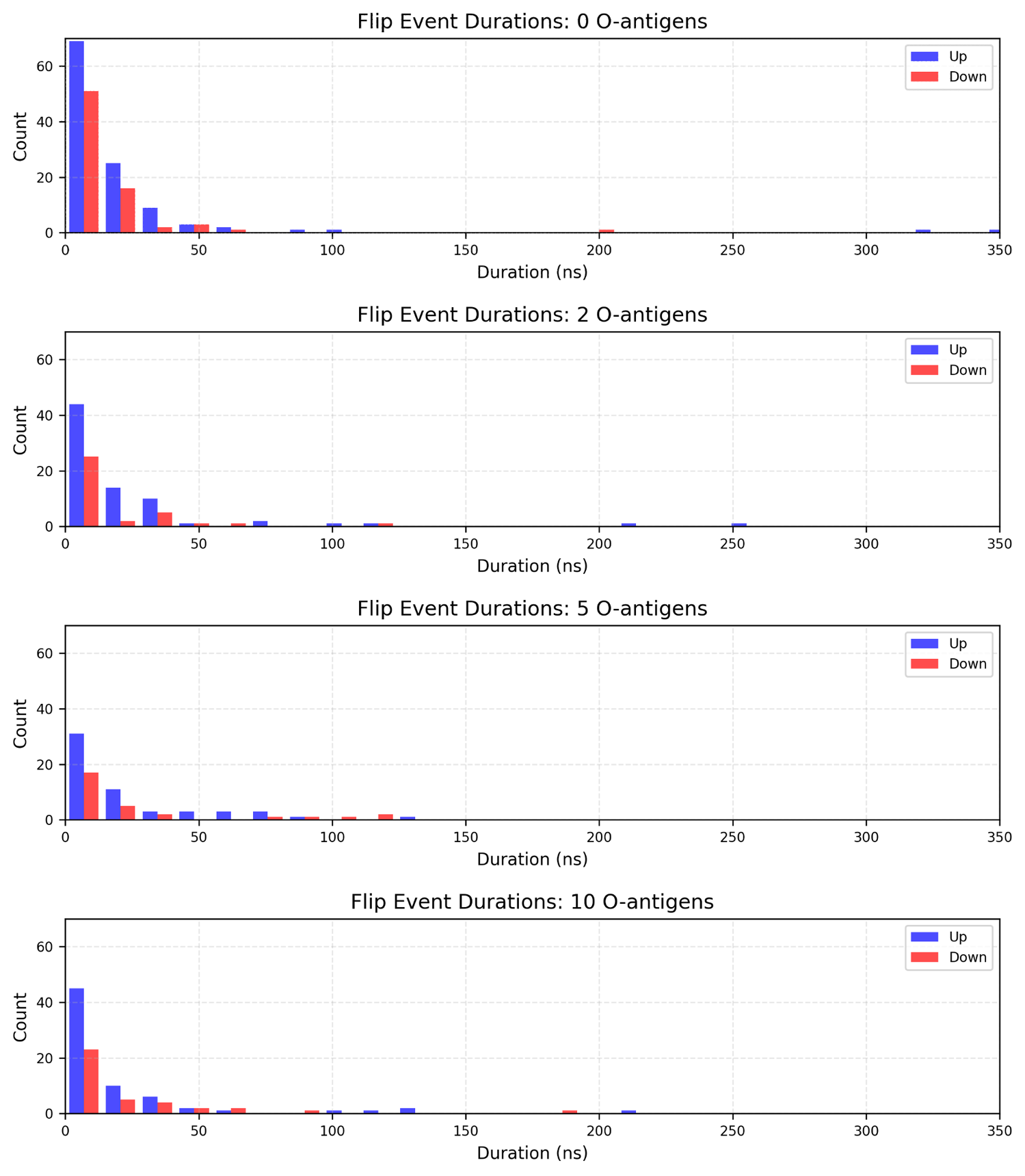

Figure S32: Flip event durations for my 4 sets of simulations, measured as the time between the point where the flipping lipid traversed 20% of the membrane region, to the time where it traversed 80%.

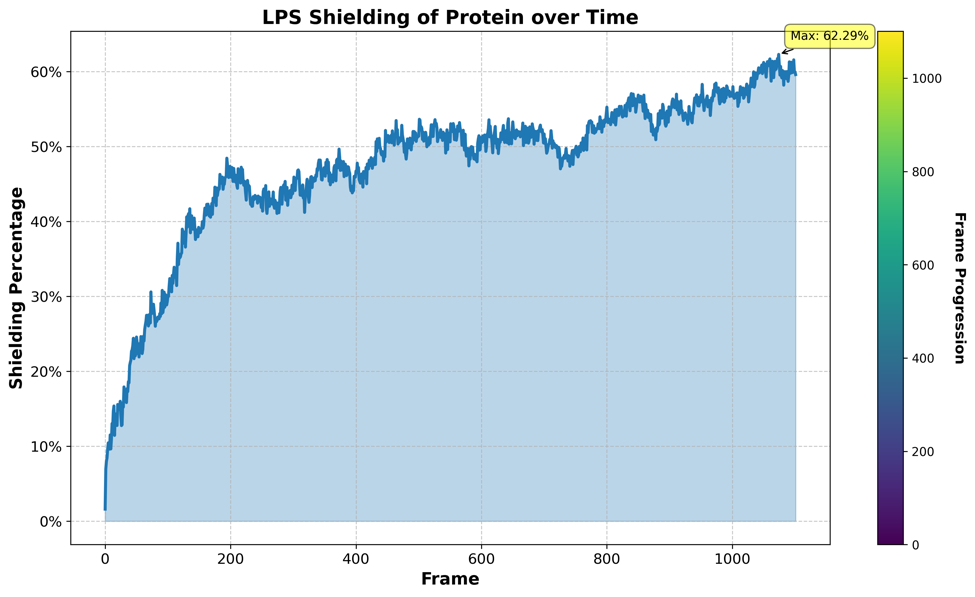

Figure S33: Time-course of percent-shielding for OmpC with 10 O-antigens during the 110 ns equilibration period for the simulation. Shielding increases rapidly during the first 20 ns, then increases more slowly throughout the final 90 ns.

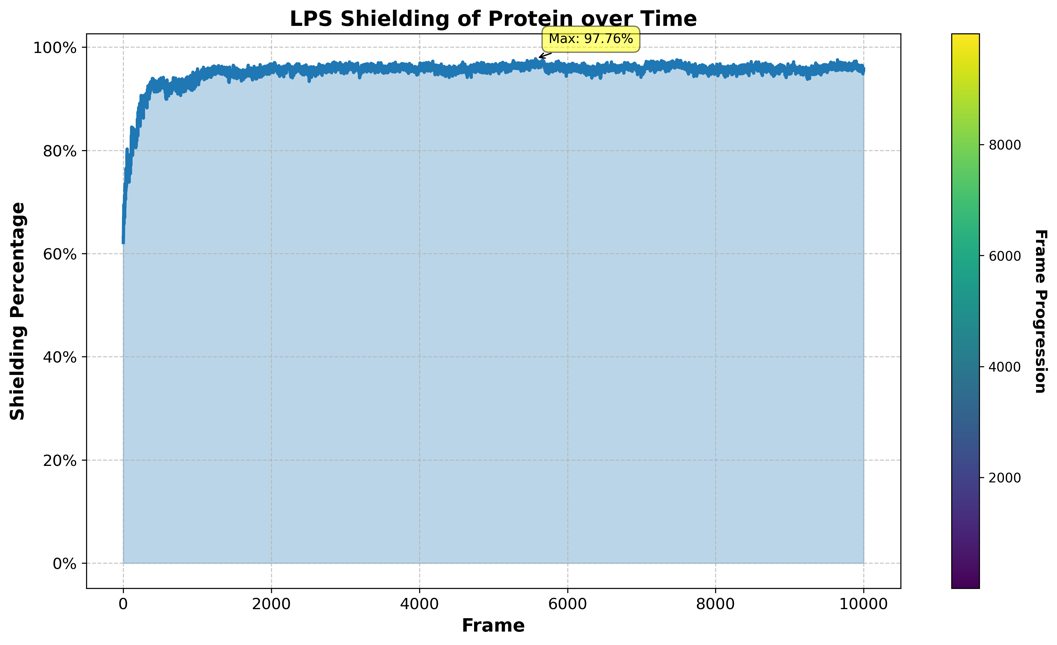

Figure S34: Shielding percentage for OmpC with LPS O-antigen length 10 throughout the entire 100 μs simulation, using every 100^th^ frame for the analysis

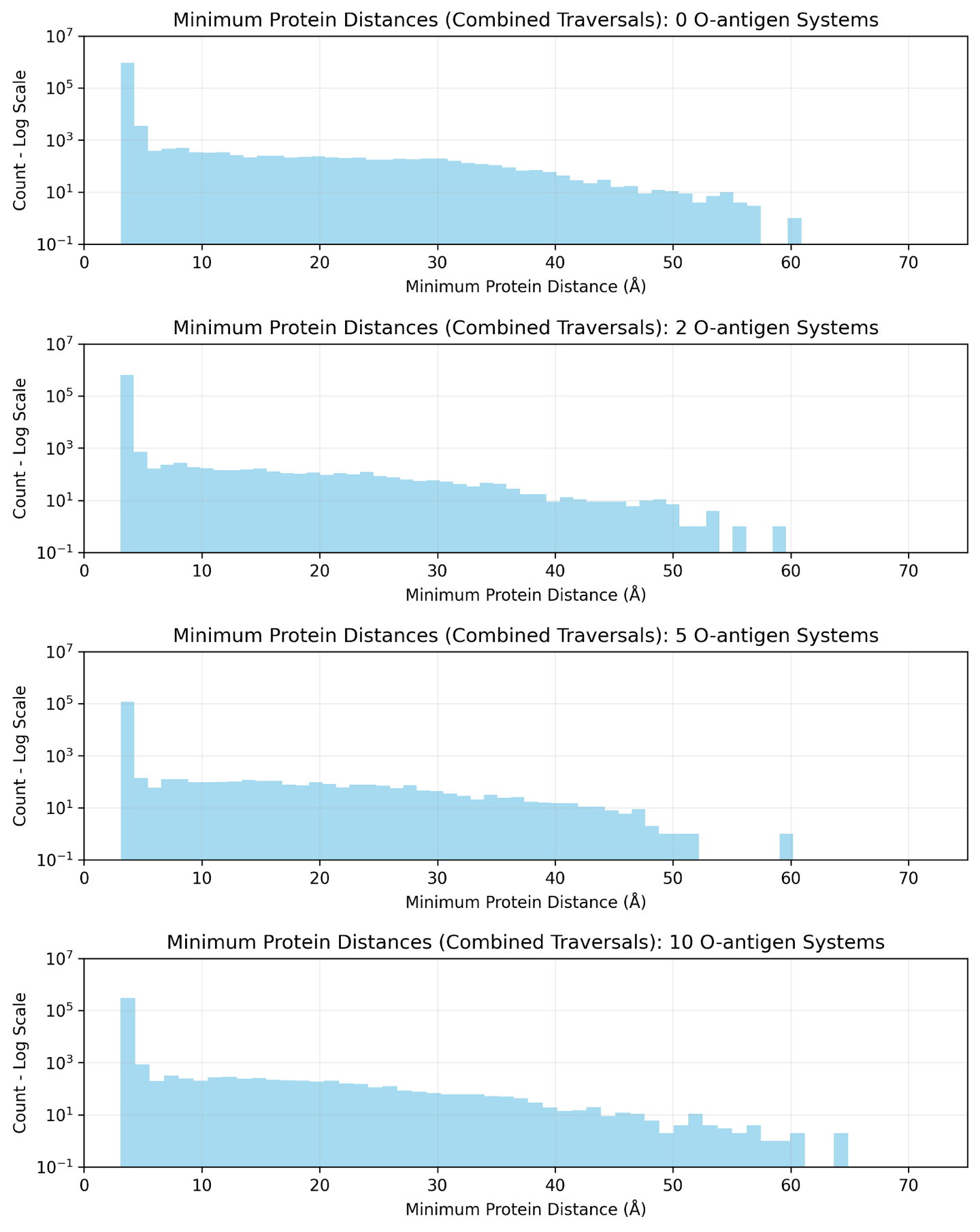

Figure S35: Minimum distance to the protein throughout each water traversal, with systems grouped according to O-antigen chain length

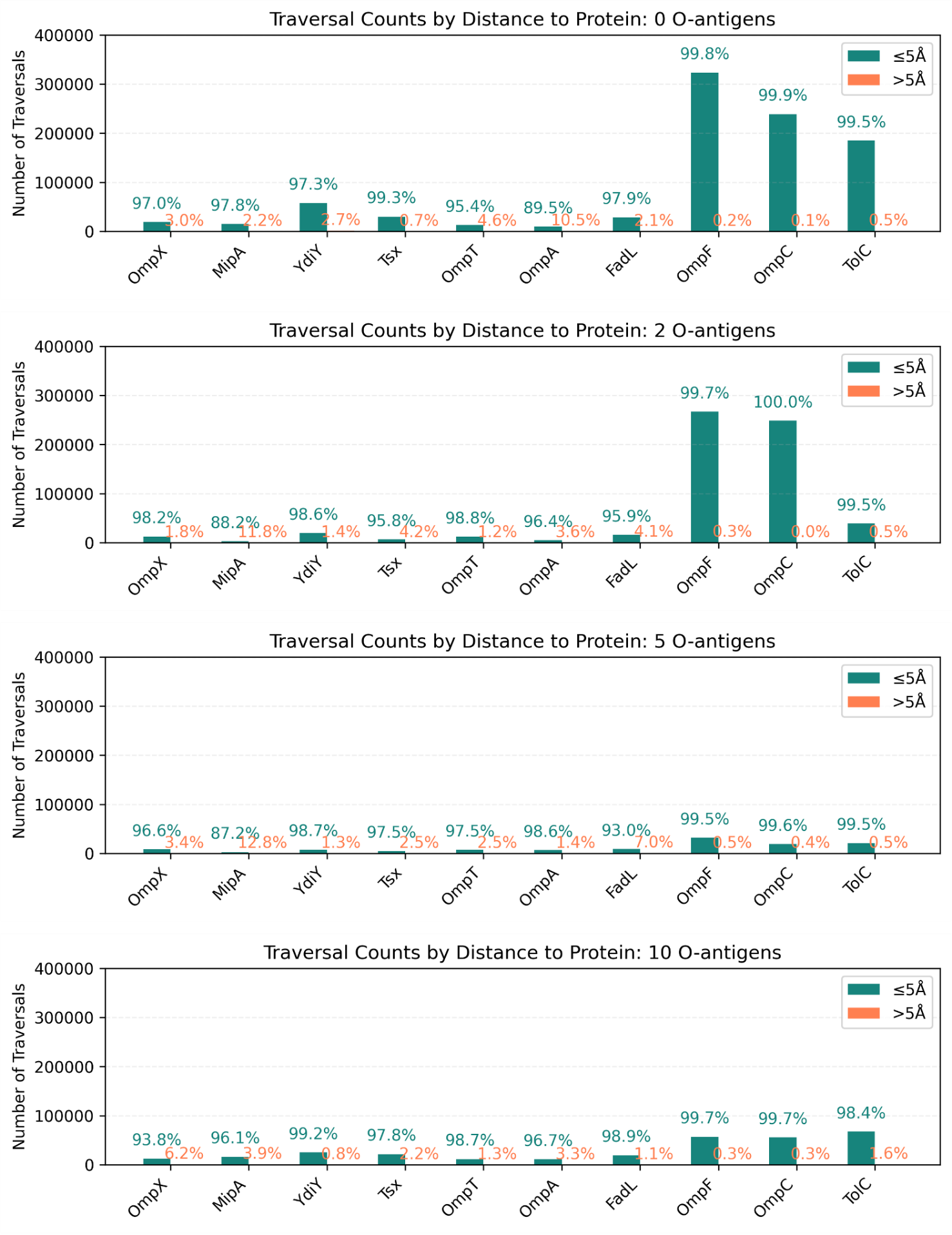

Figure S36: Water traversal counts by distance to the protein (greater or less than 5 Å). The vast majority of waters traversed the membrane within 5 Å of the protein.

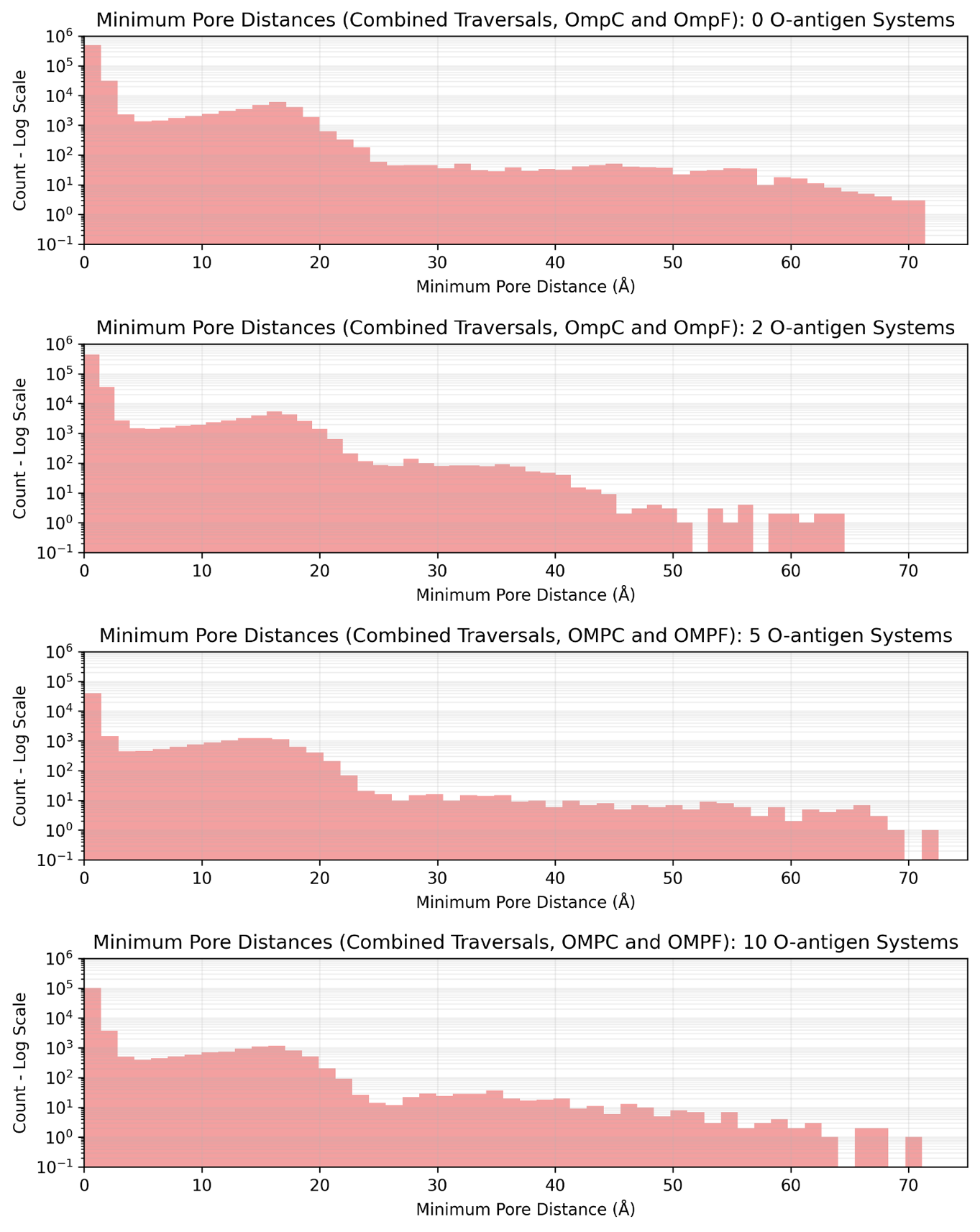

Figure S37: Minimum distance to the nearest pore for water traversals in the OmpC and OmpF simulations, grouped by O-antigen chain length

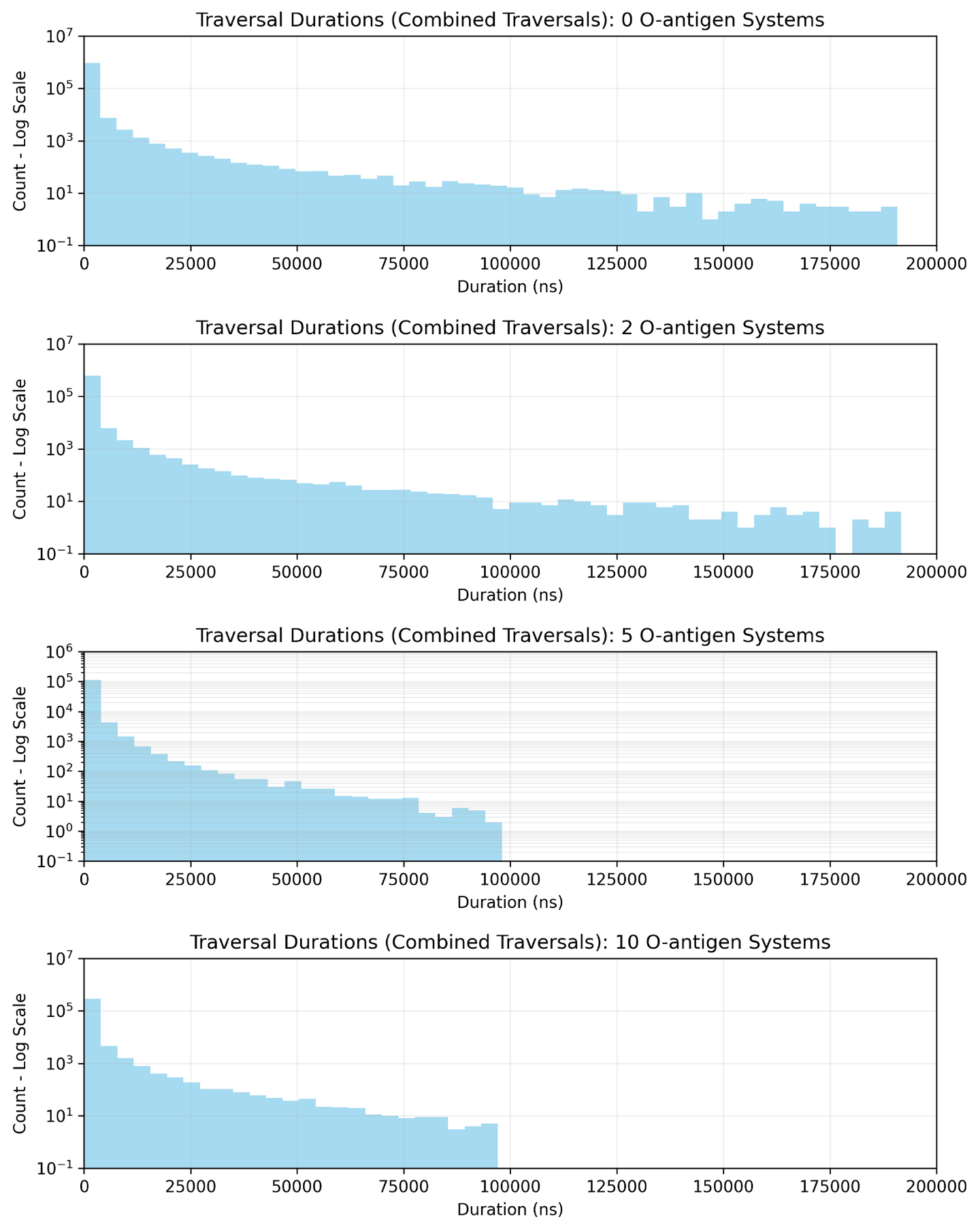

Figure S38: Duration of water traversals for all protein systems grouped by O-antigen chain length

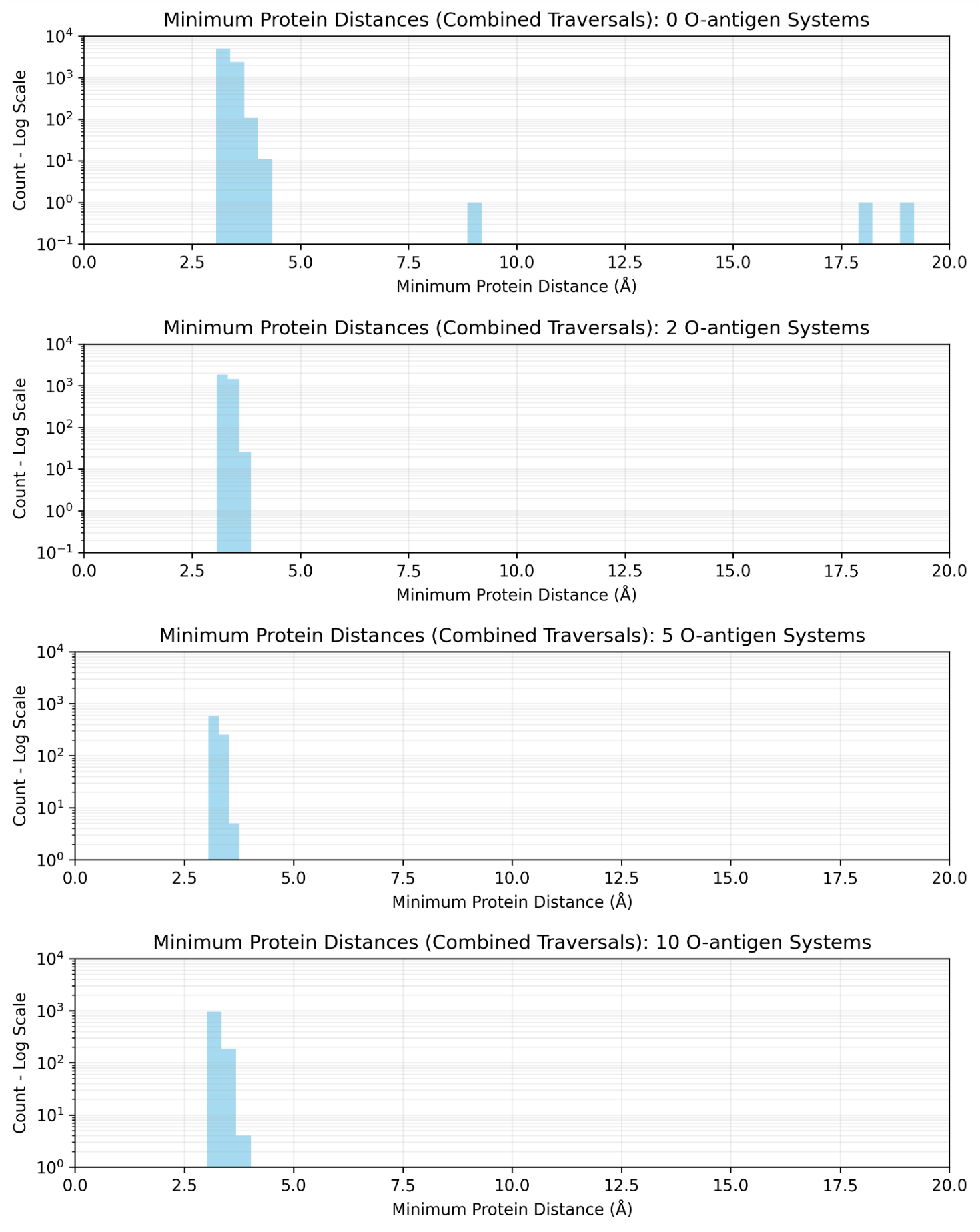

Figure S39: Minimum distance to protein for sodium ion membrane traversal events, with systems grouped by O-antigen chain length

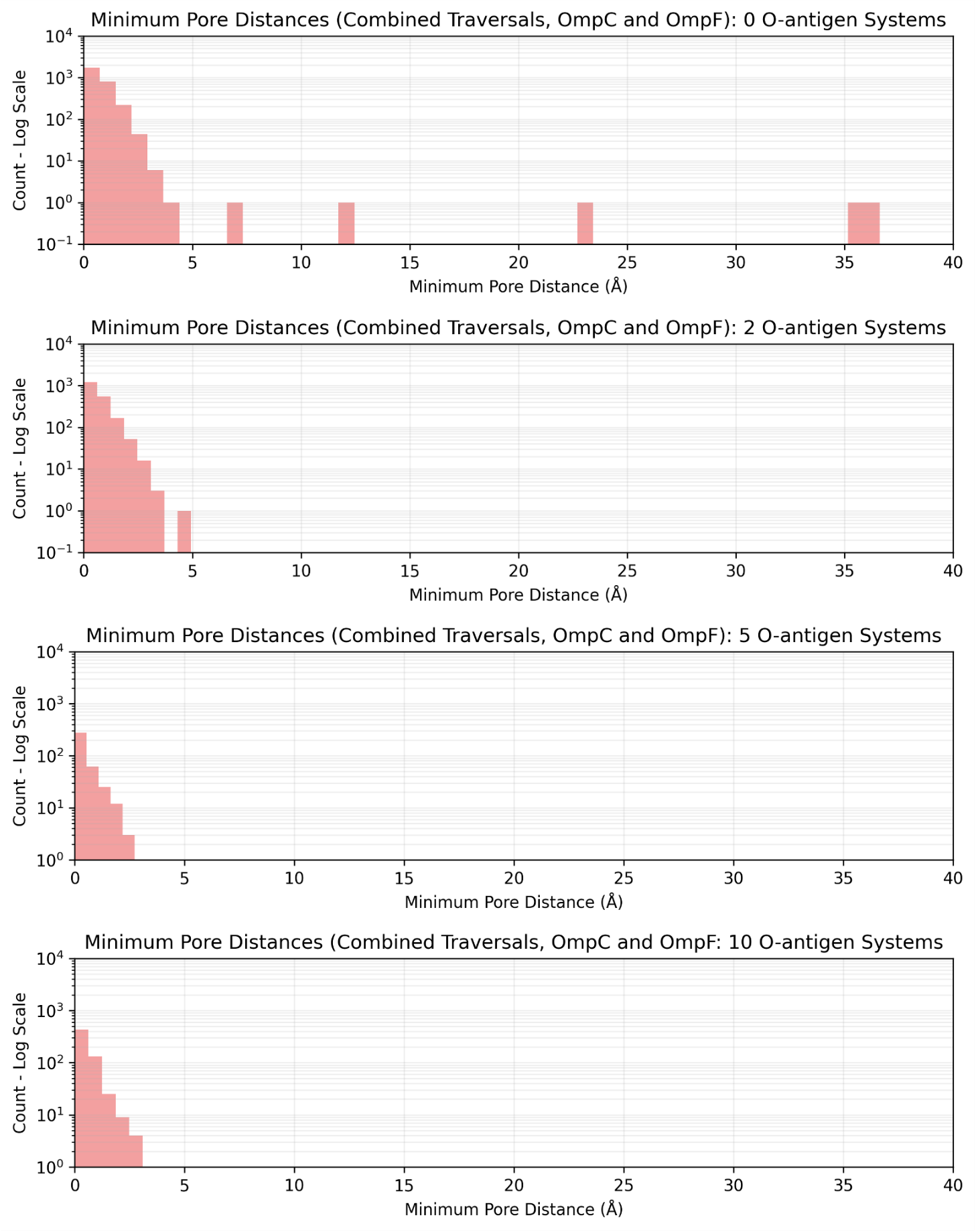

Figure S40: Minimum distance to a pore for sodium ion membrane traversal events, with systems grouped by O-antigen chain length

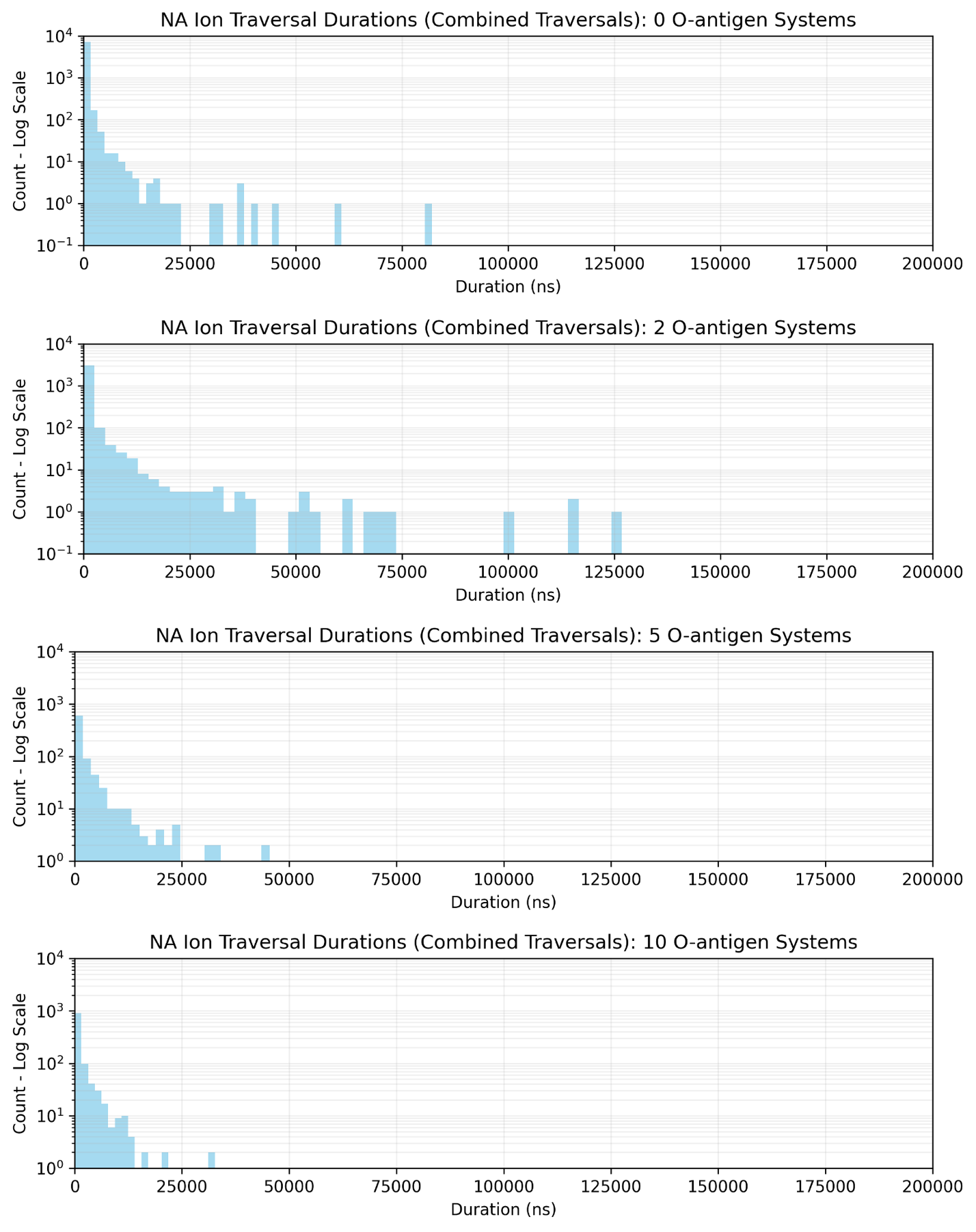

Figure S41: Duration of sodium ion traversal events, grouped according to O-antigen chain length

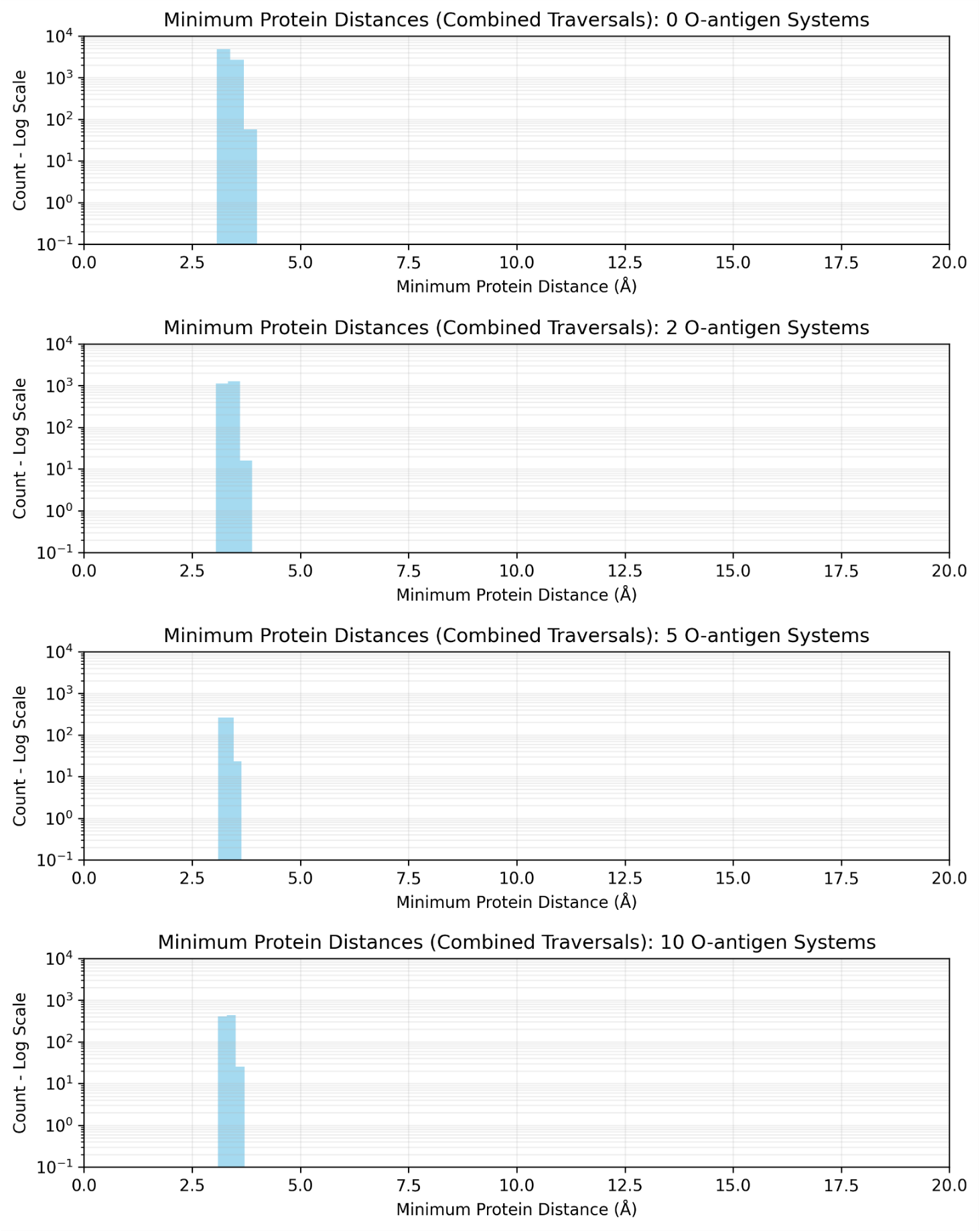

Figure S42: Minimum distance to the protein during chloride ion traversal events for the various protein systems, grouped according to O-antigen chain length

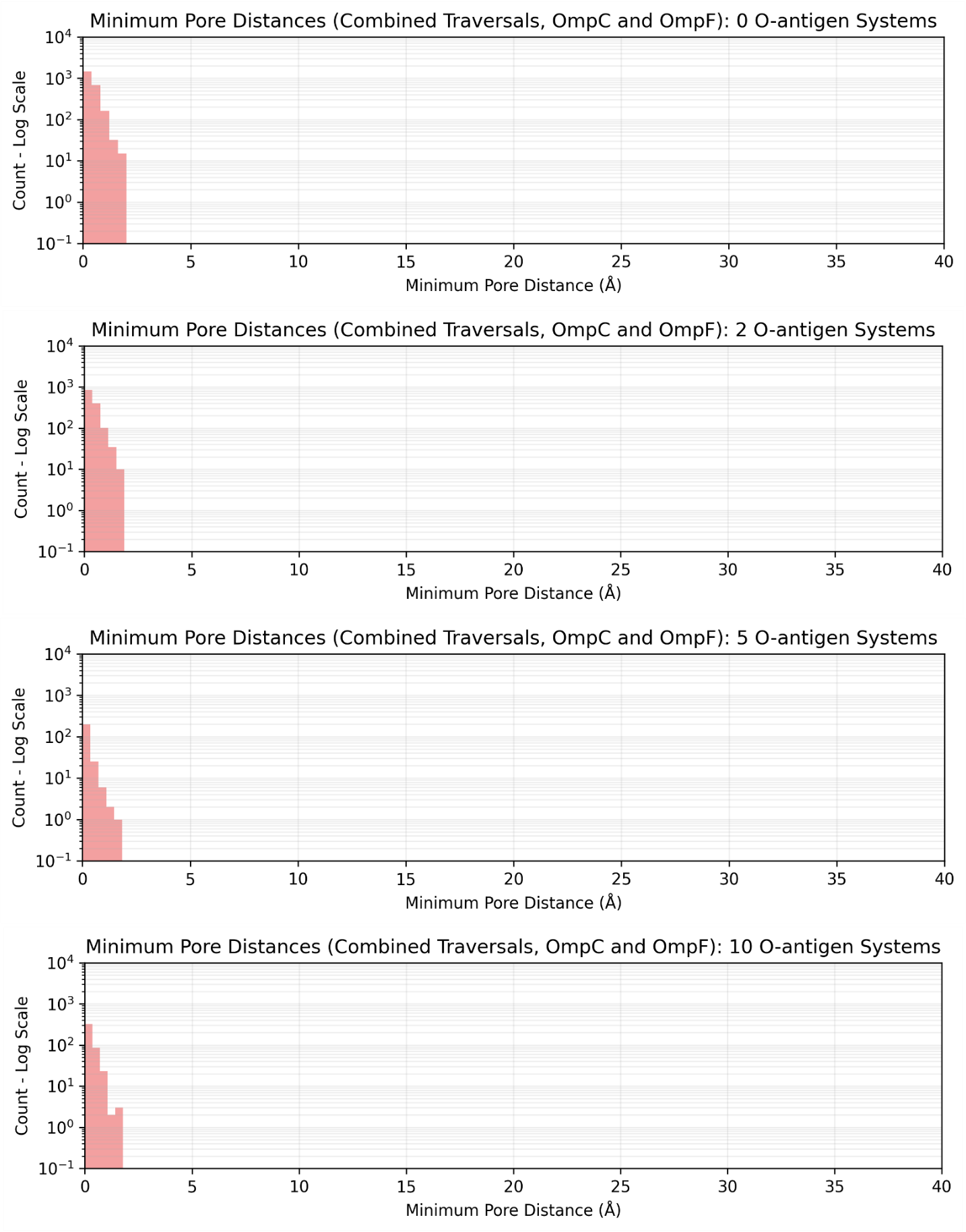

Figure S43: Minimum distance to a pore during a traversal event for OmpC, OmpF, and TolC, with systems grouped according to O-antigen chain length

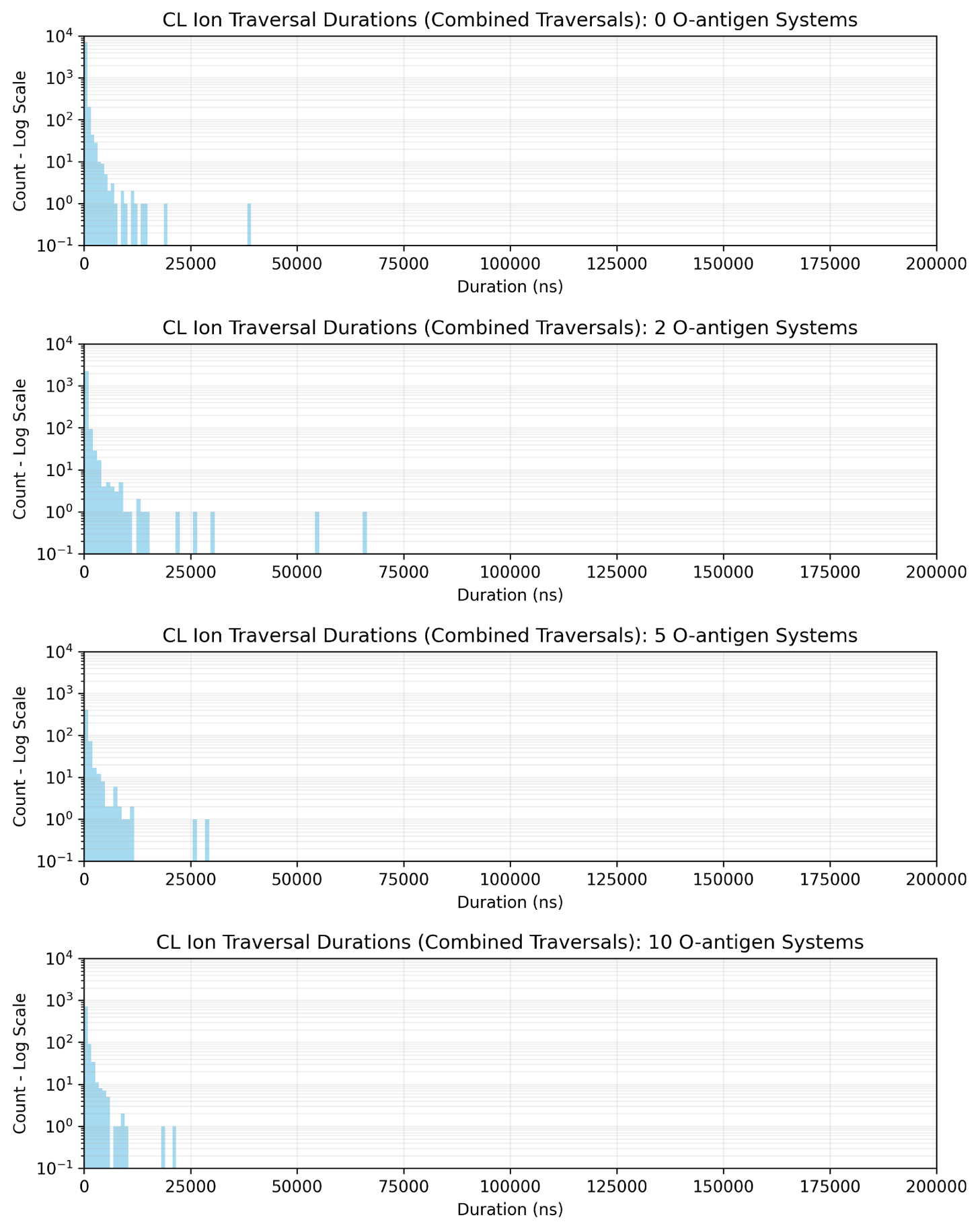

Figure S44: Chloride ion traversal event duration for the various protein systems, grouped according to O-antigen chain length
